# Computational method for mapping mass signatures along developmental gradients reveals a novel role for a monosaccharide tetrose in maize salt-stress response

**DOI:** 10.1101/2025.09.22.677919

**Authors:** Andrea M. Sama, Sinead B. Cahill, Shihong Luo, Abigail Tripka, Yifan Meng, Sarah E. Noll, Richard N. Zare, Pavak Shah, Alexandra Jazz Dickinson

## Abstract

Metabolic processes are essential for regulating and maintaining developmental transitions, from stem cell quiescence through differentiation. However, the distinct metabolite-driven mechanisms that are critical for development remain poorly characterized due to inherent challenges in measuring their production, localization, and function *in situ*. We employed desorption electrospray ionization mass spectrometry imaging (DESI-MSI) to map metabolites in the developing maize root, which has a well-characterized longitudinal gradient that encompasses developmental transitions from quiescence through proliferation and maturation. DESI-MSI enables *in situ* analysis of the chemical composition of tissue sections with high spatial resolution (∼50-100 µm). To identify metabolites with specific developmental enrichment patterns, we developed a computational approach called Developmental Imaging Mass Spectrometry Pipeline for Linear Evaluation (DIMPLE). DIMPLE processes mass signatures along linear gradients, generating clusters of metabolites with specific developmental enrichment patterns in the maize root. We employed this method to compare developmental enrichment of metabolites in a salt-resilient maize variety, Oaxacan Green, and a salt-susceptible variety, B73. DIMPLE identifies specific differences in the mass signatures and the overall enrichment patterns between these varieties. DIMPLE also revealed a metabolite, D-erythrose, that had different localization patterns in these varieties. We found that in salt-sensitive maize varieties, treatment with D-erythrose improves stress tolerance by increasing primary root length. Overall, DIMPLE enables comprehensive and rapid analysis of metabolite patterns along a linear gradient, revealing new biology in plant growth and stress response.

## Introduction

Distinct metabolic processes are critical for developmental transitions between cell stages such as stem cell quiescence, proliferation, and differentiation. Each developmental transition requires specific cellular energetic states as well as metabolite-driven signaling and protein interactions. In plants, metabolites play particularly important roles as regulators that bridge development and environmental responses (Erb & Kliebenstein, 2020). For example, a classic control point regulating both development and stress responses in plants is the phytohormone abscisic acid, which is activated by abiotic stress and regulates seed germination and root growth (Hauser et al., 2017; Nakashima & Yamaguchi-Shinozaki, 2013; Tuteja, 2007). However, despite the importance of metabolites in development and stress responses, metabolite-based mechanisms remain poorly characterized due to inherent challenges in measuring their production, localization, and function *in situ*. These problems are amplified when considering metabolites with precise spatial enrichment patterns, such as those that occur during developmental transitions. Metabolomics approaches typically require relatively large amounts of tissue (mg – g scale), masking the metabolic signatures of rare cell types, such as stem cells (Jackson & Finley, 2024). Current approaches to study metabolites in rare cell types often require physically isolating these cells from tissue (Binek et al., 2019; Jackson & Finley, 2024). However, these approaches can influence the cell metabolome (Binek et al., 2019), which can change on a timescale of seconds to minutes. In addition, isolation of cells removes microenvironment cues (Jackson & Finley, 2024), which may be critical, particularly during development. Additionally, bulk tissue approaches do not capture the heterogeneity of developing cells, which may be in a variety of transitional states in terms of differentiation, quiescence, and stress response (Miyazawa & Aulehla, 2018).

Mass spectrometry imaging (MSI) offers an alternative method of approaching metabolomic investigations in developing tissue (Gao et al., 2023). There are many approaches to MSI such as desorption electrospray ionization (DESI), matrix assisted laser desorption ionization (MALDI) and secondary ion mass spectrometry (SIMS). These different techniques have their own advantages and limitations, and have been thoroughly reviewed (Buchberger et al., 2018; Gao et al., 2023). MSI provides *in situ* analysis of the chemical makeup of tissue sections with high spatial resolution ranging from ∼5-100 µm depending on the ionization method. This can enable near single cell visualization of metabolomic changes throughout tissue, which is advantageous for studying rare cell types (Lanni et al., 2012; Römpp & Spengler, 2013). Furthermore, imaging cells in their native microenvironment maintains the structure of tissue and the cues that regulate developmental transitions. Therefore, MSI is a valuable tool for studying the metabolism of stem cell decisions as well as understanding how metabolite enrichment changes throughout tissue development.

To investigate metabolite enrichment patterns during development, we utilized DESI-MSI to map the chemistry of the developing maize root (T. Zhang et al., 2023). DESI-MSI uses a charged solvent spray to desorb compounds from the surface of the tissue. The desorbed analytes are then ionized in secondary droplets that are detected by the mass spectrometer. Moving the desorption spot along the tissue’s surface generates a rasterized image, where every pixel contains a full mass spectrum. By collecting full mass spectra, the varying intensities of individual molecular mass signatures can be mapped throughout the tissue, providing spatial information for a range of metabolites, lipids, and peptides. This information can be used to predict the identity and function of endogenous chemical compounds.

In a previous study, we applied this technique to longitudinal cryosections of maize roots at a spatial resolution of approximately 80 µm (T. Zhang et al., 2023). Plant roots represent a highly stereotyped developmental gradient. At the root tip is the meristem region, which encompasses the stem cell niche (SCN) and dividing pluripotent cells. Progressing along the root axis, cells lose their ability to divide and begin to lengthen in the elongation zone (EZ). These developmental transitions are encompassed in a ∼5 mm longitudinal section of the maize root tip (Alarcón et al., 2014). Cells finally transition to full maturity in the differentiation zone (DZ) (Hochholdinger, 2009; Marcon et al., 2015). Our previous work applying DESI-MSI to the maize root tip identified 63 ions with specific localization patterns along this developmental axis (T. Zhang et al., 2023). One finding from this work was that the tricarboxylic acid (TCA) cycle metabolites have distinct localization patterns across the developmental gradient of the root. For instance, of all the TCA cycle metabolites we measured, succinate was the only one enriched in the meristem region of maize roots. Further study of this specificity revealed that exogenous succinate treatment increased meristem cell divisions in Arabidopsis roots, demonstrating that localization patterns can inform predictions of developmentally relevant metabolite functions (T. Zhang et al., 2023).

Although MSI has enormous potential as a tool to characterize metabolites in developing tissue, there are challenges to analyzing and interpreting MSI datasets, particularly in identifying new metabolites with strong enrichment to specific developmental transitions. MSI datasets are large; each pixel contains a full mass spectrum with hundreds to thousands of peaks. This sheer volume of data presents a challenge, especially when conducting an untargeted analysis of chemical distributions throughout tissues. Several commercial and open-source software options have been developed to aid in the processing and analysis of MSI data (Bemis et al., 2015; Bokhart et al., 2018; Hu & Laskin, 2022; Källback et al., 2016; Robichaud et al., 2013). A common software, previously open-source and now commercial, used for MSI data analysis is MSiReader (Bokhart et al., 2018; Robichaud et al., 2013). MSiReader v1.03 allows users to optimize visualization of *m/z* features by adjusting parameters such as normalization method, image overlay, and co-localization of metabolite signals. MSiReader v1.03 also provides several tools to aid in the analysis of untargeted metabolomics such as MSi Correlation and MSi Peakfinder strategies. However, these methods rely on user defined regions of interest or peak selections and the processing times for these analyses can be significant. MSiReader v1.03 includes a plug-in to METASPACE that enables database searching to annotate unknown mass signatures. METASPACE is an open-source cloud platform for spatial data interpretation and metabolite identification (Palmer et al., 2017). It performs automated, false discovery rate (FDR)-controlled metabolite identification using an array of user-selected databases, such as the Human Metabolome Database (HMDB) and LipidMaps, creates visualization profiles of specific annotations, and compares localization patterns across tissues. Other mass spectrometry resources such as Metlin Classic and LOTUS can also be useful in annotating mass signatures. Metlin Classic is a metabolite search engine that provides hits for *m/z* values based on ionization mode and defined mass error; it also includes fragmentation patterns for additional confirmation (Guijas et al., 2018; Smith et al., 2005). LOTUS is an open-source tool for natural products identification and is particularly helpful for annotating plant metabolites (Rutz et al., 2022). These resources were all valuable in identifying molecular patterns of interest and/or predicting the chemical identities of *m/z* signatures in our datasets.

We applied many of these resources to analyze our maize root DESI-MSI data and found dozens of mass signatures with distinct patterns during development (T. Zhang et al. 2023). However, these identifications required a labor-intensive manual interpretation of the raw data and sorting of developmental patterns, in addition to the utilization of open-source software. This work revealed the need for a new computational method to identify mass signatures along the linear root developmental gradient. To address this gap and streamline untargeted analysis, we developed a tool specifically for interpretation of plant root MSI data. MSI analytical tools typically include four main steps, 1) pre-processing, 2) peak picking, 3) de-noising and 4) spatial clustering (Alexandrov et al., 2010, 2013; Alexandrov & Kobarg, 2011; Ovchinnikova et al., 2020). There are several approaches that utilize variations of these steps to identify peaks and cluster them based on measures of 2D spatial similarity (Alexandrov et al., 2010; Y. Zhang et al., 2023). While unbiased spatial clustering is broadly applicable to many types of tissue, we sought to reveal patterns based on a specific spatial prediction – that the linear enrichment of a molecule along the developmental gradient would reveal useful biological information. Therefore, we created a computational method called Developmental Imaging Mass Spectrometry Pipeline for Linear Evaluation (DIMPLE), which is designed to leverage the linear developmental axis of plant roots to reveal distinct metabolite patterns that correspond to the transitions between quiescence, proliferation, elongation, and differentiation. We validated its performance and utilized DIMPLE to compare developmental patterns in a stress-tolerant maize variety compared to a stress-susceptible variety. DIMPLE led to the detection of a meristem and EZ localized metabolite, D-erythrose. We further found that D-erythrose has different localization patterns in the drought and salt tolerant maize variety we characterized. In salt-sensitive varieties, treatment with D-erythrose improves root growth during stress.

Overall, DIMPLE contributes to the growing number of tools used for MSI analysis. It provides a simple and effective approach to cluster mass signatures along linear developmental gradients, allowing for characterization of root metabolite patterns. Application of this method to better understand the relationship between environmental stress and development also demonstrates the power of MSI to inform biological hypotheses regarding the significance of metabolite localization patterns.

## Results and Discussion

### Development and implementation of DIMPLE

The DIMPLE workflow consists of two core steps. First, raw DESI-MSI data is pre-processed to identify a set of consensus peaks and their corresponding *m/z* values across all pixels; then, observed peaks across all acquired spectra are assigned to the *m/z* values identified in the consensus set. Second, these harmonized spectra are converted to a hyperspectral image that can be analyzed using conventional image analysis and signal processing approaches to identify linear patterns of enrichment along the primary tissue axis. Pre-processing consists of the following steps: *1)* Optional hot pixel detection and suppression, *2)* Manual segmentation of the sample boundary from background, *3)* Generation of a pseudobulk spectrum by summing all peaks detected in every pixel, *4)* Identification of consensus peaks in the pseudobulk spectrum by identifying local maxima within a ± 2.5 ppm window, *5)* Suppression of peaks that are not enriched in the sample over background, *6)* Assignment of every observed sample-enriched peak in each pixel to the peak centers identified in the consensus spectrum, *7)* Reorganization of the harmonized spectra into a hyperspectral image, and *8)* Removal of peaks that show sparse localization. The removed peaks are retained separately and can be manually inspected for any potential patterns that were erroneously removed. This workflow is meant to ensure that all observed peaks in the sample are aligned to each other independent of instrument jitter within a reasonable error window.

Following this preprocessing, we devised a simple strategy for the unbiased discovery of spatial motifs in metabolite enrichment in the maize root. We focus specifically on the analysis of patterns along the longitudinal (developmental) axis of the root, as the development and growth of the root along this axis is well-characterized (Figure 1A). Adopting a standardized orientation of the longitudinal axis of the root along the y-axis of the image, we compute a maximum projection of the 2D hyperspectral image generated by the preprocessing step along the x-axis reducing each image to a 1D longitudinal linescan (Figure 1B). Each linescan is normalized such that its maximum value is 1 and smoothed with a 10-pixel wide median filter. We then perform hierarchical clustering of these linescans, each corresponding to an *m/z* value, using the Euclidean norm between linescans as the distance metric (Figure 1C). Common limitations to hierarchical clustering with MSI data is the amount of computational memory and power required to process the linkages between spectral patterns (Alexandrov & Kobarg, 2011; Deininger et al., 2008). Since our method reduces the spectral patterns to a single pixel-wide (one dimensional) linescan across a consensus set of high-quality channels, data volume is reduced drastically resulting in a less intensive processing for hierarchical clustering. By using Ward’s linkage method, each spectral linescan is first considered its own cluster, and are sequentially merged using a bottom-up approach where patterns are merged to preference the smallest within cluster variances. For each dataset, we determined an appropriate number of clusters that encompass general developmental patterns by examining the structure of the resulting dendrograms (Figure 1C, Supplemental Figure 1) and by examining the distributions of localization patterns among clusters (Figure 1D). We implement a simple GUI in MATLAB that allows the user to browse the clustered linescan data and extract the original 2D image for any corresponding channel for export (Figure 1E).

**Figure 1.**
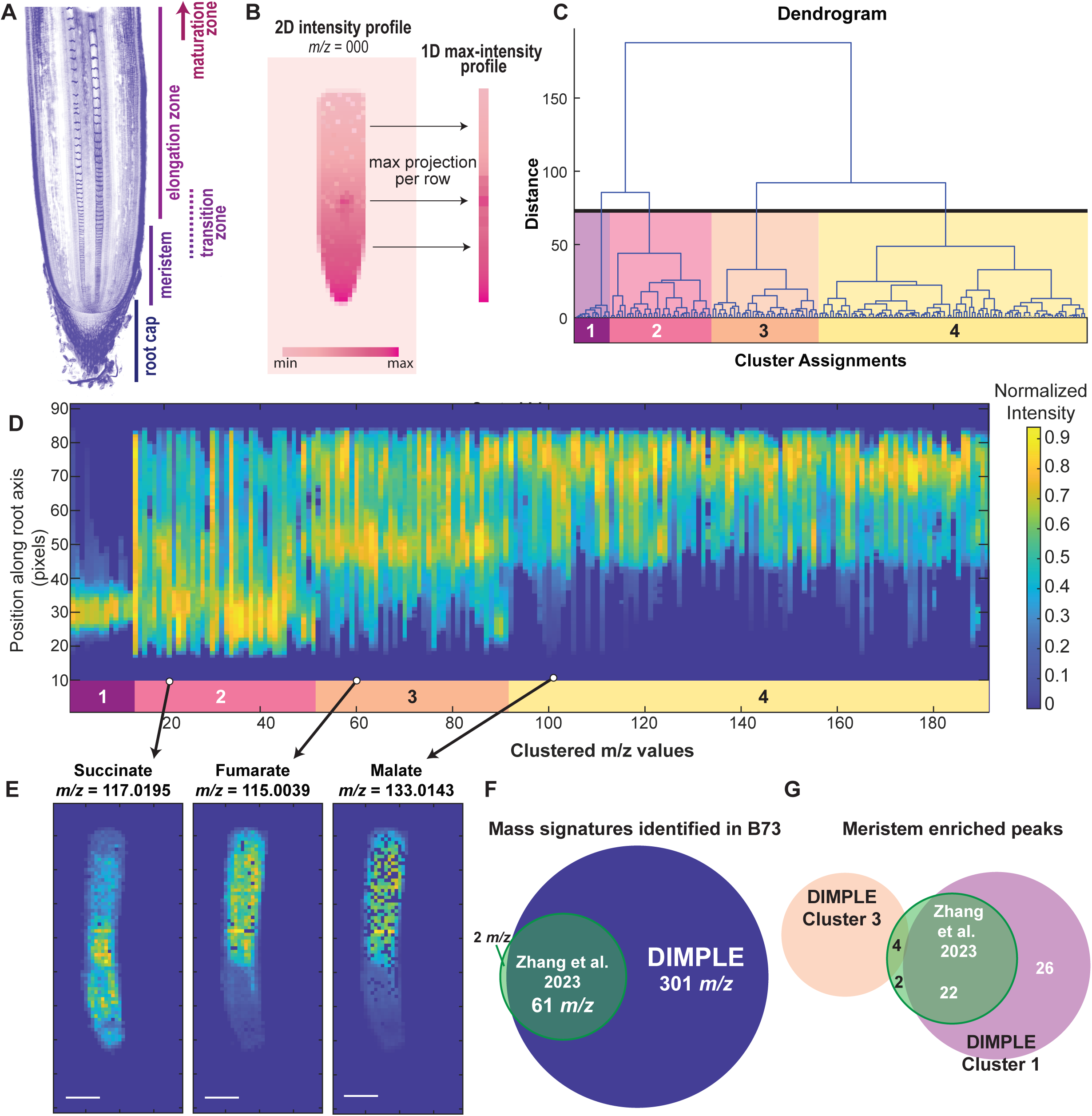
DIMPLE’s clustering method increases mass signature identification and annotation. **A)** Schematic of maize developmental root axis, characterized by the root cap, meristem, transition zone, elongation zone and maturation zone. Confocal image of a maize root section stained with calcofluor white. **B)** Schematic of DIMPLE’s approach, which reduces the 2D intensity profile of the MSI data into a 1D linescan profile, based on the maximum projection per each row of the 2D image. The resulting heatmap is normalized to the maximum intensity. **C)** The linescans are processed using Ward’s hierarchical clustering method resulting in four distinct groups that relate to the spatial similarity of linescan intensity profiles. User-defined number of clusters can then be visualized as shown in D. **D)** Linescan graph showing the intensity profiles of the mass signatures in each cluster. Clusters are designated by the colored bars arranged left to right and labelled with corresponding cluster number. The y-axis is the position along the root axis in pixels where 0 is closest to the meristem and 90 is the mature tissue of the root. The x-axis corresponds to the clustered *m/z* linescans arranged smallest to largest within each cluster. The intensity of each linescan is normalized to a maximum value of 1, which is represented in the “Normalized Intensity” color bar legend. **E)** 2D spatial maps of the signal enrichment for succinate (*m/z* 117.0195), fumarate (*m/z* 115.0038) and malate (*m/z* 133.0143) in a B73 maize root. Images were generated using the DIMPLE GUI, which shows the 2D figure for each line scan in the graphic shown in D. Scale bar = 10 pixels. **F)** The DIMPLE method identified 301 unique *m/z* values compared to the 63 that were manually identified and included in the earlier Zhang et al. 2023 paper. All but two of the peaks in the previous method were identified by DIMPLE. For the full list of values see Supplemental Table 1 and Supplemental Table 2. **G)** DIMPLE identified 48 mass signatures grouped to Cluster 1 across three B73 replicates. Twenty-eight root tip enriched metabolites were identified in Zhang et al. 2023. Twenty-two of the previously identified peaks were included in Cluster 1, four were grouped to Cluster 3 (see Supplemental Figure 4) and two were not identified by DIMPLE. Images and peak lists can be found in Supplemental Figure 3, 4 and Supplemental Table 3.

To confirm that DIMPLE can recover previously identified peaks from our datasets, we compared the localization patterns of several high enrichment metabolites from the DIMPLE GUI to MSiReader generated images. Specifically, we imaged malate, succinate and fumarate, which have distinct localization patterns, and are represented in all the samples we have previously analyzed (Figure 1E) (T. Zhang et al., 2023). We found that DIMPLE generated images like those obtained with MSiReader (Figure 1E, Supplemental Figure 2). The main differences are that the DIMPLE images are not normalized and tend to have more empty pixels which are the result of background suppression steps during analysis of the raw spectrum.

The enrichment patterns represented in each cluster generally correspond to known developmental domains along the root axis. The clusters we generated for B73 maize roots encompass four metabolite distribution patterns, which we term Clusters 1-4: *Cluster 1)* strong meristem enrichment, *Cluster 2)* distribution throughout the meristem and transition zone (TZ), *Cluster 3)* peak enrichment in the TZ and EZ, and *Cluster 4)* strong enrichment in non-meristematic, differentiating tissue. Of note is the varied enrichment patterns found in Cluster 2, with approximately 40-78% of the signatures in this cluster showing characteristics of noise upon visual inspection of the 2D images (Figure 1D, Supplemental Figure 1).

Overall, DIMPLE enables the visualization of hundreds of *m/z* signatures throughout the developmental gradient. It additionally provides information on how metabolites are related to one another through hierarchical clustering of the linear patterns of enrichment along the root’s developmental axis. This provides a useful approach to identify uncharacterized compounds with specific developmental localization patterns.

### DIMPLE identifies metabolites with distinct localization patterns

To validate DIMPLE’s clustering and utility, we applied it to previously published data sets in B73 maize roots. DIMPLE was able to identify an average of 196 *m/z* signatures per root (overall it identified 301 unique *m/z* values in three replicates (Supplemental Tables 1, 2)). 63 peaks were included in the original paper, which were manually curated and excluded noise (Figure 1F, Supplemental Table 1). DIMPLE identified 97% of the *m/z* signatures published in the Zhang et al. 2023 paper. The two signatures that were not identified by DIMPLE in the final processed lists were either at very low abundance in the replicates investigated here or were only detected in a small number of pixels and thus removed by the image sparsity filter. We also used DIMPLE to map previously uncharacterized mass signatures localized to specific regions of the root. DIMPLE assigned 48 mass signatures to Cluster 1 across three B73 replicates, excluding peaks that were noise upon 2D visualization. 28 meristem and root tip enriched mass signatures were previously identified in Zhang et al. 2023. Of the 28 mass signatures identified previously, 22 are included in Cluster 1 peaks, four were grouped to Cluster 3 peaks, and two were not pulled out by DIMPLE (Figure 1G, Supplemental Table 3). DIMPLE provided 26 new root tip mass signatures to characterize (Supplemental Table 3, Supplemental Figure 3).

Notably, more signatures can be identified using DIMPLE by relaxing parameters such as the ppm tolerance (increasing the ppm threshold from 2.5 to 5 ppm increases the average by 14%, to 223 *m/z* peaks per B73 root) or the minimum threshold enrichment compared to background (decreasing this to 1.1 from 1.5 slightly increases the average by 3%, to 199 *m/z* peaks per B73 root). Relaxing these parameters can provide more targets but should be analyzed carefully to identify any false signal or noise selected by running a less stringent analysis. Additionally, including the sparse signal removed peaks can result in over 10-fold increase in the number of peaks identified. However, unless the goal of analysis is to generate a completely comprehensive list of signals from the MSI run, it is not recommended that the sparse signal removal step be skipped, as many of these peaks may be noise or overlapping signals for the same mass signature.

Our analysis using DIMPLE resulted in the discovery of new enrichment patterns along the developmental axis. For example, the peaks in Cluster 1 of B73 roots (Figure 2 A, B), have high enrichment in the root tip and the meristem. Additionally, as evidenced by the four root-tip localized peaks identified in Zhang et al. that were associated with a different cluster, the DIMPLE clustering method can provide a more comprehensive analysis of localization patterns. For example, both *m/z* 149.0118 and succinate (*m/z* 117.0195) were manually assigned as root tip and/or meristem-enriched compounds (T. Zhang et al., 2023). However, while DIMPLE consistently clusters *m/z* 149.0118 with other metabolites that are strongly enriched in the root tip (Cluster 1) (Figure 2A, B, Supplemental Tables 4-6), succinate is assigned to Cluster 2 in two of our replicates (Figure 2A, B, Figure 1D, E, Supplemental Figure 1, 2, Supplemental Tables 4-6). When comparing the metabolite localization of succinate and *m/z* 149.0118, succinate is more enriched throughout the developmental axis, while *m/z* 149.0118 is highly specific to the root tip region (Figure 2A). DIMPLE can parse these differences and this specificity in localization characterization may help to better identify compounds that are linked in developmental function through shared enrichment. In addition to succinate, several other root tip and/or meristem-enriched mass signatures previously identified, such as a molecule predicted to be succinic anhydride (*m/z* 99.0090) also show enrichment along the transition from the root tip to the meristem and EZ (Supplemental Figure 4), as shown in Figure 2A, and are clustered correspondingly to Cluster 3 instead of Cluster 1. This indicates that, similarly to the succinate pattern, while these mass signatures have enrichment in the meristem, their distribution patterns are more closely related with peaks concentrated in the region of transition between the meristem and EZ, based on the hierarchical clustering method (Figure 1C, Supplemental Figure 1, 4). This highlights the more nuanced and holistic approach that DIMPLE provides for mapping the chemistry of developmental transitions, as well as the importance of closely investigating enrichment patterns in all designated clusters to identify compounds of interest.

**Figure 2.**
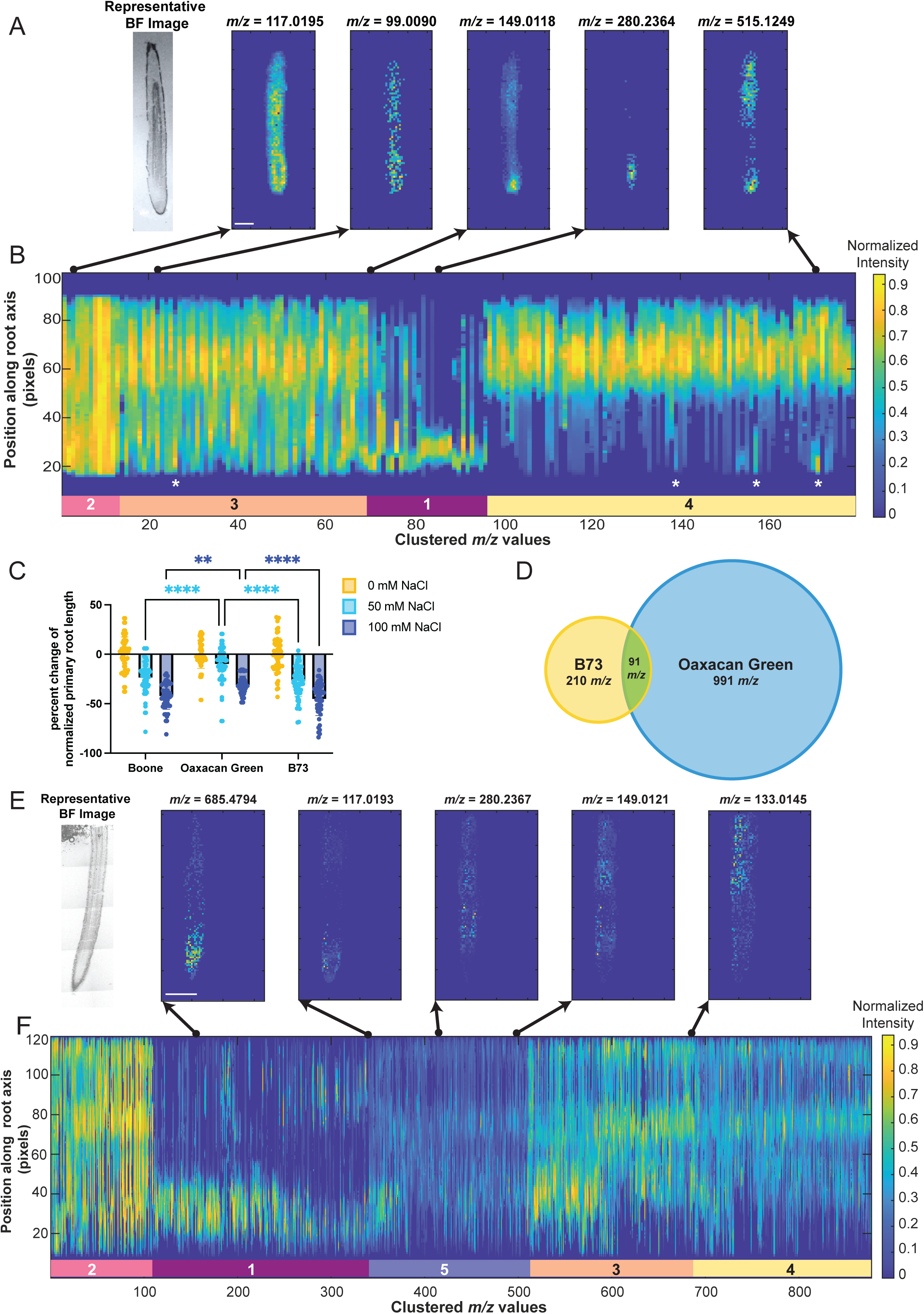
DIMPLE identifies key developmental differences within and between maize varieties. **A)** From left to right: representative brightfield image of B73 maize cryosection used for DESI-MSI, and the DIMPLE-generated MSI images for *m/z* 117.0195 (succinate), *m/z* 99.0090, *m/z* 149.0118, *m/z* 280.2364, and *m/z* 515.1249 in a B73 maize root. Scale bar is 10 pixels. Images were taken at approximately 80 µm resolution. **B)** Linescan graph for the 1D intensity profiles of mass signatures identified in a B73 maize root. Clusters are designated by the colored bars and labelled with the corresponding cluster number. The y-axis corresponds to position along the root axis (pixels) where signal closest to 0 is the root tip and signal closest to 100 corresponds to the mature root tissue. The x-axis represents the clustered *m/z* linescans arranged smallest to largest *m/z* value within each cluster. The arrows point to the 2D image in A that corresponds to that position in the linescan. White stars denote observed areas with bimodal patterns in the linescan. The intensity is normalized to 1. Intensity value is represented according to the colored “Normalized Intensity” legend. **C)** Percent change of normalized primary root length for Oaxacan Green, Boone and B73 maize under control, 50 mM and 100 mM NaCl conditions. 2-way ANOVA interaction p-value <0.0001 = ****, p-value <0.005 = **. Sample size for all varieties was n= 60 for 0 mM NaCl and 50 mM NaCl conditions, and n=48 for 100 mM NaCl conditions. **D)** DIMPLE identified 301 unique mass signatures in three B73 root replicates, and 1082 unique mass signatures in three Oaxacan Green replicates. Ninety-one of these mass signatures were conserved between the two varieties. For full peak lists see Supplemental Tables 2, 8 and 9. **E)** From left to right: representative brightfield image of a Oaxacan Green root cryosection, DIMPLE-generated MSI images for *m/z* 685.4794, *m/z* 117.0193 (succinate), *m/z* 280.2367, *m/z* 149.0121, and *m/z* 133.0145 (malate) in a Oaxacan Green maize root. Scale bar is 10 pixels. Images were taken at approximately 50 µm resolution. **F)** Linescan graph of the 1D intensity profiles of mass signatures identified in a Oaxacan Green root. Clusters are designated by the colored bars and labelled with the corresponding cluster number. The y-axis corresponds to position along the root axis (pixels) with 0 closest to the meristem and 120 closest to the mature root tissue. The x-axis corresponds to clustered *m/z* linescans arranged largest to smallest *m/z* value within each cluster. Cluster 5 (lavender) is specific to Oaxacan Green and contains signatures with low signal enriched in the transition and elongation zone. Arrows point to the 2D image in E that corresponds to that position in the linescan. The intensity is normalized to 1. Intensity value is represented according to the colored “Normalized Intensity” legend.

Building off the observation that meristem-enriched metabolites could be grouped outside of the expected meristem enriched cluster (Cluster 1), we identified several bimodally distributed metabolite patterns. Clusters 3 and 4 have several *m/z* signatures with strong enrichment in the meristem region, decreased signal in the TZ, and then strong signal again in the maturing tissue (Figure 2 A, B: white stars, Supplemental Figure 5, Supplemental Table 7). Scanning through these clusters with the GUI identified at least 16 signatures with distinct bimodal distributions, defined by enrichment in the meristem and then again in the mature elongation region (Figure 2A, Supplemental Figure 5, Supplemental Table 7). Of note, not all *m/z* features showed a bimodal pattern across all three replicates (Supplemental Table 7), which could be due to differences in sectioning, imaging artifacts or biological variation. However, this is an interesting developmental pattern as it might be expected that the metabolic demands of stem cells and differentiating tissue would be distinct. Further investigation into the identity of these bimodal metabolites could reveal new understanding of metabolite-driven regulation of distinct developmental stages.

In addition to revealing more nuanced developmental patterns in the meristem region, DIMPLE also facilitates analyses of distinct patterns during differentiation. An observation in the supplement of Zhang et al. 2023 was that several mass signatures seemed localized to specific cell layers of the root axis such as the cortex or vasculature. While DIMPLE condenses 2D information into a 1D linescan to perform the clustering, the DIMPLE GUI recovers the 2D information by visualizing the chemical map corresponding to a selected feature in the linescan. Using the DIMPLE GUI to assess the localization patterns within clusters, we identified more than 15 mass signatures in Cluster 4 of B73 roots with specific enrichment in the cortex (Supplemental Figure 6, Supplemental Table 7), and more than 10 mass signatures with enrichment in the vasculature (Supplemental Figure 7, Supplemental Table 7). This demonstrates that DIMPLE can be used to investigate variations within developmental localization clusters and further provide uncharacterized patterns for future investigation.

DIMPLE allows for a more comprehensive analysis of untargeted MSI data. The clustering and GUI functions enable annotation and discovery of novel mass signatures with distinct localization patterns. The ability to scan through multiple biological replicates with the GUI enables identification of instrumentation or biological variation within samples that can inform hypotheses on conserved developmental localization. We have successfully used this method to identify dozens of new metabolite candidates with developmentally specific patterns that have exciting potential for uncovering novel biological functions. Overall, this demonstrates that DIMPLE is a useful tool for approaching the untargeted analysis of MSI data along the developmental root axis.

### Comparing metabolite localization and distribution across maize varieties with different stress-tolerances

The ability to cluster MSI data based on developmentally relevant patterns increases the speed and depth of our analysis pipeline. To utilize these advantages to uncover new biology, we sought to compare developmental chemistry across different maize genotypes. Specifically, we wanted to understand how the chemistry along the root developmental gradient differs between stress-resistant and stress-susceptible maize varieties. To do this, we tested several heirloom varieties (Supplemental Figure 8), including Oaxacan Green, which has a reputation for stress tolerance. We also tested Boone County Dent (Boone), a commercially available varietal, and the research line B73 for their response to salt stress (Figure 2C). We chose to compare Boone and B73 to the Oaxacan Green variety as it exhibited similar control primary root length at 5 days post germination to the Boone variety (Supplemental Figure 8). All varieties showed significantly reduced root growth in 50 mM NaCl treatment, however primary root growth in Oaxacan Green was significantly less sensitive to 50 mM salt treatment compared to Boone and B73 (Figure 2C, 2-way ANOVA interaction p value <0.0001). Oaxacan Green was also significantly less susceptible to salt stress under 100 mM NaCl conditions compared to Boone and B73 (Figure 2C, ANOVA, p value <0.0001, p value <0.005). Based on these differences in salt stress responses, we proceeded to perform DESI-MSI on Oaxacan Green to map how chemical localization changes across development in a genotype with resistance to salt stress.

We applied DIMPLE to newly generated DESI-MSI data for Oaxacan Green roots at approximately 50 µm resolution. To compare the metabolite composition of B73 and Oaxacan Green maize varieties, we pooled the DIMPLE-processed mass signatures (within 0.001 tolerance) for three replicates of each maize variety. There were 301 unique mass signatures identified across three B73 roots (Supplemental Table 2, 8), and 1082 unique mass signatures identified in three Oaxacan Green roots (Figure 2D, Supplemental Table 8, 9). The increase in mass signatures in Oaxacan Green compared to B73 is likely due to instrumentation variability over time, which is a known challenge in MSI. HPLC-MS data suggests that B73 and Oaxacan Green produce similar numbers of mass signatures, with 1,110 and 1,139 unique mass signatures (within a 0.001 ppm tolerance) identified respectively for B73 and Oaxacan Green. An initial comparison revealed 737 of the HPLC-MS mass signatures were shared between varieties, with 383 unique to B73 and 412 unique to Oaxacan Green (Supplemental Table 10). A comparison (tolerance of 0.001) of the unique DESI-MSI mass signatures reported for B73 and Oaxacan Green identified 91 mass signatures conserved between the varieties, 210 mass signatures unique to B73 and 991 unique to Oaxacan Green (Figure 2D, Supplemental Table 8). The conserved signatures are interesting because they may represent a core set of metabolites in developing root tissue. The unique mass signatures identified in each variety, particularly Oaxacan Green, are also exciting candidates as they may confer different developmental responses to the environment.

To continue characterizing differences between the varieties, we compared the linescan graphs along the developmental axis of each root variety which revealed striking differences in the developmental localization patterns of metabolites (Figure 2A-B, E-F). B73 roots generally have four distinct clusters that align closely to developmental regions (Figure 2B, Supplemental Figure 1), as previously described. Oaxacan Green roots, however, typically group into five distinct clusters, the first four of which are generally consistent with the B73 clusters (Figure 2E-F, Supplemental Figure 9). In Oaxacan Green, the main clusters encompass: *Cluster 1)* enrichment in the meristem and TZ, *Cluster 2)* distribution throughout the root axis (in one sample, the intensity was primarily meristem localized in this cluster), *Cluster 3)* strong signal intensity in the TZ and EZ, *Cluster 4)* enrichment in the mature EZ, and finally *5)* overall low signal intensity in the TZ and EZ, with only a few pixels at normalized max intensity (Supplemental Figure 9, Supplemental Tables 11-13). In Oaxacan Green, there is more variation within each cluster, and further segmentation of these clusters could identify even more biologically meaningful localization patterns.

We focused on the 91 conserved *m/z* values identified in B73 and Oaxacan Green for our preliminary analysis of this data. The TCA metabolites malate, fumarate and succinate seem to maintain general localization trends between B73 and Oaxacan Green (Figure 1 D, E, Figure 2 A, B, E, F, Supplemental Figure 2, 10). Malate and fumarate both localize to the mature EZ (Figure 2E, Supplemental Figures 2, 10) and succinate is more strongly enriched in the meristem (Figure 2A, E, Supplemental Figures 2, 10). However, we found that several metabolites demonstrated different localization patterns in Oaxacan Green compared to B73. A noticeable difference is observed for meristem enriched metabolites. Unlike B73, in which Cluster 1 is composed mainly of metabolites concentrated in the meristem, Cluster 1 in Oaxacan Green is characterized by meristem and TZ enriched metabolites. Of the 48 meristem localized metabolites identified in B73 (Figure 1G), 20 are conserved in Oaxacan Green (Supplemental Table 3, 8). However, around half of these shared mass signatures do not share meristem localization patterns in Oaxacan Green (Supplemental Figure 11). For example, *m/z* 149.0118 shows specific enrichment in the root tip of B73 (Figure 2A, B), however it is localized more broadly in Oaxacan Green roots (Cluster 5) (Figure 2E, F). A similar pattern is observed for *m/z* 280.2367, which shows strong specific localization to the cortex of the EZ in Oaxacan Green roots (as viewed with total ion current (TIC) normalization in MSiReader) (Figure 2E). However, *m/z* 280.2367 is very specifically root tip enriched in B73 (Figure 2 A, Supplemental Figure 11). These differences do not appear to be related to root anatomy or growth, as B73 and Oaxacan Green roots have similar meristem anatomies. The differences in metabolic enrichment patterns between this stress-resilient and stress-susceptible variety are an exciting source of information for understanding how metabolite localization can lead to functional differences. In the next section, we will discuss a metabolite with different enrichment patterns and physiological effects in these two varieties.

Notably, DIMPLE can rapidly identify top candidates with potential developmental significance; however, it is not designed to test the effects of different normalization approaches. By comparing DIMPLE to MSiReader visualizations, we have found that most patterns remain conserved after TIC normalization. However, it is worth noting that several *m/z* patterns change after normalization (Supplemental Figure 12), emphasizing the importance of including a normalization step before inferring biological hypotheses. The biological outcomes of these variations in metabolite localization patterns will continue to be compelling points of investigation in the future.

### D-Erythrose, a compound identified through DIMPLE, improves the maize salt stress response

To determine if metabolites with different localization patterns in B73 and Oaxacan Green influence root developmental responses to stress, we focused on a compound with an *m/z* of 119.0350, which was identified in the maize root with DIMPLE. Using DIMPLE, we found *m/z* 119.0350 localized to Cluster 3 (TZ and EZ) in B73 (Figure 3A, Supplemental Figure 12, Supplemental Tables 4-6), and Clusters 1 (meristem) and 5 (low intensity enrichment in the TZ and EZ) in Oaxacan Green (Figure 3B, Supplemental Figure 12, Supplemental Tables 11-13). In Oaxacan Green the overall signal localization is present in the TZ and into the EZ but has stronger meristem enrichment and is more localized to the epidermis (Supplemental Figure 12). We visualized the *m/z* 119.0350 in MSiReader with TIC normalization: the enrichment of *m/z* 119.0350 shifts to be stronger in the meristem region in Oaxacan Green but remains enriched throughout the TZ and meristem in B73. (Figure 3 A-C, Supplemental Figure 12). However, the signal is still primarily localized to the epidermis in Oaxacan Green roots and is seemingly not enriched in any particular cell layer in the B73 roots. Since this is primarily a meristem and TZ enriched metabolite in diverse maize varieties, we hypothesized it might play a role in root growth during stress.

**Figure 3.**
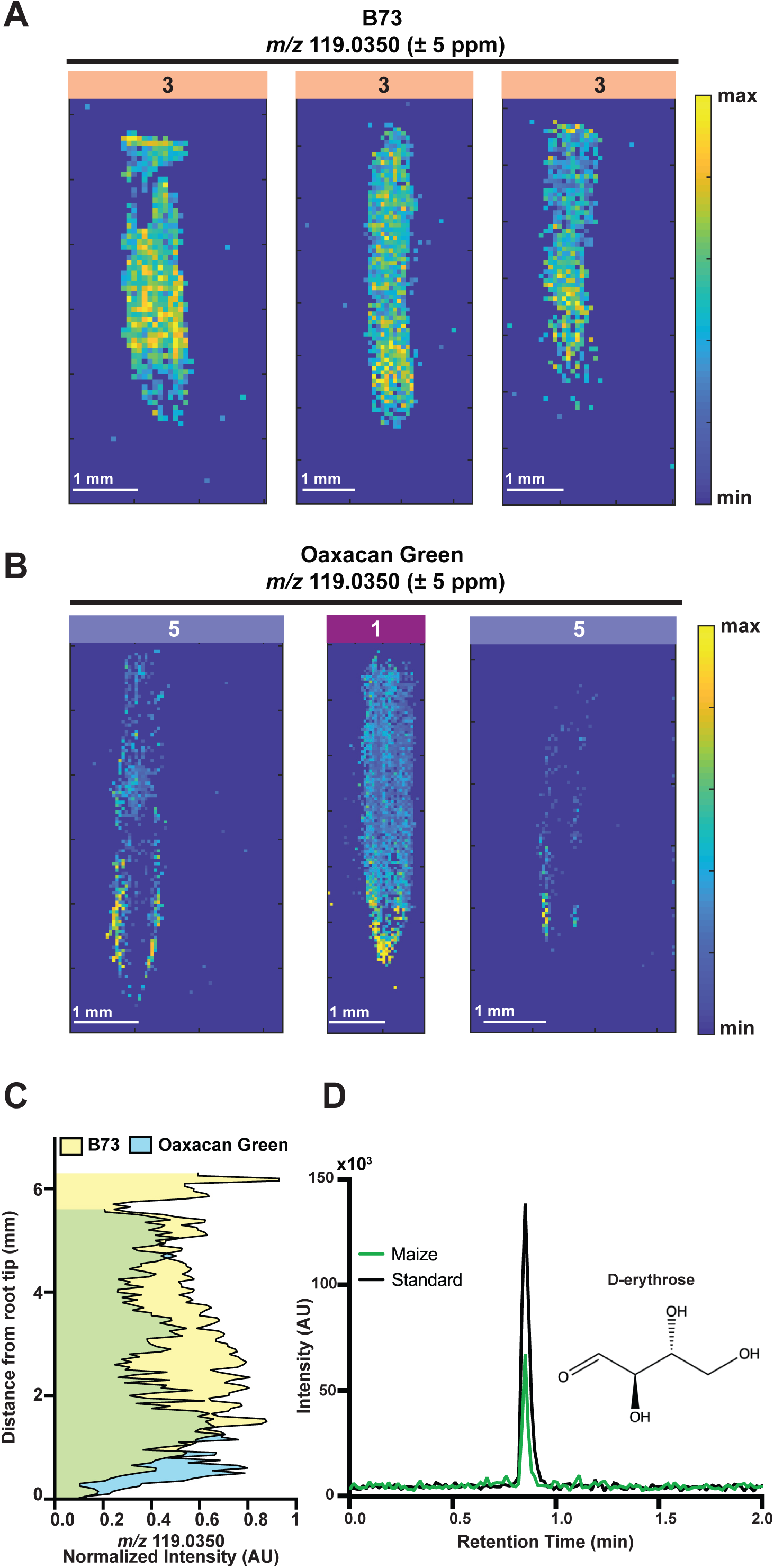
A DIMPLE metabolite candidate with different localization patterns in B73 and Oaxacan Green is identified as erythrose. **A)** Localization pattern of *m/z* 119.0350 +/- 5ppm in three B73 replicate roots. Each image was TIC normalized using MSiReader. *m/z* 119.0350 grouped to Cluster 3 in each B73 replicate. Scale bar = 1 mm. Intensity level is represented by the colored legend. **B)** Localization of *m/z* 119.0350 +/- 5 ppm in three Oaxacan Green replicate roots. Each image was TIC normalized using MSiReader. *m/z* 119.0350 grouped to Cluster 5 in two biological replicates and Cluster 1 in the other replicate. Scale bar = 1mm. Intensity level is represented by the colored legend. **C)** The intensity of *m/z* 119.0350 along each 2D image of three replicate roots per variety was measured in FIJI and normalized. The plot shows the normalized intensity along the root axis, with blue indicating Oaxacan Green, yellow indicating B73, and green showing the overlap. **D)** HPLC-MS/MS retention time for a D-erythrose standard (52 mM) (chemical structure shown) and the compound in Boone maize root extract.

Using multiple database searches, including Metlin, METASPACE, and LOTUS, we predicted this mass signature to be D-erythrose, a tetrose monosaccharide (Figure 3D). To validate the identity of this compound, we performed HPLC-MS/MS on Boone County Dent (Figure 3D, Supplemental Figure 13A) maize roots. DESI-MSI data shows that erythrose can be localized to the Boone meristem, as well (Supplementary Figure 13B). The *m/z* 119.0350 in the maize root extracts had the same MS/MS fragmentation and retention time as a D-erythrose standard, confirming that this *m/z* value likely corresponds to D-erythrose (Figure 3D, Supplemental Figure 13 A, B).

Literature search revealed that D-erythrose is largely uncharacterized in plants and has been studied mostly in the context of erythrose metabolism in bacteria, yeast and animals (Andriotis & Smith, 2019; den Hartog et al., 2010; Gallardo-Pérez et al., 2023; Lee et al., 2002; LIU et al., 2015; Park et al., 2011; Williams et al., 1980; J. Zhang et al., 2022). Past metabolic studies primarily have focused on erythrose-4-phosphate, an intermediate of the pentose phosphate pathway, shikimic acid pathway and the Calvin Cycle (Backhausen et al., 1997; Herrmann & Weaver, 1999; Kruger & von Schaewen, 2003; Maeda & Dudareva, 2012; Sharkey, 2021). Some connections for a role of erythrose and stress response have been identified: a study in yeast found that an erythrose reductase gene is upregulated under osmotic and salt stress conditions (Park et al., 2011). Another study of salt-stressed tomato plants inoculated with and without a plant growth-promoting rhizobacterium reported that erythrose accumulated in the inoculated plants prior to stress (Mellidou et al., 2021). After salt-stress, erythrose showed higher accumulation only in the non-inoculated plants (Mellidou et al., 2021). To further investigate a biological role of this compound, we performed exogenous treatments of D-erythrose on B73, and Boone combined with salt stress (Figure 4A). We found no significant improvement in primary root length in control conditions with D-erythrose treatment (Figure 4B, Supplemental Figure 14). However, we found that D-erythrose treatment significantly improved Boone and B73 primary root length at 50 mM NaCl and 100 mM NaCl respectively (Figure 4A-D). D-erythrose treatment combined with 50 mM NaCl improved Boone County Dent primary root length to that of control conditions (Figure 4C). This suggests that D-erythrose confers stress resistance to roots grown in salt, particularly in genotypes that have increased sensitivity to stress.

**Figure 4.**
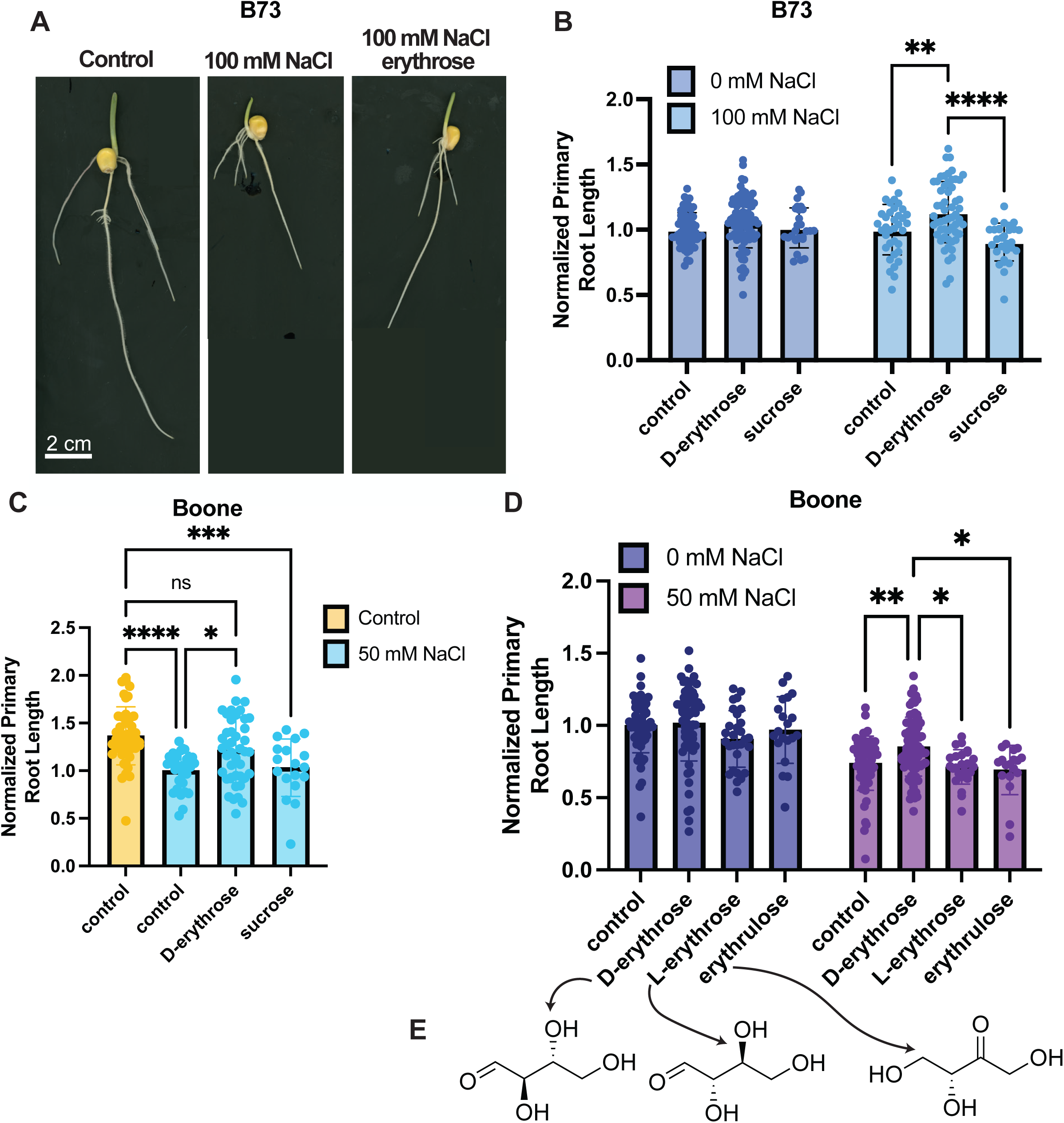
D-erythrose, identified with DIMPLE, improves primary root growth under salt conditions. **A)** Representative images of B73 day 5 seedlings in control, 100 mM NaCl or 100 mM NaCl and 200 µM D-erythrose treatment. Scale bar = 2 cm. **B)** Normalized B73 primary root length in control and 100 mM NaCl conditions with 200 µM D-erythrose or 200 µM sucrose treatment. Ordinary 2-way ANOVA Tukey’s multiple comparisons test, with a single pooled variance p-value < 0.0001 = ****, p-value = 0.0016 = **. Sample size for control conditions: control: n=58, D-erythrose: n=80, sucrose: n= 27, for NaCl conditions, control: n= 36, D-erythrose: n=56, sucrose: n= 29. **C)** Normalized primary root length to 50 mM NaCl control conditions, with control, 200 µM D-erythrose and 200 µM sucrose treatment. Ordinary 1-way ANOVA, Tukey’s multiple comparisons with single pooled variance, p-value <0.0001 = ****, p-value = 0.0005 = ***, p-value = 0.0142 = *. Sample size for control conditions: control n= 44. Sample size for 50 mM NaCl conditions: control n = 30, D-erythrose = 42, sucrose = 18. **D)** Normalized primary root length for Boone maize roots to control conditions. Control and 50 mM NaCl conditions treated with 200 µM D-erythrose, 200 µM erythrulose, and 200 µM L-erythrose treatment. Ordinary 2-way ANOVA, Šídák’s multiple comparisons test, with a single pooled variance, p-value < 0.05 = *, p-value < 0.01 = **. Sample size for control conditions, control n= 56, D-erythrose n= 63, L-erythrose n= 29, erythrulose n= 20. Sample size for 50 mM NaCl conditions: control n= 60, D-erythrose n= 78, L-erythrose n=24, erythrulose n=18. **E)** Molecular structures for D-erythrose, L-erythrose and erythrulose.

To confirm that the stress resilience phenotype was unique to D-erythrose treatment, we treated the plants with the same concentration of sucrose, as an energy-source control. If D-erythrose is enhancing growth due solely to providing additional carbohydrates to the plant, we would expect to see a similar phenotype in the sucrose treated plants. Our results show that D-erythrose treatment significantly enhances root growth compared to the sucrose treated plants and the control in both Boone County Dent and B73 at 50 mM NaCl and 100 mM NaCl respectively, (Figure 4B, Supplemental Figure 14) suggesting that D-erythrose has an effect outside of supplemental energy on growth during stress.

To investigate whether D-erythrose may be acting as an osmolyte, or another non-specific tetrose function, we included additional experiments testing D-erythrose isomers, L-erythrose, and erythrulose (Figure 4E). The D-erythrose treatment significantly increased growth compared to the L-erythrose and erythrulose treatments (Figure 4D). L-erythrose and erythrulose had insignificant effects compared to the control. These results strongly support our hypothesis that D-erythrose has a highly specific effect in increasing root growth during salt stress.

Further investigation of the biological mechanism that controls the D-erythrose response during salt stress will reveal whether there is a functional significance to this compound’s enrichment to the meristem and TZ regions and its mechanism of stress response. Understanding the mechanism of this molecule may also enable the development of new strategies for promoting growth in stress-susceptible crops. In addition, the DIMPLE method provided an abundance of compounds that have not been identified or characterized for their biological significance in plant root development or stress responses. Future work to gain insight into the function of these plant molecules holds great potential to discover novel regulators of development and stress.

## Conclusion

Overall, we have developed a computational method, DIMPLE, which leverages the linear developmental axis of plant roots to perform an untargeted screen for metabolites that have distinct developmental localization patterns. DIMPLE revealed mass signatures with specific localization to the root tip that we had not identified previously. The clustering method resulted in distinct groups corresponding generally to metabolites enriched in a variety of patterns across the developmental gradient of the B73 maize root. We then used DIMPLE to compare metabolite signatures of a salt-resilient maize variety, Oaxacan Green. We were able to identify distinct differences in the clustering between Oaxacan Green and B73. This led to the characterization of the metabolite, D-erythrose, with different localization patterns between varieties. We found that in salt-sensitive varieties, treatment with D-erythrose improves root growth during stress by increasing primary root length.

This application of DIMPLE demonstrates that it is a useful tool for identifying uncharacterized compounds and informing biological hypotheses by providing crucial context of the developmental patterns of these compounds. Here, we followed up on one of the many metabolites of interest that DIMPLE identified. The robust nature of this method results in dozens of metabolites that can continue to be characterized and investigated. We utilized DIMPLE to analyze maize root development, but DIMPLE can be applied to MSI data for many tissue types and research questions. It will be particularly useful for investigating metabolites that have differential enrichment patterns along a single linear axis. DIMPLE is also useful for hierarchical clustering of metabolites that share spatial patterns. For the data presented in this manuscript, this clustering method is useful for identifying mass signatures along distinct regions of development. For instance, DIMPLE roughly doubled the number of meristem-enriched metabolites that we were able to identify. Overall, DIMPLE provides an approach for reversibly converting two-dimensional MSI data to one-dimensional data for biological processes that occur along a single axis. This enables fast and statistically relevant analysis for testing specific hypotheses. The results from the maize root demonstrate that this tool identifies developmental patterns in MSI data. For example, other systems with linear developmental gradients include the proximal-distal axis of maize leaves, the crypt-villus axis in the intestine, and the antero-posterior axis in drosophila embryos.

This work provides a new tool for MSI analysis, which is valuable for direct measurements of cell metabolites in their native environment. As demonstrated here, this information can inform new hypotheses of metabolite functions in development. However, a major remaining challenge is that the majority of metabolites in both plants and animals remain unannotated. Therefore, to accelerate exploration of metabolite function, it is instrumental to improve the annotation of mass signatures across biological systems. The data presented here has sufficient annotations to begin the process of investigating how the precise enrichment patterns of metabolites in the maize root inform developmental functions. For example, understanding how D-erythrose patterning influences growth during stress will be an exciting avenue for future research. Future applications of DIMPLE could reveal metabolite patterns across different tissues, genotypes, and environmental conditions, with broad applications in chemical biology.

## Methods

### Maize sample preparation for DESI-MSI

The B73 data for DESI-MSI was prepared as described in Zhang et al. 2023. Boone and Oaxacan Green were first plated on autoclaved paper towels in a sterile 13×13 cm plate with 20 mL of DI water then transferred after two days and grown to five days in CYG Large Germination Pouches treated with 80 mL DI water. They were grown in a growth chamber set to 27°C with a 16h:8h day night cycle. The roots were cut at ∼5 mm from the tip, embedded in OCT and frozen on dry ice. The roots were kept at –80°C until they were removed and prepared for cryosectioning. Cryosections of 20 µm thickness were attached to microscope slides and imaged with brightfield microscopy. They were then packaged on dry ice and shipped to the Zare lab at Stanford for DESI-MSI. DESI-MSI experiments were performed using a commercial DESI sprayer (Viktor Tech, Beijing, China) coupled to an Orbitrap Velos Pro mass spectrometer (Thermo Fisher Scientific, Waltham, MA, USA). The DESI spray capillary had an inner diameter of 20 µm and an outer diameter of 150 µm. The sprayer was positioned at a 60° angle relative to the sample surface, with the capillary-to-surface distance set to 4 mm and the capillary-to-inlet distance at 2 mm. Compressed nitrogen (99.999%) was used as the nebulizing gas at 120 psi. A high-voltage of −6 kV was applied to the sprayer, and the solvent (DMF:ACN, 1:1) flow rate was maintained at 0.6 µL/min. Full-scan MS data were acquired in negative ion mode over an *m/z* range of 50–600, with a resolution of 30,000. The scan rate was fixed by disabling automatic gain control (AGC). The maximum injection time was set at 150 µs, and the ion transfer capillary temperature was 300 °C. Imaging was performed by raster-scanning the sample stage at a speed of 150 µm/s along the x-axis with a step size of 50 µm along the y-axis.

### Maize treatment experiments

For salt and erythrose treatments, B73, Boone and Oaxacan Green seeds were germinated on autoclaved paper towels in sterile 13×13 cm plates with corresponding 20 mL of salt (50 mM or 100 mM NaCl) and D-erythrose (200 µM) conditions, for two days. They were transferred on day two into CYG Large Germination Pouches with 80 mL of the same treatment conditions. They were grown in a growth chamber set to 27°C with a 16h:8h day night cycle. The plants were grown until day 5 when they were scanned, and primary root lengths were measured using FIJI. Primary root lengths were normalized to the control condition for each variety, and 2-way ANOVA was performed in PRISM to assess the statistical significance of primary root lengths.

For the erythrose isomer treatments, Boone seeds were germinated on autoclaved paper towels in sterile 13×13 cm plates with 20 mL of salt (50 mM NaCl), D-erythrose (200 µM), erythrulose (200 µM) or L-erythrose (200 µM) conditions. The seedlings were transferred on day two into CYG Large Germination Pouches with 80 mL of the same treatment conditions. They were grown in a growth chamber set to 27°C with a 16h:8h day night cycle. They were grown until day 5, the primary roots were scanned and measured using FIJI. Primary root lengths were normalized to the control condition for each variety, and 2-way ANOVA was performed in PRISM to assess the statistical significance of primary root lengths.

### ESI-HPLC-MS/MS Identification of D-erythrose

Detection and identification of a D-erythrose standard (52 mM) in maize roots was performed using electrospray ionization high-performance liquid chromatography tandem mass spectrometry (ESI-HPLC-MS/MS). Boone County Dent maize was germinated as described above. At day 5 the primary root meristems (0.5 cm in length) were harvested. 3 excised root tissues were transferred to microcentrifuge tubes with 500 µL of HPLC graded water and ground with a plastic pestle. Samples were shaken for an hour using an analog vortex mixer (Ohaus) then centrifuged at 2800 rpm for 5 minutes. The extract was then filtered through a 0.2-μm syringe filter into a clean 1.5 mL tube. Finally, 100 µL of the filtrate was added to HPLC sample vial for HPLC-MS analysis. All tissue extracts were prepared on the same day. Chromatographic separation was performed on a Thermo Scientific Vanquish UHPLC system coupled to an Orbitrap Elite mass spectrometer (Thermo Scientific) equipped with a heated electrospray ionization (HESI) source. A Phenomenex Luna® C18 column (100 × 2.1 mm, 1.6 µm particle size) was used for separation. The mobile phases were mobile phases A (H2O/FA, 1000/0.1, v/v) and B (ACN/FA, 1000/0.1, v/v) with the gradient program: 0-2 min, 0% B; 2-5 min, 0% B to 50% B; 5-7min, 50% B to 99% B; 7-10 min, 99% B; 10-10.1 min, 99% B TO 0% B; 10.1-13 min, 0% B. MS parameters included sheath gas flow rate of 50 arbitrary units (AU), auxiliary gas flow rate of 20 AU, spray voltage of -3.5 kV, capillary temperature of 300 °C, S-lens RF level of 62.4, and resolution of 30,000. The mass spectrometer was operated in negative ionization mode with data collected in centroid format. A targeted MS/MS method was used with two scan events. Both scan events used higher-energy collisional dissociation (HCD) fragmentation with a normalized collision energy of 35%, a maximum injection time of 100 ms, and an isolation window of 1.0 *m/z*. The first scan event was centered on a precursor at *m/z* 119.00, with fragment ions collected across an *m/z* range of 50-125. The second scan event was centered on a precursor at *m/z* 101.00, with fragment ions collected across an *m/z* range of 50-115. In both cases, the precursor charge state was set to 1, with no compensation voltage applied. Data acquisition and method control were performed using Thermo Xcalibur software.

### ESI-HPLC-MS/MS Identification of metabolite features in B73 and Oaxacan Green

5 days post germination, primary root tips (0.5 cm) were harvested from B73 and Oaxacan Green maize grown as described above in DI water. Sample preparation was conducted as described in the above section. Chromatographic separation was performed on a Thermo Scientific Vanquish UHPLC system coupled to an Orbitrap Elite mass spectrometer (Thermo Scientific) equipped with a heated electrospray ionization (HESI) source. A Phenomenex Kinetex® C18 reverse-phase column (150 x 2.1 mm, 1.7 µm particle size) was used for separation. The mobile phases were mobile phase A (H2O/FA, 1000/0.1, v/v) and mobile phase B (ACN/FA, 1000/0.1, v/v) with the following gradient program: 0–1 min, 0% B; 1–9 min, 0% B to 100% B; 9–11 min, 100% B; 11–11.5 min, 100% B to 0% B; 11.5–14 min, 0% B. The flow rate was maintained at 0.4 mL/min, with a total run time of 14 min. MS parameters included a spray voltage of –3.5 kV, sheath gas flow rate of 50 arbitrary units (AU), auxiliary gas flow rate of 20 AU, capillary temperature of 300 °C, S-lens RF level of 69.1 and resolution of 30,000. The mass spectrometer was operated in negative ionization mode, and data were collected in centroid format. A data-dependent top six scan MS/MS method was employed with seven scan events per duty cycle. The resulting MS/MS spectra were acquired at a resolution of 30,000. Data acquisition and method control were performed using Thermo Xcalibur software.

### R-code

R was used to compile a list of all the peaks present across three biological replicates. Unique peaks were defined within 0.001 ppm tolerance of each other. The compiled lists for B73 and Oaxacan Green were compared to identify shared and unique peaks between the two groups. The code was used to identify peaks for DIMPLE generated and ESI-HPLC-MS/MS data. The full R code analysis can be found in the Dickinson Lab Github linked in the Supplementary Information.

### Technical aspects of DIMPLE

Source code for DIMPLE is available on the Dickinson Lab Github linked in the Supplement. (https://github.com/dickinsonlab)

MSI data of each tissue section consists of a series of .raw files with each file containing the mass spectrum for a full row of pixels. To import MSI data into MATLAB, we converted the .raw files to .cdf using Xcalibur 3.0 Thermo File Converter. Then used the self-written MATLAB function ‘Batchcdfread’, provided by the Zare lab, to read the data with the appropriate number of .cdf files specified. This function is linked in the GitHub repository for the DIMPLE code. Each file contains one row of pixels that is imported into a nested MATLAB cell array by batchcdfread and reorganized into a 2D layout corresponding to the image. A sum-projection is calculated for each pixel and used to draw a mask for the tissue area of interest. Everything outside the masked region is considered background. Once a mask is selected, all peaks in each spectrum (pixel) are identified and assigned to *m/z* values based on the highest intensity for each peak. An array is then prepared with the summed intensity values corresponding to the center *m/z* value. A median filter is then applied to the summed spectrum to reduce noise and create a cleaner spectrum to detect meaningful features in the bulk spectrum. The filtered data is then downsampled to a 2.5 ppm tolerance window to identify significant peaks and avoid conflicts between closely spaced *m/z* values. The tolerance window can be adjusted to fit user’s instrument acquisition parameters. The background peaks, defined as peaks present outside of the mask, are similarly filtered and prepared. The sample peaks are compared to the background peaks and filtered depending on whether the sample intensity is greater than the minimum fold enrichment of the background peaks, for our analysis this was set to 1.5. This final set of peaks thus corresponds to matching center *m/z* positions across the entire image. The data is reorganized into a hyperspectral image where each ‘channel’ corresponds to the intensity distribution across the image at each unique *m/z* value.

The DIMPLE script includes code for exporting the hyperspectral image in a TIFF container for visualization and analysis using standard image processing tools such as ImageJ. DIMPLE itself performs simple analyses of the spatial distribution of signal in each channel. First, channels that are only sparsely present in a subset of pixels are removed using morphological erosion as channels where all pixels with > 0 intensity are exclusively surrounded by 0-valued pixels will be removed.

For the analysis of developmental patterns in plant roots, a line-scan is generated by performing a max projection along the lateral axis of the root. For pattern detection, each linescan is normalized to a maximum intensity of 1. The linescan data is then hierarchically clustered using Ward’s method with a Euclidean norm as the pair-wise measure of pattern similarity, to a defined number of cluster groups selected after inspecting the resulting dendrogram shape. The linescans in each cluster can be visualized with the linescan graph or using the linescan GUI. The GUI enables annotation of features in the linescan graph and recovers the 2D image for individual linescans. These can be exported as images.

## Supporting information

Supplemental Images

## Acknowledgments

We would like to thank Dr. Justin Whalley, Dr. Erin Sparks, Dr. Johannes Scharwies and Dr. José Dinneny for providing B73 seeds for our experiments. We would also like to thank Neal Arakawa of the Environmental and Complex Analysis Laboratory (ECAL) at UC San Diego for his advice, training, and assistance. Thank you to all the members of the Dickinson Lab for their helpful discussions during this work. This work was supported by the National Institute of Health (NIH grant R35GM151199 to P.S., NIGMS 1T32GM133351-01 to A.M.S, NIGMS 1R35GM147216 to A.J.D., and NIGMS 1T32GM127235-01 to A.T.), the Air Force Office of Scientific Research through the Multidisciplinary University Research Initiative program (AFOSR FA9550-21-1-0170) to R.N.Z., the Moore Foundation Inventor Fellowship to A.J.D., the Howard Hughes Medical Institute Gilliam Fellowship to A.M.S. and A.J.D and the University of California San Diego Graduate Division Cota Robles Fellowship to A.M.S.

## Author Contributions Statement

A.J.D., P.S., and A.M.S. supervised and designed the study. A.M.S., S.B.C., S.L., A.T., Y.M., and S.E.N. performed the experiments. P.S., wrote the DIMPLE script with consultation from A.M.S., and A.J.D.. A.M.S., A.J.D., P.S., S.B.C., S.L., and A.T. interpreted the data. A.M.S., P.S., and A.J.D. wrote the manuscript with the feedback from all authors.

## Conflict of Interest Statement

The authors declare that there is no conflict of personal or financial interests.

## Data Availability

The majority of the data can be found in the manuscript and supplementary materials. Raw data files and code can be accessed in our GitHub at https://github.com/dickinsonlab.

## Notes

https://github.com/dickinsonlab

### Competing Interest Statement

The authors have declared no competing interest.

