## Supplemental Tables for "Computational method for mapping mass signatures along developmental gradients reveals a novel role for a monosaccharide tetrose in maize salt-stress response"

**Supplemental Table 1** red highlight is peaks not identifed with DIMPLE

| Zhang et al. 2023 | DIMPLE |
|---|---|
| 71.0141 | 59.0142 |
| 72.9934 | 61.9887 |
| 99.009 | 66.1564 |
| 104.0355 | 69.7194 |
| 115.0038 | 71.0142 |
| 117.0195 | 72.9934 |
| 131.0463 | 73.0298 |
| 132.0303 | 74.025 |
| 133.0143 | 74.0407 |
| 145.0619 | 75.009 |
| 146.0459 | 76.9884 |
| 149.0118 | 85.0298 |
| 164.0352 | 87.009 |
| 165.0404 | 88.0407 |
| 173.0091 | 89.0247 |
| 179.056 | 92.9282 |
| 191.0559 | 96.9604 |
| 194.0457 | 96.9699 |
| 210.0406 | 99.009 |
| 210.9649 | 101.025 |
| 215.0325 | 102.056 |
| 225.0614 | 103.004 |
| 226.9961 | 103.04 |
| 240.051 | 104.036 |
| 246.0171 | 105.02 |
| 254.015 | 111.009 |
| 255.2326 | 112.986 |
| 268.1034 | 112.988 |
| 279.2323 | 113.025 |
| 280.2359 | 113.036 |
| 281.2483 | 114.02 |
| 284.0983 | 114.056 |
| 298.1139 | 115.004 |
| 306.0761 | 115.04 |
| 310.1502 | 115.077 |
| 311.1091 | 116.007 |
| 313.0772 | 116.036 |
| 323.0281 | 116.072 |
| 325.1247 | 117.02 |
| 330.2647 | 118.023 |

| | |
|---|---|
| 341.0597 | 118.051 |
| 354.2654 | 118.993 |
| 357.0336 | 119.035 |
| 359.1189 | 119.947 |
| 375.0538 | 120.031 |
| 377.0849 | 121.03 |
| 379.083 | 128.035 |
| 390.1034 | 129.02 |
| 391.0278 | 129.056 |
| 404.104 | 131.035 |
| 421.0844 | 131.046 |
| 435.2963 | 132.03 |
| 459.2958 | 133.014 |
| 490.2805 | 133.051 |
| 515.1244 | 134.018 |
| 552.1564 | 135.019 |
| 555.117 | 135.03 |
| 559.4724 | 138.02 |
| 571.091 | 138.056 |
| 719.2005 | 142.999 |
| 721.5019 | 143.108 |
| 745.5011 | 145.014 |
| 956.5492 | 145.062 |
| | 146.046 |
| | 146.065 |
| | 147.03 |
| | 147.049 |
| | 149.012 |
| | 149.046 |
| | 154.062 |
| | 156.067 |
| | 157.037 |
| | 157.051 |
| | 157.124 |
| | 160.062 |
| | 161.046 |
| | 162.982 |
| | 163.061 |
| | 164.035 |
| | 164.072 |
| | 164.63 |
| | 165.039 |

165.041
167.997
168.007
168.989
173.009
174.012
175.047
175.061
177.077
179.022
179.056
180.059
189.069
190.973
191.056
192.059
193.974
194.046
194.991
195.051
197.061
199.17
199.176
200.174
203.084
206.947
209.067
210.041
210.965
215.033
216.036
217.03
217.037
218.033
218.139
225.061
226.996
228.993
236.099
237.061
239.044
239.077

239.129
240.051
242.051
243.054
245.043
246.017
254.015
255.233
256.236
259.022
264.106
267.072
268.104
269.087
274.202
274.588
275.125
277.033
277.217
277.699
279.233
280.236
281.248
284.08
284.098
285.083
287.05
288.965
289.017
290.963
293.099
293.176
294.029
294.111
294.12
294.179
295.13
296.099
296.135
297.082
298.114
299.097

299.098
301.065
304.992
306.076
307.114
309.046
310.15
310.151
311.11
312.093
312.113
312.13
313.077
313.113
313.114
314.08
314.081
314.097
321.063
321.357
322.165
322.166
323.029
325.041
325.125
326.108
326.11
326.128
327.045
329.108
330.265
333.058
334.125
336.13
341.06
341.108
344.135
346.055
349.153
349.154
353.071
353.073

354.265
357.034
359.119
360.123
363.042
367.104
367.106
368.976
371.119
372.122
375.054
376.057
377.084
377.085
378.088
379.082
379.083
379.164
379.234
382.442
383.083
387.01
387.115
390.104
390.117
391.028
392.031
395.095
397.093
402.103
404.104
406.039
412.966
413.969
421.089
435.296
438.081
439.086
441.203
448.986
451.099
456.171

456.173
459.295
459.297
469.028
471.259
473.162
473.238
475.13
480.093
487.177
488.162
490.281
503.092
504.224
515.125
521.172
526.119
531.097
533.172
535.472
537.107
543.104
552.155
553.08
555.117
556.122
557.148
559.078
559.473
560.476
563.334
571.092
572.096
579.33
586.101
593.03
602.075
671.465
672.468
684.395
695.466
696.468

721.502
722.505
745.501
746.505
747.508
833.518
851.529
925.563
982.094

Supplemental Table 2

| B73-Root 1 | B73-Root 2 | B73-Root 3 |
|---|---|---|
| 59.01421 | 59.014126 | 59.014133 |
| 61.988682 | 69.719414 | 61.988682 |
| 71.014236 | 71.014236 | 66.156418 |
| 72.993477 | 72.993477 | 71.014122 |
| 73.029884 | 73.029877 | 72.993347 |
| 74.04068 | 74.025032 | 73.029762 |
| 85.029892 | 85.029732 | 75.009003 |
| 87.009048 | 87.009048 | 76.988358 |
| 88.040779 | 89.024643 | 85.029739 |
| 89.024803 | 96.969933 | 87.009048 |
| 96.969925 | 99.009033 | 88.040619 |
| 99.009033 | 101.02468 | 89.024643 |
| 101.02469 | 103.0039 | 92.928238 |
| 102.05634 | 104.03566 | 96.960381 |
| 103.00391 | 105.01953 | 99.009041 |
| 104.03566 | 112.98586 | 101.02467 |
| 105.01953 | 113.03577 | 102.05634 |
| 111.009 | 114.0199 | 103.0039 |
| 112.98835 | 114.05623 | 103.04025 |
| 113.02465 | 115.00388 | 104.03545 |
| 115.00388 | 116.00727 | 105.01954 |
| 116.00726 | 117.01955 | 111.009 |
| 116.03559 | 118.05118 | 112.98813 |
| 117.01954 | 119.03517 | 115.00388 |
| 118.05118 | 120.03053 | 115.04022 |
| 118.99276 | 129.0195 | 115.07657 |
| 119.03518 | 131.03528 | 116.00727 |
| 120.03055 | 131.04631 | 116.07195 |
| 121.02964 | 132.03036 | 117.01955 |
| 129.01952 | 133.01428 | 118.02284 |
| 131.03526 | 133.05077 | 118.05119 |
| 131.04631 | 134.01768 | 119.03518 |
| 132.03036 | 135.01859 | 119.94714 |
| 133.01456 | 135.03014 | 121.02963 |
| 133.05077 | 145.0144 | 128.03546 |
| 134.0177 | 145.0619 | 129.01952 |
| 135.01859 | 146.04585 | 129.05579 |
| 135.03014 | 146.06522 | 131.03497 |
| 145.0144 | 147.03009 | 131.04631 |
| 145.06192 | 149.01183 | 132.03036 |
| 146.04585 | 149.04549 | 133.01427 |
| 146.06552 | 156.06664 | 133.05077 |

| | | |
|---|---|---|
| 147.03009 | 157.03694 | 134.01768 |
| 149.01183 | 157.12357 | 135.01859 |
| 149.04582 | 160.06165 | 135.02985 |
| 154.06248 | 161.04558 | 138.01964 |
| 160.06165 | 162.98227 | 138.05608 |
| 161.04558 | 163.06128 | 142.99854 |
| 162.98268 | 164.03535 | 143.10774 |
| 164.03535 | 164.63013 | 145.0144 |
| 164.07187 | 165.04059 | 145.0619 |
| 165.04059 | 167.99716 | 146.04585 |
| 167.99716 | 168.9893 | 146.0652 |
| 173.00922 | 173.00922 | 147.03009 |
| 174.01277 | 174.01233 | 147.0493 |
| 175.06137 | 175.06137 | 149.01183 |
| 177.0768 | 179.02225 | 149.04549 |
| 179.02225 | 179.05618 | 154.06212 |
| 179.05618 | 189.0688 | 156.06662 |
| 180.05948 | 190.97281 | 157.05067 |
| 191.05609 | 191.05608 | 157.12357 |
| 194.04567 | 192.05945 | 160.06165 |
| 194.99097 | 194.04568 | 161.04558 |
| 195.05115 | 194.99095 | 162.98228 |
| 197.06058 | 195.05116 | 163.06129 |
| 199.17603 | 200.17355 | 164.03535 |
| 203.08456 | 206.94672 | 165.03899 |
| 206.94672 | 209.06673 | 165.04059 |
| 209.06673 | 210.04065 | 167.99715 |
| 210.04063 | 215.03282 | 168.00664 |
| 210.96521 | 216.03618 | 168.98932 |
| 215.03282 | 217.02963 | 173.00922 |
| 216.03616 | 217.0369 | 174.01233 |
| 217.02963 | 218.03302 | 175.04738 |
| 217.03687 | 225.06171 | 175.06093 |
| 218.03302 | 226.99594 | 179.02225 |
| 225.06171 | 239.12883 | 179.05618 |
| 226.99661 | 242.05142 | 180.05948 |
| 228.99319 | 245.04311 | 189.0688 |
| 236.09914 | 255.2328 | 190.97281 |
| 239.07716 | 256.23608 | 191.05608 |
| 240.05121 | 259.02243 | 193.97374 |
| 242.05142 | 264.1055 | 194.04568 |
| 245.04312 | 267.07187 | 194.99098 |
| 246.01746 | 268.10355 | 195.05064 |
| 254.01509 | 269.08783 | 197.06058 |

| | | |
|---|---|---|
| 255.23282 | 274.20215 | 199.1702 |
| 256.23608 | 277.03305 | 203.08401 |
| 259.02243 | 277.21689 | 210.04065 |
| 268.10355 | 279.2327 | 210.96463 |
| 277.03308 | 280.23642 | 215.03282 |
| 277.69867 | 281.24826 | 216.03616 |
| 279.23267 | 284.09879 | 217.02963 |
| 280.23642 | 287.04984 | 218.03302 |
| 281.24826 | 289.01712 | 218.13947 |
| 284.0798 | 293.09912 | 225.06105 |
| 284.09879 | 293.1759 | 226.99594 |
| 288.96521 | 294.02905 | 228.99319 |
| 290.96259 | 294.11954 | 236.09914 |
| 294.11954 | 294.17862 | 237.06125 |
| 296.13507 | 295.12997 | 239.04367 |
| 297.08243 | 296.09851 | 239.07718 |
| 298.11401 | 296.13507 | 239.12883 |
| 306.07678 | 297.08243 | 240.05124 |
| 310.15103 | 298.11401 | 242.05142 |
| 311.1095 | 299.09729 | 243.05446 |
| 312.11301 | 301.06458 | 245.04312 |
| 313.07742 | 304.99158 | 246.01746 |
| 313.11404 | 306.07581 | 254.01508 |
| 314.08102 | 307.11395 | 255.2328 |
| 321.35748 | 310.15103 | 259.02243 |
| 322.16626 | 311.10953 | 268.10352 |
| 323.02853 | 312.11301 | 269.08704 |
| 325.04111 | 312.12967 | 274.58844 |
| 325.12521 | 313.07739 | 275.12457 |
| 326.1095 | 313.11407 | 277.03305 |
| 326.12836 | 314.08105 | 277.69864 |
| 327.04459 | 321.06281 | 279.23267 |
| 330.26492 | 322.16626 | 280.23553 |
| 336.12952 | 323.02853 | 281.24826 |
| 341.05951 | 325.04108 | 284.0979 |
| 341.10828 | 325.12518 | 285.08255 |
| 344.13483 | 326.1095 | 293.17587 |
| 349.1539 | 326.12839 | 294.11099 |
| 353.07269 | 327.04459 | 294.17859 |
| 354.26468 | 329.1084 | 296.13507 |
| 357.0336 | 330.26495 | 297.08243 |
| 359.11938 | 333.05847 | 298.11401 |
| 360.12268 | 336.12952 | 299.0983 |
| 367.10556 | 341.05951 | 301.06461 |

| | | |
|---|---|---|
| 368.97638 | 341.10831 | 306.07581 |
| 371.11923 | 346.05493 | 309.04611 |
| 375.05365 | 353.07269 | 310.14999 |
| 376.05704 | 354.26471 | 311.10953 |
| 377.08517 | 357.0336 | 312.0932 |
| 378.08838 | 359.11938 | 313.07745 |
| 379.08301 | 367.10556 | 313.11301 |
| 379.23361 | 371.1192 | 314.07996 |
| 387.11456 | 372.12177 | 314.0968 |
| 390.10352 | 375.05365 | 321.35748 |
| 391.02774 | 377.08517 | 322.16516 |
| 392.03137 | 378.08838 | 323.0285 |
| 395.09543 | 379.08304 | 325.04111 |
| 397.09332 | 387.00955 | 325.12521 |
| 402.10309 | 387.11453 | 326.1084 |
| 404.10385 | 390.10352 | 326.12842 |
| 412.96552 | 390.11661 | 327.04459 |
| 413.96899 | 391.02774 | 329.1084 |
| 421.08862 | 395.09543 | 330.26495 |
| 435.2962 | 397.09335 | 333.05841 |
| 438.08121 | 404.10388 | 334.12512 |
| 439.08557 | 412.96552 | 336.12955 |
| 451.09885 | 413.96896 | 341.05951 |
| 456.17316 | 435.29623 | 341.10828 |
| 459.29691 | 439.08557 | 344.13483 |
| 469.02795 | 441.20261 | 346.05496 |
| 473.16229 | 456.17133 | 349.15265 |
| 473.2381 | 459.29507 | 353.07141 |
| 475.12967 | 473.16229 | 354.26474 |
| 480.09314 | 473.2381 | 357.0336 |
| 487.17743 | 475.12964 | 359.11938 |
| 488.1618 | 480.09314 | 363.04166 |
| 490.28091 | 487.17743 | 367.10425 |
| 515.12488 | 488.1618 | 371.11926 |
| 521.17224 | 490.28091 | 375.05365 |
| 537.10675 | 503.09219 | 376.05704 |
| 552.15533 | 504.2236 | 377.0838 |
| 553.0802 | 515.12494 | 377.08517 |
| 555.11737 | 521.17224 | 379.08163 |
| 557.14832 | 526.11908 | 379.16394 |
| 571.0918 | 531.09735 | 379.23361 |
| 572.09576 | 533.17218 | 382.44217 |
| 593.02979 | 535.47205 | 383.08279 |
| 602.07495 | 537.10675 | 387.11456 |

| | | |
|---|---|---|
| 671.46515 | 555.11737 | 390.10358 |
| 721.50159 | 559.47278 | 391.02774 |
| 745.50079 | 560.47626 | 392.0314 |
| 746.50525 | 571.09186 | 395.09543 |
| 982.09424 | 593.02979 | 397.09332 |
| | 671.46515 | 404.10382 |
| | 672.46844 | 406.03918 |
| | 695.46619 | 413.96899 |
| | 696.46832 | 421.08862 |
| | 721.50159 | 435.29623 |
| | 722.50549 | 439.0856 |
| | 745.50085 | 448.98566 |
| | 746.50525 | 451.09882 |
| | 747.50769 | 456.17136 |
| | 833.51825 | 459.29507 |
| | 851.52899 | 471.25928 |
| | 925.56348 | 473.2381 |
| | | 480.09314 |
| | | 487.1774 |
| | | 488.1618 |
| | | 490.28091 |
| | | 503.09222 |
| | | 504.22357 |
| | | 515.12488 |
| | | 526.11908 |
| | | 543.10406 |
| | | 552.15533 |
| | | 555.11737 |
| | | 556.12152 |
| | | 559.07813 |
| | | 563.33392 |
| | | 571.0918 |
| | | 579.32965 |
| | | 586.10095 |
| | | 593.02979 |
| | | 602.07495 |
| | | 671.46521 |
| | | 672.46844 |
| | | 684.39508 |
| | | 695.46619 |
| | | 696.46826 |
| | | 721.50165 |
| | | 745.50085 |
| | | 746.50519 |

833.51831

**Supplemental Table 3**

blue highlight are peaks not included in DIMPLE cluster 1

red highlight are peaks not identified by DIMPLE

| Zhang et al. root tip ID | DIMPLE: purple cluster |
|---|---|
| 99.009 | 73.0298 |
| 117.0195 | 117.0195 |
| 149.0118 | 118.0228 |
| 164.0352 | 138.0561 |
| 194.0457 | 149.0118 |
| 210.0406 | 157.0507 |
| 226.9961 | 164.0354 |
| 240.051 | 165.039 |
| 246.0171 | 175.0614 |
| 255.2326 | 177.0768 |
| 279.2323 | 179.0222 |
| 280.2359 | 194.0457 |
| 281.2483 | 210.0406 |
| 330.2647 | 228.9932 |
| 354.2654 | 240.0512 |
| 377.0849 | 246.0175 |
| 379.083 | 254.0151 |
| 390.1032 | 255.2328 |
| 404.104 | 256.2361 |
| 421.0844 | 277.2169 |
| 490.2805 | 279.2327 |
| 515.1244 | 280.2364 |
| 552.1564 | 281.2483 |
| 559.4724 | 309.0461 |
| 719.2005 | 330.265 |
| 721.5019 | 341.1083 |
| 745.5011 | 354.2647 |
| 956.5492 | 377.0838 |
| | 377.0852 |
| | 379.0816 |
| | 387.1146 |
| | 402.1031 |
| | 404.1038 |
| | 421.0886 |
| | 435.2962 |
| | 451.0988 |
| | 459.2969 |
| | 471.2593 |
| | 487.1774 |
| | 488.1618 |
| | 490.2809 |

| |
|---|
| 504.2236 |
| 515.1249 |
| 535.472 |
| 552.1553 |
| 559.4728 |
| 560.4763 |
| 721.5016 |

**Supplemental Table 4**

| Cluster | m/z value |
|---|---|
| 2 | 74.04068 |
| 2 | 105.0195 |
| 2 | 117.0195 |
| 2 | 118.9928 |
| 2 | 120.0306 |
| 2 | 121.0296 |
| 2 | 162.9827 |
| 2 | 236.0991 |
| 2 | 284.0798 |
| 2 | 321.3575 |
| 2 | 349.1539 |
| 2 | 412.9655 |
| 2 | 982.0942 |
| 3 | 59.01421 |
| 3 | 61.98868 |
| 3 | 71.01424 |
| 3 | 72.99348 |
| 3 | 73.02988 |
| 3 | 85.02989 |
| 3 | 87.00905 |
| 3 | 89.0248 |
| 3 | 99.00903 |
| 3 | 101.0247 |
| 3 | 103.0039 |
| 3 | 111.009 |
| 3 | 112.9884 |
| 3 | 115.0039 |
| 3 | 116.0356 |
| 3 | 119.0352 |
| 3 | 129.0195 |
| 3 | 131.0353 |
| 3 | 132.0304 |
| 3 | 133.0146 |
| 3 | 133.0508 |
| 3 | 134.0177 |
| 3 | 135.0186 |
| 3 | 135.0301 |
| 3 | 145.0144 |
| 3 | 147.0301 |
| 3 | 149.0458 |
| 3 | 160.0617 |
| 3 | 161.0456 |

| | |
|---|---|
| 3 | 165.0406 |
| 3 | 167.9972 |
| 3 | 173.0092 |
| 3 | 174.0128 |
| 3 | 191.0561 |
| 3 | 195.0512 |
| 3 | 197.0606 |
| 3 | 203.0846 |
| 3 | 226.9966 |
| 3 | 242.0514 |
| 3 | 277.6987 |
| 3 | 297.0824 |
| 3 | 313.114 |
| 3 | 322.1663 |
| 3 | 325.0411 |
| 3 | 336.1295 |
| 3 | 341.1083 |
| 3 | 368.9764 |
| 3 | 371.1192 |
| 3 | 390.1035 |
| 3 | 404.1039 |
| 3 | 413.969 |
| 3 | 473.2381 |
| 3 | 593.0298 |
| 3 | 671.4652 |
| 3 | 745.5008 |
| 3 | 746.5053 |
| 1 | 149.0118 |
| 1 | 164.0354 |
| 1 | 175.0614 |
| 1 | 177.0768 |
| 1 | 179.0223 |
| 1 | 194.0457 |
| 1 | 199.176 |
| 1 | 210.0406 |
| 1 | 228.9932 |
| 1 | 240.0512 |
| 1 | 246.0175 |
| 1 | 254.0151 |
| 1 | 255.2328 |
| 1 | 256.2361 |
| 1 | 279.2327 |
| 1 | 280.2364 |
| 1 | 281.2483 |

| | |
|---|---|
| 1 | 330.2649 |
| 1 | 354.2647 |
| 1 | 402.1031 |
| 1 | 421.0886 |
| 1 | 435.2962 |
| 1 | 451.0989 |
| 1 | 459.2969 |
| 1 | 490.2809 |
| 1 | 552.1553 |
| 1 | 721.5016 |
| 4 | 88.04078 |
| 4 | 96.96993 |
| 4 | 102.0563 |
| 4 | 104.0357 |
| 4 | 113.0247 |
| 4 | 116.0073 |
| 4 | 118.0512 |
| 4 | 131.0463 |
| 4 | 145.0619 |
| 4 | 146.0459 |
| 4 | 146.0655 |
| 4 | 154.0625 |
| 4 | 164.0719 |
| 4 | 179.0562 |
| 4 | 180.0595 |
| 4 | 194.991 |
| 4 | 206.9467 |
| 4 | 209.0667 |
| 4 | 210.9652 |
| 4 | 215.0328 |
| 4 | 216.0362 |
| 4 | 217.0296 |
| 4 | 217.0369 |
| 4 | 218.033 |
| 4 | 225.0617 |
| 4 | 239.0772 |
| 4 | 245.0431 |
| 4 | 259.0224 |
| 4 | 268.1036 |
| 4 | 277.0331 |
| 4 | 284.0988 |
| 4 | 288.9652 |
| 4 | 290.9626 |
| 4 | 294.1195 |

| | |
|---|---|
| 4 | 296.1351 |
| 4 | 298.114 |
| 4 | 306.0768 |
| 4 | 310.151 |
| 4 | 311.1095 |
| 4 | 312.113 |
| 4 | 313.0774 |
| 4 | 314.081 |
| 4 | 323.0285 |
| 4 | 325.1252 |
| 4 | 326.1095 |
| 4 | 326.1284 |
| 4 | 327.0446 |
| 4 | 341.0595 |
| 4 | 344.1348 |
| 4 | 353.0727 |
| 4 | 357.0336 |
| 4 | 359.1194 |
| 4 | 360.1227 |
| 4 | 367.1056 |
| 4 | 375.0537 |
| 4 | 376.057 |
| 4 | 377.0852 |
| 4 | 378.0884 |
| 4 | 379.083 |
| 4 | 379.2336 |
| 4 | 387.1146 |
| 4 | 391.0277 |
| 4 | 392.0314 |
| 4 | 395.0954 |
| 4 | 397.0933 |
| 4 | 438.0812 |
| 4 | 439.0856 |
| 4 | 456.1732 |
| 4 | 469.028 |
| 4 | 473.1623 |
| 4 | 475.1297 |
| 4 | 480.0931 |
| 4 | 487.1774 |
| 4 | 488.1618 |
| 4 | 515.1249 |
| 4 | 521.1722 |
| 4 | 537.1068 |
| 4 | 553.0802 |

| | |
|---|---|
| 4 | 555.1174 |
| 4 | 557.1483 |
| 4 | 571.0918 |
| 4 | 572.0958 |
| 4 | 602.075 |

**Supplemental Table 5**

| Cluster | m/z value |
|---|---|
| 1 | 255.2328 |
| 1 | 256.2361 |
| 1 | 277.2169 |
| 1 | 279.2327 |
| 1 | 280.2364 |
| 1 | 281.2483 |
| 1 | 330.265 |
| 1 | 354.2647 |
| 1 | 490.2809 |
| 1 | 504.2236 |
| 1 | 535.4721 |
| 1 | 559.4728 |
| 1 | 560.4763 |
| 2 | 69.71941 |
| 2 | 73.02988 |
| 2 | 74.02503 |
| 2 | 99.00903 |
| 2 | 103.0039 |
| 2 | 112.9859 |
| 2 | 117.0196 |
| 2 | 120.0305 |
| 2 | 149.0118 |
| 2 | 157.1236 |
| 2 | 162.9823 |
| 2 | 164.0354 |
| 2 | 167.9972 |
| 2 | 168.9893 |
| 2 | 175.0614 |
| 2 | 179.0223 |
| 2 | 189.0688 |
| 2 | 200.1736 |
| 2 | 210.0407 |
| 2 | 239.1288 |
| 2 | 264.1055 |
| 2 | 274.2022 |
| 2 | 293.1759 |
| 2 | 294.1786 |
| 2 | 295.13 |
| 2 | 322.1663 |
| 2 | 341.1083 |
| 2 | 387.1145 |
| 2 | 404.1039 |

| | |
|---|---|
| 2 | 412.9655 |
| 2 | 413.969 |
| 2 | 441.2026 |
| 2 | 459.2951 |
| 2 | 473.2381 |
| 2 | 488.1618 |
| 2 | 696.4683 |
| 2 | 722.5055 |
| 2 | 833.5183 |
| 3 | 59.01413 |
| 3 | 71.01424 |
| 3 | 72.99348 |
| 3 | 87.00905 |
| 3 | 89.02464 |
| 3 | 101.0247 |
| 3 | 104.0357 |
| 3 | 105.0195 |
| 3 | 115.0039 |
| 3 | 119.0352 |
| 3 | 131.0353 |
| 3 | 132.0304 |
| 3 | 133.0508 |
| 3 | 145.0144 |
| 3 | 146.0459 |
| 3 | 147.0301 |
| 3 | 149.0455 |
| 3 | 156.0666 |
| 3 | 160.0617 |
| 3 | 161.0456 |
| 3 | 163.0613 |
| 3 | 164.6301 |
| 3 | 165.0406 |
| 3 | 179.0562 |
| 3 | 194.0457 |
| 3 | 195.0512 |
| 3 | 226.9959 |
| 3 | 242.0514 |
| 3 | 269.0878 |
| 3 | 297.0824 |
| 3 | 313.1141 |
| 3 | 323.0285 |
| 3 | 390.1035 |
| 3 | 435.2962 |
| 3 | 593.0298 |

| | |
|---|---|
| 3 | 671.4652 |
| 3 | 672.4684 |
| 3 | 695.4662 |
| 3 | 721.5016 |
| 3 | 745.5009 |
| 4 | 85.02973 |
| 4 | 96.96993 |
| 4 | 113.0358 |
| 4 | 114.0199 |
| 4 | 114.0562 |
| 4 | 116.0073 |
| 4 | 118.0512 |
| 4 | 129.0195 |
| 4 | 131.0463 |
| 4 | 133.0143 |
| 4 | 134.0177 |
| 4 | 135.0186 |
| 4 | 135.0301 |
| 4 | 145.0619 |
| 4 | 146.0652 |
| 4 | 157.0369 |
| 4 | 173.0092 |
| 4 | 174.0123 |
| 4 | 190.9728 |
| 4 | 191.0561 |
| 4 | 192.0595 |
| 4 | 194.991 |
| 4 | 206.9467 |
| 4 | 209.0667 |
| 4 | 215.0328 |
| 4 | 216.0362 |
| 4 | 217.0296 |
| 4 | 217.0369 |
| 4 | 218.033 |
| 4 | 225.0617 |
| 4 | 245.0431 |
| 4 | 259.0224 |
| 4 | 267.0719 |
| 4 | 268.1036 |
| 4 | 277.0331 |
| 4 | 284.0988 |
| 4 | 287.0498 |
| 4 | 289.0171 |
| 4 | 293.0991 |

| | |
|---|---|
| 4 | 294.0291 |
| 4 | 294.1195 |
| 4 | 296.0985 |
| 4 | 296.1351 |
| 4 | 298.114 |
| 4 | 299.0973 |
| 4 | 301.0646 |
| 4 | 304.9916 |
| 4 | 306.0758 |
| 4 | 307.114 |
| 4 | 310.151 |
| 4 | 311.1095 |
| 4 | 312.113 |
| 4 | 312.1297 |
| 4 | 313.0774 |
| 4 | 314.0811 |
| 4 | 321.0628 |
| 4 | 325.0411 |
| 4 | 325.1252 |
| 4 | 326.1095 |
| 4 | 326.1284 |
| 4 | 327.0446 |
| 4 | 329.1084 |
| 4 | 333.0585 |
| 4 | 336.1295 |
| 4 | 341.0595 |
| 4 | 346.0549 |
| 4 | 353.0727 |
| 4 | 357.0336 |
| 4 | 359.1194 |
| 4 | 367.1056 |
| 4 | 371.1192 |
| 4 | 372.1218 |
| 4 | 375.0537 |
| 4 | 377.0852 |
| 4 | 378.0884 |
| 4 | 379.083 |
| 4 | 387.0096 |
| 4 | 390.1166 |
| 4 | 391.0277 |
| 4 | 395.0954 |
| 4 | 397.0934 |
| 4 | 439.0856 |
| 4 | 456.1713 |

| | |
|---|---|
| 4 | 473.1623 |
| 4 | 475.1296 |
| 4 | 480.0931 |
| 4 | 487.1774 |
| 4 | 503.0922 |
| 4 | 515.1249 |
| 4 | 521.1722 |
| 4 | 526.1191 |
| 4 | 531.0974 |
| 4 | 533.1722 |
| 4 | 537.1068 |
| 4 | 555.1174 |
| 4 | 571.0919 |
| 4 | 746.5053 |
| 4 | 747.5077 |
| 4 | 851.529 |
| 4 | 925.5635 |

**Supplemental Table 6**

| Cluster | m/z value |
|---|---|
| 2 | 66.15642 |
| 2 | 121.0296 |
| 2 | 143.1077 |
| 2 | 157.1236 |
| 2 | 162.9823 |
| 2 | 167.9972 |
| 2 | 189.0688 |
| 2 | 197.0606 |
| 2 | 199.1702 |
| 2 | 236.0991 |
| 2 | 274.5884 |
| 2 | 277.6986 |
| 2 | 293.1759 |
| 2 | 294.111 |
| 2 | 321.3575 |
| 2 | 322.1652 |
| 2 | 349.1527 |
| 2 | 382.4422 |
| 3 | 59.01413 |
| 3 | 61.98868 |
| 3 | 71.01412 |
| 3 | 72.99335 |
| 3 | 75.009 |
| 3 | 76.98836 |
| 3 | 85.02974 |
| 3 | 87.00905 |
| 3 | 89.02464 |
| 3 | 99.00904 |
| 3 | 101.0247 |
| 3 | 102.0563 |
| 3 | 103.0039 |
| 3 | 103.0403 |
| 3 | 105.0195 |
| 3 | 111.009 |
| 3 | 112.9881 |
| 3 | 115.0039 |
| 3 | 115.0402 |
| 3 | 115.0766 |
| 3 | 119.0352 |
| 3 | 119.9471 |
| 3 | 128.0355 |
| 3 | 129.0195 |

| | |
|---|---|
| 3 | 129.0558 |
| 3 | 131.035 |
| 3 | 132.0304 |
| 3 | 133.0143 |
| 3 | 133.0508 |
| 3 | 134.0177 |
| 3 | 135.0186 |
| 3 | 135.0299 |
| 3 | 138.0196 |
| 3 | 142.9985 |
| 3 | 145.0144 |
| 3 | 146.0459 |
| 3 | 147.0301 |
| 3 | 147.0493 |
| 3 | 149.0455 |
| 3 | 156.0666 |
| 3 | 160.0617 |
| 3 | 161.0456 |
| 3 | 163.0613 |
| 3 | 165.0406 |
| 3 | 168.9893 |
| 3 | 173.0092 |
| 3 | 174.0123 |
| 3 | 175.0609 |
| 3 | 179.0562 |
| 3 | 191.0561 |
| 3 | 195.0506 |
| 3 | 203.084 |
| 3 | 210.9646 |
| 3 | 218.1395 |
| 3 | 226.9959 |
| 3 | 228.9932 |
| 3 | 237.0613 |
| 3 | 239.0437 |
| 3 | 239.1288 |
| 3 | 242.0514 |
| 3 | 245.0431 |
| 3 | 275.1246 |
| 3 | 285.0826 |
| 3 | 294.1786 |
| 3 | 297.0824 |
| 3 | 299.0983 |
| 3 | 312.0932 |
| 3 | 313.113 |

| | |
|---|---|
| 3 | 314.08 |
| 3 | 323.0285 |
| 3 | 325.0411 |
| 3 | 326.1084 |
| 3 | 329.1084 |
| 3 | 336.1296 |
| 3 | 346.055 |
| 3 | 353.0714 |
| 3 | 367.1043 |
| 3 | 371.1193 |
| 3 | 379.1639 |
| 3 | 383.0828 |
| 3 | 390.1036 |
| 3 | 406.0392 |
| 3 | 413.969 |
| 3 | 435.2962 |
| 3 | 439.0856 |
| 3 | 448.9857 |
| 3 | 459.2951 |
| 3 | 473.2381 |
| 3 | 586.101 |
| 3 | 593.0298 |
| 3 | 602.075 |
| 3 | 671.4652 |
| 3 | 672.4684 |
| 3 | 695.4662 |
| 3 | 696.4683 |
| 3 | 721.5017 |
| 3 | 745.5009 |
| 3 | 746.5052 |
| 3 | 833.5183 |
| 4 | 88.04062 |
| 4 | 92.92824 |
| 4 | 96.96038 |
| 4 | 104.0355 |
| 4 | 116.0073 |
| 4 | 116.072 |
| 4 | 118.0512 |
| 4 | 131.0463 |
| 4 | 145.0619 |
| 4 | 146.0652 |
| 4 | 154.0621 |
| 4 | 168.0066 |
| 4 | 175.0474 |

| | |
|---|---|
| 4 | 180.0595 |
| 4 | 190.9728 |
| 4 | 193.9737 |
| 4 | 194.991 |
| 4 | 215.0328 |
| 4 | 216.0362 |
| 4 | 217.0296 |
| 4 | 218.033 |
| 4 | 225.0611 |
| 4 | 239.0772 |
| 4 | 243.0545 |
| 4 | 259.0224 |
| 4 | 268.1035 |
| 4 | 269.087 |
| 4 | 277.0331 |
| 4 | 284.0979 |
| 4 | 296.1351 |
| 4 | 298.114 |
| 4 | 301.0646 |
| 4 | 306.0758 |
| 4 | 310.15 |
| 4 | 311.1095 |
| 4 | 313.0775 |
| 4 | 314.0968 |
| 4 | 325.1252 |
| 4 | 326.1284 |
| 4 | 327.0446 |
| 4 | 333.0584 |
| 4 | 334.1251 |
| 4 | 341.0595 |
| 4 | 344.1348 |
| 4 | 357.0336 |
| 4 | 359.1194 |
| 4 | 363.0417 |
| 4 | 375.0537 |
| 4 | 376.057 |
| 4 | 379.2336 |
| 4 | 391.0277 |
| 4 | 392.0314 |
| 4 | 395.0954 |
| 4 | 397.0933 |
| 4 | 456.1714 |
| 4 | 480.0931 |
| 4 | 503.0922 |

| | |
|---|---|
| 4 | 526.1191 |
| 4 | 543.1041 |
| 4 | 555.1174 |
| 4 | 556.1215 |
| 4 | 559.0781 |
| 4 | 563.3339 |
| 4 | 571.0918 |
| 4 | 579.3297 |
| 4 | 684.3951 |
| 1 | 73.02976 |
| 1 | 117.0196 |
| 1 | 118.0228 |
| 1 | 138.0561 |
| 1 | 149.0118 |
| 1 | 157.0507 |
| 1 | 164.0354 |
| 1 | 165.039 |
| 1 | 179.0223 |
| 1 | 194.0457 |
| 1 | 210.0407 |
| 1 | 240.0512 |
| 1 | 246.0175 |
| 1 | 254.0151 |
| 1 | 255.2328 |
| 1 | 279.2327 |
| 1 | 280.2355 |
| 1 | 281.2483 |
| 1 | 309.0461 |
| 1 | 330.265 |
| 1 | 341.1083 |
| 1 | 354.2647 |
| 1 | 377.0838 |
| 1 | 377.0852 |
| 1 | 379.0816 |
| 1 | 387.1146 |
| 1 | 404.1038 |
| 1 | 421.0886 |
| 1 | 451.0988 |
| 1 | 471.2593 |
| 1 | 487.1774 |
| 1 | 488.1618 |
| 1 | 490.2809 |
| 1 | 504.2236 |
| 1 | 515.1249 |

1 552.1553

**Supplemental Table 7**

| bimodal | cortex | vasculature |
|---|---|---|
| 129.0195 | 88.0408 | 215.0328 |
| 145.0144 | 104.0357 | 216.0362 |
| 341.1083 | 118.0512 | 217.0296 |
| 377.0852 | 131.0463 | 217.0369 |
| 378.0884 | 145.0619 | 288.9652 |
| 379.083 | 146.0459 | 290.9626 |
| 387.1146 | 210.9652 | 377.0852 |
| 390.1035 | 268.1035 | 378.0884 |
| 404.1039 | 298.114 | 379.083 |
| 488.1618 | 299.0973 | 395.0954 |
| 515.1249 | 311.1095 | 397.0933 |
| 537.1068 | 325.1252 | 438.0812 |
| 553.0802 | 326.1095 | 469.028 |
| 695.4662 | 326.1284 | |
| 721.5016 | 341.0595 | |
| 745.5009 | 346.0549 | |
| 746.5052 | 357.0336 | |
| 834.519 | 375.0536 | |
| | 391.0277 | |

key:

localization pattern conserved in..

| 1 rep | 2 reps | 3 reps |
|---|---|---|

**Supplemental Table 8**

| B73 unique | OG unique | conserved between OG and B73 |
|---|---|---|
| 59.014156 | 110.976145 | 85.029788 |
| 61.988682 | 114.93723 | 89.024696 |
| 66.156418 | 115.021433 | 96.960381 |
| 69.719414 | 116.00892 | 111.009 |
| 71.014198 | 120.010155 | 115.00388 |
| 72.993434 | 122.02525 | 115.07657 |
| 73.029841 | 122.02641 | 116.007267 |
| 74.025032 | 124.008282 | 117.019547 |
| 74.04068 | 127.05257 | 118.02284 |
| 75.009003 | 127.07706 | 118.99276 |
| 76.988358 | 130.023023 | 119.035177 |
| 87.009048 | 131.02367 | 121.029635 |
| 88.040699 | 133.05216 | 128.03546 |
| 92.928238 | 134.02486 | 129.019513 |
| 96.969929 | 135.97268 | 129.05579 |
| 99.009036 | 136.005243 | 131.03517 |
| 101.02468 | 137.02531 | 131.04631 |
| 102.05634 | 138.05714 | 132.03036 |
| 103.0039 | 139.006905 | 133.01437 |
| 103.04025 | 142.05179 | 134.017687 |
| 104.03559 | 142.999597 | 135.01859 |
| 105.01953 | 145.051576 | 135.030043 |
| 112.98586 | 146.96708 | 138.01964 |
| 112.98824 | 149.047163 | 145.0144 |
| 113.02465 | 150.016118 | 145.061907 |
| 113.03577 | 150.01868 | 146.04585 |
| 114.0199 | 153.023447 | 147.03009 |
| 114.05623 | 154.02658 | 149.01183 |
| 115.04022 | 155.01917 | 154.0623 |
| 116.03559 | 155.02745 | 157.05067 |
| 116.07195 | 158.01443 | 157.12357 |
| 118.05118 | 159.011033 | 162.98241 |
| 119.94714 | 160.063113 | 164.03535 |
| 120.03054 | 161.046673 | 165.04059 |
| 133.05077 | 163.0627 | 167.997157 |
| 138.05608 | 163.999925 | 168.98931 |
| 142.99854 | 165.00386 | 173.00922 |
| 143.10774 | 166.05244 | 174.012477 |

| | | |
|---|---|---|
| 146.06531 | 169.99364 | 175.061223 |
| 147.0493 | 171.066765 | 179.02225 |
| 149.0456 | 171.103155 | 179.05618 |
| 156.06663 | 171.13968 | 191.056083 |
| 157.03694 | 173.02641 | 192.05945 |
| 160.06165 | 173.05708 | 194.045677 |
| 161.04558 | 173.08241 | 194.990967 |
| 163.06129 | 175.01445 | 195.050983 |
| 164.07187 | 175.025972 | 199.1702 |
| 164.63013 | 177.07782 | 210.040643 |
| 165.03899 | 177.09294 | 210.96492 |
| 168.00664 | 178.01495 | 215.03282 |
| 175.04738 | 178.01596 | 216.036167 |
| 177.0768 | 178.01698 | 217.02963 |
| 180.05948 | 179.023423 | 218.03302 |
| 189.0688 | 179.02512 | 225.06149 |
| 190.97281 | 179.05686 | 226.996163 |
| 193.97374 | 179.05791 | 228.99319 |
| 197.06058 | 179.05893 | 237.06125 |
| 199.17603 | 183.013585 | 239.07717 |
| 200.17355 | 183.01532 | 242.05142 |
| 203.08429 | 184.98393 | 254.015085 |
| 206.94672 | 184.985 | 255.232807 |
| 209.06673 | 186.04535 | 256.23608 |
| 217.03689 | 186.04647 | 259.02243 |
| 218.13947 | 187.04157 | 267.07187 |
| 236.09914 | 187.042643 | 269.087435 |
| 239.04367 | 187.04375 | 277.03306 |
| 239.12883 | 187.04483 | 277.21689 |
| 240.05123 | 187.09737 | 279.23268 |
| 243.05446 | 187.09847 | 280.236123 |
| 245.04312 | 188.04465 | 289.01712 |
| 246.01746 | 188.04575 | 306.076133 |
| 264.1055 | 188.04686 | 321.06281 |
| 268.10354 | 188.05789 | 333.05844 |
| 274.20215 | 188.05901 | 341.10829 |
| 274.58844 | 188.062015 | 367.10556 |
| 275.12457 | 189.00468 | 376.05704 |
| 277.69866 | 189.05806 | 378.08838 |
| 281.24826 | 191.019225 | 379.083025 |
| 284.0798 | 191.02035 | 387.11455 |
| 284.09849 | 191.05537 | 402.10309 |

| | | |
|---|---|---|
| 285.08255 | 191.05764 | 404.10385 |
| 287.04984 | 191.05878 | 439.08558 |
| 288.96521 | 192.02379 | 473.16229 |
| 290.96259 | 192.06021 | 533.17218 |
| 293.09912 | 193.073875 | 579.32965 |
| 293.17589 | 194.04642 | 671.46517 |
| 294.02905 | 194.04758 | 695.46619 |
| 294.11099 | 194.04991 | 696.46829 |
| 294.11954 | 195.04938 | 745.50083 |
| 294.17861 | 195.05172 | 746.50523 |
| 295.12997 | 195.05314 | 747.50769 |
| 296.09851 | 196.0631 | |
| 296.13507 | 196.98409 | |
| 297.08243 | 196.98524 | |
| 298.11401 | 197.029 | |
| 299.09729 | 197.030195 | |
| 299.0983 | 197.11896 | |
| 301.0646 | 199.0065 | |
| 304.99158 | 199.00771 | |
| 307.11395 | 199.09781 | |
| 309.04611 | 200.05666 | |
| 310.14999 | 201.057825 | |
| 310.15103 | 201.11397 | |
| 311.10952 | 202.01616 | |
| 312.0932 | 202.01741 | |
| 312.11301 | 203.037027 | |
| 312.12967 | 203.10515 | |
| 313.07742 | 203.10637 | |
| 313.11301 | 204.08812 | |
| 313.11406 | 204.08939 | |
| 314.07996 | 204.09064 | |
| 314.08104 | 205.053155 | |
| 314.0968 | 208.9864 | |
| 321.35748 | 209.06789 | |
| 322.16516 | 209.06917 | |
| 322.16626 | 210.04303 | |
| 323.02852 | 211.04477 | |
| 325.0411 | 211.04608 | |
| 325.1252 | 211.13399 | |
| 326.1084 | 211.135305 | |
| 326.1095 | 213.02327 | |
| 326.12839 | 213.11375 | |

| | |
|---|---|
| 327.04459 | 213.18698 |
| 329.1084 | 213.18831 |
| 330.26494 | 214.03503 |
| 334.12512 | 214.03636 |
| 336.12953 | 214.04979 |
| 341.05951 | 214.05113 |
| 344.13483 | 215.033793 |
| 346.05495 | 215.03514 |
| 349.15265 | 216.03545 |
| 349.1539 | 216.05716 |
| 353.07141 | 217.01639 |
| 353.07269 | 217.030473 |
| 354.26471 | 217.03185 |
| 357.0336 | 217.052335 |
| 359.11938 | 217.053725 |
| 360.12268 | 218.10399 |
| 363.04166 | 218.10538 |
| 367.10425 | 220.7668 |
| 368.97638 | 222.765283 |
| 371.11923 | 223.02916 |
| 372.12177 | 224.05574 |
| 375.05365 | 224.05717 |
| 377.0838 | 224.76329 |
| 377.08517 | 225.08794 |
| 379.08163 | 225.08943 |
| 379.16394 | 225.09087 |
| 379.23361 | 225.18771 |
| 382.44217 | 226.99828 |
| 383.08279 | 227.03922 |
| 387.00955 | 227.129645 |
| 390.10354 | 227.20087 |
| 390.11661 | 227.203538 |
| 391.02774 | 228.05296 |
| 392.03139 | 229.05301 |
| 395.09543 | 230.035878 |
| 397.09333 | 231.03215 |
| 406.03918 | 231.03368 |
| 412.96552 | 232.03437 |
| 413.96898 | 232.035865 |
| 421.08862 | 232.05099 |
| 435.29622 | 233.04768 |
| 438.08121 | 233.04919 |

| | |
|---|---|
| 441.20261 | 233.15581 |
| 448.98566 | 234.05081 |
| 451.09884 | 236.0052 |
| 456.17135 | 236.09518 |
| 456.17316 | 236.09675 |
| 459.29507 | 236.98848 |
| 459.29691 | 239.05963 |
| 469.02795 | 241.05441 |
| 471.25928 | 241.2175 |
| 473.2381 | 241.21912 |
| 475.12966 | 242.05304 |
| 480.09314 | 243.06904 |
| 487.17742 | 243.10487 |
| 488.1618 | 244.04675 |
| 490.28091 | 245.044455 |
| 503.09221 | 246.046715 |
| 504.22359 | 247.02736 |
| 515.1249 | 247.06227 |
| 521.17224 | 247.063895 |
| 526.11908 | 248.05867 |
| 531.09735 | 248.08118 |
| 535.47205 | 253.091413 |
| 537.10675 | 253.21681 |
| 543.10406 | 253.218303 |
| 552.15533 | 253.22029 |
| 553.0802 | 254.01677 |
| 555.11737 | 255.07 |
| 556.12152 | 255.23178 |
| 557.14832 | 255.23519 |
| 559.07813 | 255.23692 |
| 559.47278 | 256.23764 |
| 560.47626 | 257.0802 |
| 563.33392 | 257.24069 |
| 571.09182 | 258.98267 |
| 572.09576 | 259.063355 |
| 586.10095 | 260.04678 |
| 593.02979 | 260.98032 |
| 602.07495 | 261.04166 |
| 672.46844 | 261.04346 |
| 684.39508 | 262.06219 |
| 721.50161 | 263.057695 |
| 722.50549 | 263.05954 |

| | |
|---|---|
| 833.51828 | 265.14714 |
| 851.52899 | 265.14904 |
| 925.56348 | 265.15126 |
| 982.09424 | 266.15216 |
| | 266.15472 |
| | 267.10754 |
| | 267.23343 |
| | 269.08606 |
| | 269.12152 |
| | 269.12341 |
| | 269.212175 |
| | 269.24937 |
| | 270.94568 |
| | 271.22827 |
| | 272.94272 |
| | 272.94467 |
| | 274.002593 |
| | 274.02423 |
| | 274.02618 |
| | 275.020465 |
| | 275.02237 |
| | 275.057515 |
| | 275.05948 |
| | 276.02411 |
| | 276.02606 |
| | 276.07608 |
| | 276.07715 |
| | 276.078705 |
| | 277.07147 |
| | 277.072863 |
| | 277.074645 |
| | 277.07669 |
| | 277.218765 |
| | 278.07642 |
| | 278.07767 |
| | 278.07895 |
| | 278.22055 |
| | 278.22256 |
| | 279.07816 |
| | 279.07938 |
| | 279.23387 |
| | 279.23546 |

| |
|---|
| 279.23712 |
| 280.23849 |
| 280.98251 |
| 280.98456 |
| 281.12158 |
| 281.12283 |
| 281.24725 |
| 281.249474 |
| 281.25238 |
| 282.25107 |
| 282.25211 |
| 282.25418 |
| 283.10086 |
| 283.10193 |
| 283.136645 |
| 283.138685 |
| 283.263105 |
| 283.26521 |
| 283.266965 |
| 284.14117 |
| 284.267 |
| 284.268533 |
| 284.270755 |
| 288.04184 |
| 289.036926 |
| 289.03989 |
| 289.073465 |
| 290.04095 |
| 290.05573 |
| 290.057085 |
| 291.053347 |
| 291.089363 |
| 292.072095 |
| 293.068957 |
| 293.122375 |
| 293.17795 |
| 293.17917 |
| 293.18304 |
| 293.21308 |
| 293.21451 |
| 294.18417 |
| 294.21655 |

294.21793
295.13623
295.13843
295.22731
295.229535
295.23169
296.14218
296.23188
296.23364
297.15297
297.1572
298.15683
298.15906
299.15082
299.153015
300.15097
300.152725
300.15442
302.069033
302.07095
305.03241
305.06888
305.9743
306.07892
306.08121
306.088367
307.08237
307.085003
307.13766
307.19525
308.0885
309.11819
309.1532
309.17642
309.20892
309.28101
309.28217
310.1554
310.15775
311.14145
311.14282
311.16742

311.16974
311.22372
311.22604
312.16998
312.17239
312.17474
312.228605
313.14667
313.16568
313.16705
313.169385
314.162215
314.166975
314.168803
314.1709
315.16898
315.17142
319.048187
319.12161
320.065
320.068053
321.06171
321.06415
321.06683
321.06784
321.15289
321.154175
321.2121
322.068187
322.14502
323.02981
323.03232
323.06752
323.06952
323.13297
323.13416
323.13617
323.13916
323.16898
324.13782
324.17029
324.17276

325.15622
325.15778
325.18396
325.18643
325.18893
326.15933
326.16058
326.18454
326.18713
326.18964
326.19217
327.17465
327.17633
327.18378
327.21899
327.22028
328.17694
328.17944
328.183075
329.14294
329.14413
329.18637
329.23499
330.04739
332.030475
332.07965
333.02655
333.02917
333.060315
333.06149
333.06296
333.09933
333.10056
333.13701
335.07944
335.08072
335.11615
335.16867
337.09412
337.09573
337.183155
337.185805

| |
|---|
| 337.20703 |
| 337.20969 |
| 338.18652 |
| 338.18771 |
| 338.18918 |
| 339.16504 |
| 339.168745 |
| 339.17139 |
| 339.19705 |
| 339.19974 |
| 339.20087 |
| 339.20355 |
| 339.2052 |
| 339.32819 |
| 339.32968 |
| 340.204115 |
| 341.08093 |
| 341.11062 |
| 341.11337 |
| 341.14337 |
| 341.17801 |
| 341.19147 |
| 341.19308 |
| 342.112625 |
| 342.11542 |
| 342.11652 |
| 342.18173 |
| 342.19217 |
| 342.19931 |
| 342.20108 |
| 343.15991 |
| 344.16281 |
| 346.05744 |
| 346.06076 |
| 346.976297 |
| 346.9792 |
| 349.05978 |
| 349.09604 |
| 351.07526 |
| 351.08276 |
| 351.08563 |
| 351.1109 |

351.11197
351.20102
352.09329
353.08932
353.09219
353.178425
353.18015
353.20294
353.214215
353.217085
354.22046
355.15891
364.108995
365.08878
365.09012
365.13953
365.14072
365.14252
366.108655
366.1109
367.102765
367.10852
367.10968
367.229845
367.23187
367.3595
367.36139
368.109343
368.11072
368.23636
369.109545
369.11151
369.117595
369.11923
369.12671
369.127765
375.18591
377.053205
377.05505
377.086393
377.08963
378.09085

| |
|---|
| 378.09195 |
| 378.09396 |
| 378.10757 |
| 378.10962 |
| 378.11078 |
| 379.06863 |
| 379.07071 |
| 379.08661 |
| 379.10347 |
| 379.10547 |
| 379.10669 |
| 379.15381 |
| 379.156333 |
| 380.08624 |
| 380.0914 |
| 380.1084 |
| 380.10992 |
| 381.08569 |
| 381.0867 |
| 382.129853 |
| 382.13272 |
| 382.20618 |
| 382.25098 |
| 382.25308 |
| 383.0611 |
| 383.09958 |
| 383.10068 |
| 383.10275 |
| 383.1434 |
| 385.070235 |
| 387.11356 |
| 391.1405 |
| 392.10196 |
| 392.10318 |
| 392.13626 |
| 392.13846 |
| 392.18842 |
| 393.10358 |
| 393.17029 |
| 393.17236 |
| 393.17361 |
| 395.10165 |

| |
|---|
| 395.12854 |
| 395.26062 |
| 395.26297 |
| 396.14517 |
| 396.14853 |
| 396.26663 |
| 397.14883 |
| 397.149885 |
| 398.10535 |
| 404.105145 |
| 404.10873 |
| 405.10782 |
| 407.15604 |
| 407.15805 |
| 408.097505 |
| 409.145465 |
| 409.14743 |
| 409.27878 |
| 409.28009 |
| 410.1611 |
| 410.163465 |
| 411.16379 |
| 411.16577 |
| 412.121295 |
| 413.07495 |
| 413.07614 |
| 413.078535 |
| 415.07297 |
| 415.07541 |
| 417.10303 |
| 417.11115 |
| 421.0354 |
| 421.03693 |
| 421.225295 |
| 421.22769 |
| 422.03876 |
| 422.03995 |
| 423.03186 |
| 423.034465 |
| 423.16104 |
| 423.23929 |
| 423.25296 |

| |
|---|
| 423.25421 |
| 424.037525 |
| 424.177965 |
| 424.17947 |
| 426.13596 |
| 426.137065 |
| 427.09073 |
| 427.093065 |
| 427.28714 |
| 428.94687 |
| 429.08987 |
| 429.09128 |
| 431.11746 |
| 431.12326 |
| 431.12518 |
| 431.12711 |
| 431.14441 |
| 433.2352 |
| 433.23636 |
| 433.23904 |
| 434.23987 |
| 437.09512 |
| 437.09665 |
| 437.09906 |
| 439.08224 |
| 439.08389 |
| 439.127155 |
| 439.12961 |
| 444.91925 |
| 445.13211 |
| 445.13339 |
| 445.13611 |
| 445.14011 |
| 445.14209 |
| 445.14413 |
| 447.24994 |
| 447.25287 |
| 448.253955 |
| 448.256075 |
| 450.29074 |
| 450.29208 |
| 451.287185 |

| |
|---|
| 451.28979 |
| 451.32672 |
| 452.27994 |
| 452.29214 |
| 461.15137 |
| 461.2662 |
| 461.26901 |
| 463.24747 |
| 465.30585 |
| 465.30835 |
| 465.340195 |
| 465.3432 |
| 467.17062 |
| 467.23108 |
| 471.276505 |
| 471.27945 |
| 473.28156 |
| 473.28317 |
| 473.28604 |
| 476.27759 |
| 476.279947 |
| 476.28214 |
| 479.20343 |
| 479.2063 |
| 479.20789 |
| 479.21945 |
| 479.22134 |
| 479.35605 |
| 479.35797 |
| 480.21017 |
| 482.12003 |
| 483.117875 |
| 483.27237 |
| 483.27371 |
| 483.27518 |
| 483.27695 |
| 484.11877 |
| 484.12177 |
| 484.27502 |
| 484.27836 |
| 487.178455 |
| 487.18169 |

487.18326
494.27911
494.28131
494.28256
495.13922
495.19852
495.200485
495.20352
495.27591
495.27908
496.14285
496.14468
496.20273
496.20612
496.28177
501.1955
503.19678
507.16821
507.27264
507.27423
507.27582
508.27603
511.19397
511.19545
515.26569
515.26904
519.23444
519.23596
519.2395
521.18347
527.24377
528.16174
528.24432
528.24628
528.24807
528.24957
529.14764
529.15656
529.15808
529.16174
529.2395
529.241485

| |
|---|
| 529.24329 |
| 529.24432 |
| 529.24677 |
| 529.28314 |
| 530.16089 |
| 530.16449 |
| 535.22925 |
| 535.23102 |
| 535.23285 |
| 535.23456 |
| 545.1214 |
| 551.22656 |
| 554.25824 |
| 554.26367 |
| 555.15594 |
| 555.26404 |
| 555.26788 |
| 559.13892 |
| 563.33643 |
| 564.33508 |
| 564.33875 |
| 564.34085 |
| 571.19067 |
| 571.19281 |
| 572.192595 |
| 573.18726 |
| 573.18927 |
| 573.19312 |
| 574.195925 |
| 577.35431 |
| 580.33307 |
| 580.335235 |
| 587.02881 |
| 587.29407 |
| 587.30103 |
| 587.32098 |
| 587.33313 |
| 587.335695 |
| 588.33911 |
| 589.35382 |
| 603.34723 |
| 603.35101 |

| |
|---|
| 604.33496 |
| 606.37579 |
| 606.38013 |
| 617.29199 |
| 619.35999 |
| 619.36221 |
| 622.37109 |
| 627.36444 |
| 627.36554 |
| 627.3692 |
| 631.30499 |
| 631.30664 |
| 631.308655 |
| 631.31061 |
| 631.31342 |
| 632.3121 |
| 632.31467 |
| 633.30634 |
| 634.37427 |
| 637.36865 |
| 637.371155 |
| 637.37555 |
| 638.37482 |
| 638.37775 |
| 639.26587 |
| 639.26837 |
| 639.27002 |
| 639.27274 |
| 640.26959 |
| 641.27216 |
| 641.2757 |
| 643.36148 |
| 643.36487 |
| 645.32404 |
| 645.32593 |
| 645.327915 |
| 645.33301 |
| 646.32489 |
| 646.32797 |
| 646.32996 |
| 646.33191 |
| 647.28497 |

| |
|---|
| 647.28912 |
| 647.30603 |
| 650.36749 |
| 653.27881 |
| 653.28156 |
| 653.28424 |
| 653.28604 |
| 653.36481 |
| 653.36981 |
| 654.28308 |
| 654.28735 |
| 654.29047 |
| 655.28369 |
| 655.28735 |
| 655.29083 |
| 656.28943 |
| 659.33502 |
| 659.33704 |
| 659.34235 |
| 659.34491 |
| 660.3399 |
| 660.34299 |
| 660.34497 |
| 660.34723 |
| 661.29935 |
| 661.30219 |
| 661.31665 |
| 662.3045 |
| 663.29999 |
| 667.29779 |
| 667.30072 |
| 667.303955 |
| 668.30475 |
| 669.30359 |
| 671.33032 |
| 672.46472 |
| 672.470245 |
| 672.47229 |
| 673.35333 |
| 673.35876 |
| 673.47095 |
| 673.47327 |

673.47839
673.48096
674.3584
674.36151
675.3139
675.31854
676.31348
676.31622
676.32098
677.4057
681.31439
681.3194
682.31635
682.31964
685.47943
685.48529
686.47852
686.48077
686.48322
686.48444
686.48624
687.45959
688.46802
689.32965
689.33426
695.46277
695.467865
696.46558
696.47345
697.47308
697.47534
697.47778
697.48093
698.48276
698.48785
699.49463
700.50085
703.45502
704.45929
709.48505
710.48468
710.48853

| |
|---|
| 711.46173 |
| 712.42871 |
| 712.46942 |
| 713.47931 |
| 714.50754 |
| 714.51282 |
| 715.58728 |
| 716.59555 |
| 716.59845 |
| 717.59775 |
| 717.60065 |
| 717.60236 |
| 718.60205 |
| 721.49731 |
| 721.5032 |
| 721.50555 |
| 722.50153 |
| 722.50739 |
| 722.51001 |
| 723.50537 |
| 723.50903 |
| 725.388 |
| 725.39044 |
| 726.4447 |
| 727.45081 |
| 727.45441 |
| 727.45923 |
| 728.45905 |
| 729.4693 |
| 732.513 |
| 734.513 |
| 738.50311 |
| 738.5058 |
| 738.50916 |
| 739.50861 |
| 741.38354 |
| 741.38757 |
| 742.50281 |
| 743.48712 |
| 743.615205 |
| 743.62061 |
| 744.48877 |

| |
|---|
| 744.61914 |
| 745.50323 |
| 745.50616 |
| 746.50018 |
| 746.50275 |
| 746.50891 |
| 747.51334 |
| 747.51532 |
| 748.51099 |
| 748.51373 |
| 748.51489 |
| 748.51965 |
| 749.52814 |
| 760.54419 |
| 760.55072 |
| 761.49847 |
| 761.52063 |
| 761.52472 |
| 762.5014 |
| 762.54648 |
| 763.54736 |
| 763.55231 |
| 767.53424 |
| 769.50272 |
| 770.50342 |
| 775.53738 |
| 776.53906 |
| 776.54089 |
| 776.54535 |
| 777.49042 |
| 777.49609 |
| 778.49298 |
| 787.54712 |
| 789.422455 |
| 789.42596 |
| 790.4292 |
| 791.53302 |
| 792.46143 |
| 793.51837 |
| 803.44031 |
| 803.44238 |
| 803.44586 |

| |
|---|
| 803.566955 |
| 804.4447 |
| 804.45044 |
| 804.57159 |
| 804.57471 |
| 804.578 |
| 805.57806 |
| 806.57672 |
| 817.45477 |
| 817.46063 |
| 818.46149 |
| 818.46552 |
| 820.49268 |
| 822.492 |
| 822.49695 |
| 831.59283 |
| 831.59668 |
| 832.59915 |
| 833.51453 |
| 834.51404 |
| 834.52075 |
| 834.52429 |
| 835.52557 |
| 835.53021 |
| 848.52759 |
| 849.52777 |
| 850.51886 |
| 850.52594 |
| 851.52786 |

**Supplemental Table 9**

| OG Root 1 | OG Root 2 | OG Root 3 |
|---|---|---|
| 110.9759 | 111.0089 | 85.02996 |
| 110.9764 | 115.0037 | 85.03028 |
| 111.0089 | 115.0042 | 85.03062 |
| 111.0094 | 115.0047 | 89.02503 |
| 115.0037 | 115.0211 | 89.02574 |
| 115.0042 | 115.0216 | 96.96066 |
| 115.0216 | 115.0776 | 96.96107 |
| 115.077 | 117.0193 | 114.9372 |
| 116.0073 | 117.0198 | 115.0042 |
| 116.0079 | 118.9928 | 115.0047 |
| 117.0193 | 119.0356 | 115.0053 |
| 117.0198 | 122.0253 | 116.0079 |
| 118.0226 | 124.0079 | 116.0089 |
| 118.9928 | 129.0196 | 117.0204 |
| 119.035 | 129.0202 | 117.0209 |
| 119.0356 | 130.0228 | 119.0356 |
| 121.0298 | 131.0461 | 119.0361 |
| 122.0253 | 132.0303 | 119.0367 |
| 124.0079 | 132.0309 | 120.0099 |
| 124.0085 | 133.0144 | 120.0104 |
| 127.0771 | 133.0151 | 121.0304 |
| 128.0354 | 133.0157 | 122.0264 |
| 129.019 | 134.0176 | 124.0079 |
| 129.0196 | 134.0182 | 124.0085 |
| 129.0202 | 135.0187 | 124.0091 |
| 130.0228 | 138.0194 | 127.0526 |
| 130.0235 | 138.0201 | 128.036 |
| 131.0237 | 139.0069 | 128.0366 |
| 131.0346 | 142.9988 | 128.0372 |
| 131.0352 | 142.9995 | 129.0202 |
| 131.0461 | 145.0505 | 129.0208 |
| 132.0303 | 145.0617 | 129.0566 |
| 132.0309 | 145.0625 | 131.0359 |
| 133.0138 | 146.0458 | 131.0365 |
| 133.0145 | 146.0466 | 132.0309 |
| 133.0151 | 146.0473 | 132.0316 |
| 134.0175 | 146.9667 | 132.0322 |
| 134.0182 | 147.03 | 133.0151 |
| 134.0249 | 149.0122 | 133.0158 |
| 135.018 | 153.0224 | 133.0518 |
| 135.0187 | 153.0232 | 133.0525 |
| 135.0193 | 153.024 | 134.0182 |

| | | |
|---|---|---|
| 135.0301 | 153.0248 | 134.0189 |
| 138.0201 | 154.0266 | 134.0196 |
| 138.0569 | 154.0626 | 135.0194 |
| 139.0069 | 155.0192 | 135.02 |
| 142.0518 | 157.0511 | 135.0308 |
| 142.9988 | 159.0107 | 135.0314 |
| 142.9995 | 163.9989 | 135.972 |
| 145.0147 | 163.9997 | 135.9727 |
| 145.0513 | 164.0357 | 135.9734 |
| 145.0617 | 165.0406 | 136.0046 |
| 145.0625 | 167.9975 | 136.0053 |
| 146.0458 | 168.9898 | 136.0059 |
| 146.0466 | 169.9936 | 137.0253 |
| 147.03 | 171.1027 | 138.0208 |
| 147.0308 | 173.0094 | 138.0215 |
| 149.0114 | 173.0259 | 138.0569 |
| 149.0121 | 173.0269 | 138.0576 |
| 149.0129 | 173.0571 | 142.9995 |
| 150.0155 | 174.0125 | 143.0002 |
| 153.0225 | 175.0254 | 143.001 |
| 153.0233 | 175.061 | 145.0513 |
| 153.0241 | 175.062 | 145.052 |
| 154.0618 | 178.015 | 145.0528 |
| 154.0626 | 179.022 | 145.0625 |
| 155.0275 | 179.023 | 145.0632 |
| 157.0511 | 179.0559 | 145.064 |
| 157.1235 | 179.0569 | 146.0466 |
| 157.1244 | 183.0136 | 146.0473 |
| 158.0144 | 183.0151 | 146.0481 |
| 159.0108 | 183.0157 | 146.9675 |
| 159.0116 | 186.0465 | 147.0308 |
| 161.0458 | 187.0415 | 147.0315 |
| 162.983 | 187.0426 | 149.0129 |
| 163.9989 | 188.0617 | 149.0137 |
| 163.9998 | 189.0581 | 149.0464 |
| 164.0348 | 191.0192 | 149.0472 |
| 164.0357 | 191.0203 | 149.0479 |
| 164.0366 | 191.0565 | 150.0155 |
| 165.0397 | 192.0602 | 150.0163 |
| 165.0406 | 194.0452 | 150.0171 |
| 165.0415 | 194.0464 | 150.0187 |
| 167.9967 | 194.0476 | 154.0634 |
| 167.9976 | 195.0505 | 154.0638 |
| 167.9985 | 196.9852 | 154.0642 |

| | | |
|---|---|---|
| 171.0663 | 197.029 | 157.1243 |
| 171.0673 | 197.0302 | 157.1248 |
| 171.1027 | 199.0065 | 160.0624 |
| 171.1036 | 199.0077 | 160.0633 |
| 171.1391 | 201.0578 | 160.0637 |
| 171.14 | 205.0532 | 161.0467 |
| 173.0094 | 208.9864 | 161.0476 |
| 173.0104 | 210.0411 | 163.0623 |
| 173.0259 | 210.9648 | 163.0631 |
| 173.0269 | 215.0325 | 163.9998 |
| 173.0824 | 215.0338 | 164.0002 |
| 174.0125 | 217.016 | 164.0006 |
| 174.0135 | 217.0291 | 164.0016 |
| 175.0145 | 217.0305 | 164.0366 |
| 175.0254 | 217.0523 | 164.0375 |
| 175.061 | 218.104 | 165.0034 |
| 175.062 | 222.7644 | 165.0043 |
| 177.0929 | 222.7658 | 165.0406 |
| 179.022 | 223.0292 | 165.0415 |
| 179.023 | 225.0879 | 165.0424 |
| 179.0558 | 226.9968 | 166.052 |
| 179.0569 | 228.9937 | 166.0529 |
| 183.0136 | 229.053 | 171.1036 |
| 183.0151 | 230.0349 | 171.14 |
| 184.9839 | 230.0356 | 173.0104 |
| 184.985 | 230.0364 | 173.0113 |
| 186.0454 | 231.0321 | 175.0254 |
| 186.0465 | 232.0344 | 175.0264 |
| 187.0416 | 232.0359 | 175.0273 |
| 187.0426 | 232.051 | 175.062 |
| 187.0974 | 233.0477 | 175.063 |
| 187.0985 | 236.0052 | 177.0778 |
| 188.0447 | 236.0968 | 178.016 |
| 188.0458 | 236.9885 | 178.017 |
| 188.0623 | 237.0619 | 179.0235 |
| 189.0047 | 239.0763 | 179.0241 |
| 191.0192 | 241.2175 | 179.0251 |
| 191.0204 | 243.0686 | 179.0569 |
| 191.0554 | 243.1046 | 179.0579 |
| 191.0565 | 243.1052 | 179.0589 |
| 191.0576 | 245.043 | 186.0465 |
| 192.0238 | 245.0447 | 187.0427 |
| 192.0591 | 247.0639 | 187.0438 |
| 192.0602 | 253.091 | 187.0448 |

| | | |
|---|---|---|
| 194.0453 | 253.0916 | 187.0985 |
| 194.0464 | 253.2178 | 188.0469 |
| 194.9911 | 254.0151 | 188.0579 |
| 195.0494 | 255.2335 | 188.059 |
| 195.0505 | 255.2352 | 191.0577 |
| 195.0517 | 256.2358 | 191.0588 |
| 196.9841 | 256.2376 | 193.073 |
| 196.9852 | 258.9827 | 193.0736 |
| 197.029 | 259.0629 | 193.0742 |
| 197.0302 | 259.0638 | 193.0747 |
| 197.119 | 260.9803 | 194.0464 |
| 199.0978 | 261.0435 | 194.0476 |
| 199.1711 | 262.0622 | 194.0499 |
| 200.0567 | 263.0577 | 195.0517 |
| 201.0578 | 265.1471 | 195.0529 |
| 201.114 | 265.149 | 195.0534 |
| 202.0162 | 267.1075 | 196.0631 |
| 202.0174 | 267.2334 | 203.1052 |
| 203.0364 | 269.2122 | 203.1064 |
| 203.037 | 269.2489 | 204.0894 |
| 203.0376 | 270.9457 | 204.0906 |
| 204.0881 | 271.2283 | 209.0679 |
| 205.0532 | 272.9447 | 209.0692 |
| 210.0411 | 274.0029 | 210.043 |
| 210.9648 | 274.0262 | 211.1353 |
| 211.0448 | 275.0205 | 213.187 |
| 211.0461 | 275.0224 | 213.1883 |
| 211.134 | 275.0575 | 214.0498 |
| 211.1353 | 276.0241 | 214.0511 |
| 213.0233 | 276.0761 | 215.0338 |
| 213.1138 | 276.0772 | 215.0351 |
| 214.035 | 277.0715 | 217.0305 |
| 214.0364 | 277.0726 | 217.0319 |
| 215.0324 | 277.0746 | 217.0537 |
| 215.0338 | 277.2168 | 218.1054 |
| 216.0355 | 278.0777 | 225.0894 |
| 216.0368 | 279.0782 | 225.0909 |
| 216.0572 | 279.2327 | 225.1877 |
| 217.0168 | 279.2338 | 226.9983 |
| 217.0305 | 280.2365 | 227.2033 |
| 217.0524 | 281.1216 | 227.2039 |
| 217.0537 | 281.1228 | 227.2047 |
| 218.0337 | 281.2495 | 228.0524 |
| 218.104 | 282.2511 | 228.0533 |

| | | |
|---|---|---|
| 220.7668 | 282.2521 | 228.0539 |
| 222.7651 | 283.1366 | 230.0371 |
| 222.7658 | 283.1387 | 231.0337 |
| 224.0557 | 283.2631 | 233.1558 |
| 224.0572 | 283.2651 | 241.2191 |
| 224.7633 | 284.2683 | 245.0437 |
| 225.0605 | 288.0418 | 245.0447 |
| 225.0611 | 289.0365 | 245.0454 |
| 226.9968 | 289.0378 | 245.0463 |
| 227.0392 | 289.0735 | 248.0812 |
| 227.1293 | 290.041 | 253.2186 |
| 227.13 | 290.0557 | 253.2203 |
| 227.2009 | 290.0571 | 255.2352 |
| 227.2024 | 291.0528 | 255.2369 |
| 228.0524 | 291.0536 | 257.0802 |
| 228.9937 | 291.089 | 265.1491 |
| 230.0349 | 291.0898 | 265.1509 |
| 230.0364 | 292.0716 | 265.1516 |
| 231.0322 | 293.0685 | 266.1529 |
| 232.0344 | 293.0692 | 266.1547 |
| 232.0359 | 293.178 | 269.25 |
| 232.051 | 293.2131 | 276.0783 |
| 233.0477 | 295.2273 | 276.0791 |
| 233.0492 | 296.1422 | 277.0747 |
| 234.0508 | 296.2315 | 277.0767 |
| 236.0052 | 297.1529 | 277.2188 |
| 236.0952 | 298.1568 | 278.079 |
| 236.0967 | 299.153 | 278.2226 |
| 237.0619 | 300.151 | 279.2339 |
| 239.0596 | 300.1523 | 279.2351 |
| 241.0541 | 300.1532 | 279.2359 |
| 241.0547 | 302.0687 | 279.2371 |
| 241.2175 | 302.0696 | 280.2385 |
| 242.0514 | 305.0324 | 280.9846 |
| 242.053 | 305.0689 | 281.2483 |
| 243.0695 | 305.9743 | 281.2492 |
| 244.0468 | 306.0766 | 281.2503 |
| 245.0431 | 306.088 | 281.2512 |
| 245.0447 | 307.0847 | 281.2524 |
| 246.0464 | 307.1377 | 282.2542 |
| 246.047 | 308.0885 | 283.2653 |
| 247.0274 | 309.1532 | 283.2666 |
| 247.0623 | 310.1554 | 283.2673 |
| 247.0639 | 310.1578 | 284.2704 |

| | | |
|---|---|---|
| 248.0587 | 311.1415 | 284.2712 |
| 253.091 | 311.1428 | 288.0419 |
| 253.092 | 311.1697 | 289.0399 |
| 253.2168 | 312.17 | 293.183 |
| 253.2185 | 313.1657 | 293.2131 |
| 254.0151 | 313.167 | 293.2145 |
| 254.0168 | 313.1693 | 294.1842 |
| 255.0696 | 314.1622 | 294.2179 |
| 255.0705 | 314.167 | 295.2295 |
| 255.2318 | 314.1685 | 295.2317 |
| 255.2335 | 319.0476 | 296.2325 |
| 256.2359 | 319.0485 | 296.2336 |
| 256.2376 | 319.1216 | 297.1572 |
| 257.2407 | 320.0675 | 306.0789 |
| 258.9827 | 321.0617 | 306.0812 |
| 259.0219 | 321.0642 | 307.0856 |
| 259.0229 | 321.1543 | 307.1953 |
| 259.0237 | 322.0681 | 309.1764 |
| 260.0468 | 323.0298 | 309.209 |
| 260.9803 | 323.0671 | 309.2822 |
| 261.0417 | 323.0681 | 311.2237 |
| 261.0435 | 323.0695 | 311.226 |
| 262.0622 | 323.1342 | 312.1747 |
| 263.0577 | 324.1703 | 312.2282 |
| 263.0595 | 324.1728 | 312.2291 |
| 265.1472 | 325.184 | 320.069 |
| 265.1491 | 326.1871 | 321.0668 |
| 266.151 | 327.1747 | 321.0678 |
| 266.152 | 328.183 | 321.2121 |
| 266.1528 | 332.0305 | 322.0684 |
| 267.0718 | 332.0797 | 322.145 |
| 267.0728 | 333.0266 | 323.0299 |
| 267.2334 | 333.0588 | 323.0323 |
| 269.0856 | 333.0603 | 323.1367 |
| 269.0865 | 333.0615 | 323.1392 |
| 269.0875 | 333.0993 | 325.1864 |
| 269.1215 | 333.1374 | 325.1889 |
| 269.1234 | 335.0794 | 326.1896 |
| 269.2122 | 335.0807 | 326.1922 |
| 269.2492 | 337.0957 | 327.2203 |
| 270.9457 | 337.1831 | 329.235 |
| 271.2283 | 337.1858 | 333.063 |
| 272.9427 | 339.1687 | 337.207 |
| 272.9447 | 339.1971 | 337.2097 |

| | | |
|---|---|---|
| 274.002 | 340.2041 | 339.2009 |
| 274.0029 | 341.0809 | 339.2036 |
| 274.0242 | 341.1106 | 339.2052 |
| 274.0262 | 341.178 | 339.3282 |
| 275.0205 | 341.1931 | 339.3297 |
| 275.0224 | 342.1126 | 341.1107 |
| 275.0575 | 342.1817 | 341.1134 |
| 275.0595 | 346.0567 | 342.1154 |
| 276.0241 | 346.9759 | 342.1165 |
| 276.0261 | 351.0828 | 346.058 |
| 276.0771 | 351.1109 | 346.0608 |
| 277.0325 | 353.0893 | 346.976 |
| 277.0332 | 353.1784 | 346.9769 |
| 277.0726 | 353.2142 | 346.9788 |
| 277.0734 | 353.2171 | 346.9796 |
| 277.2168 | 364.109 | 351.0856 |
| 277.2188 | 365.1396 | 353.2029 |
| 278.0764 | 366.1086 | 366.1104 |
| 278.0777 | 367.1032 | 366.1116 |
| 278.2206 | 367.1053 | 367.1085 |
| 279.0782 | 367.1063 | 367.1097 |
| 279.0794 | 367.2298 | 367.3614 |
| 279.2319 | 368.1093 | 368.1095 |
| 279.2339 | 369.1096 | 368.1107 |
| 280.2367 | 369.1115 | 377.0865 |
| 280.9825 | 369.1176 | 377.0896 |
| 281.2473 | 369.1278 | 378.0909 |
| 281.2485 | 377.0533 | 378.092 |
| 281.2493 | 377.0864 | 378.094 |
| 282.2521 | 379.0686 | 379.0866 |
| 283.1009 | 379.0834 | 380.0914 |
| 283.1019 | 379.1035 | 392.1884 |
| 283.1367 | 379.1055 | 404.1087 |
| 283.1386 | 379.156 | 431.1444 |
| 283.2632 | 380.0862 | 452.2799 |
| 283.2652 | 382.1295 | 473.2862 |
| 284.1412 | 382.1327 | 476.2794 |
| 284.267 | 383.0996 | 476.2821 |
| 284.2683 | 385.0702 | 483.2752 |
| 284.269 | 387.1148 | 483.277 |
| 288.0418 | 391.1405 | 484.2785 |
| 289.0167 | 393.1724 | 487.1818 |
| 289.0356 | 395.2606 | 487.1833 |
| 289.037 | 395.2629 | 507.2742 |

| | | |
|---|---|---|
| 289.0378 | 396.1452 | 528.2482 |
| 289.0726 | 396.2666 | 528.2496 |
| 289.0735 | 397.1499 | 529.2416 |
| 289.0743 | 402.104 | 529.2433 |
| 290.0571 | 404.1029 | 529.2468 |
| 291.0536 | 404.1051 | 554.2637 |
| 291.089 | 405.1078 | |
| 291.0898 | 407.156 | |
| 292.0726 | 407.1581 | |
| 293.0692 | 408.0972 | |
| 293.1219 | 409.1455 | |
| 293.1229 | 410.1612 | |
| 293.1792 | 410.1635 | |
| 293.2131 | 412.1213 | |
| 294.2166 | 413.0762 | |
| 295.1362 | 413.0785 | |
| 295.1384 | 415.073 | |
| 295.2273 | 415.0754 | |
| 295.2296 | 417.103 | |
| 296.2312 | 421.0354 | |
| 296.2324 | 421.2253 | |
| 297.153 | 421.2277 | |
| 298.1569 | 422.0388 | |
| 298.1591 | 423.0319 | |
| 299.1508 | 423.0345 | |
| 299.1531 | 424.0375 | |
| 300.1522 | 424.1781 | |
| 300.1533 | 426.136 | |
| 300.1544 | 426.1371 | |
| 302.0687 | 427.0931 | |
| 302.071 | 427.2871 | |
| 305.0689 | 429.0899 | |
| 306.0758 | 431.1175 | |
| 306.0766 | 433.2352 | |
| 306.088 | 434.2399 | |
| 306.089 | 439.1271 | |
| 307.0824 | 445.1321 | |
| 307.0847 | 445.1334 | |
| 308.0885 | 445.1361 | |
| 309.1182 | 447.2499 | |
| 309.1532 | 447.2529 | |
| 309.2089 | 448.2539 | |
| 309.281 | 448.2556 | |
| 311.1674 | 451.287 | |

| | |
|---|---|
| 312.17 | 461.1514 |
| 312.1724 | 465.3402 |
| 312.1747 | 471.2765 |
| 313.1467 | 473.1626 |
| 313.1671 | 473.2816 |
| 313.1694 | 476.2801 |
| 314.1622 | 479.2064 |
| 314.167 | 479.2079 |
| 314.1685 | 479.3561 |
| 314.1694 | 483.1179 |
| 314.1709 | 483.2724 |
| 315.169 | 484.2783 |
| 315.1714 | 487.1785 |
| 319.0485 | 487.1816 |
| 319.1216 | 494.2813 |
| 320.065 | 495.1392 |
| 320.0674 | 495.1985 |
| 320.0683 | 495.2005 |
| 321.0617 | 495.2759 |
| 321.0631 | 495.2791 |
| 321.0642 | 496.1447 |
| 321.1529 | 503.1968 |
| 321.1541 | 507.2726 |
| 322.0681 | 507.2742 |
| 323.0298 | 508.276 |
| 323.0671 | 511.1954 |
| 323.0679 | 515.269 |
| 323.0695 | 527.2438 |
| 323.133 | 528.1617 |
| 323.1342 | 528.2443 |
| 323.1357 | 528.2481 |
| 323.169 | 529.1581 |
| 324.1378 | 529.2395 |
| 324.1703 | 530.1609 |
| 325.1562 | 533.1731 |
| 325.1578 | 535.2293 |
| 325.184 | 535.231 |
| 326.1593 | 551.2266 |
| 326.1606 | 554.2582 |
| 326.1845 | 555.264 |
| 327.1763 | 563.3364 |
| 327.1838 | 564.3388 |
| 327.219 | 564.3409 |
| 327.2203 | 571.1907 |

| | |
|---|---|
| 328.1769 | 571.1928 |
| 328.1794 | 572.1921 |
| 328.1831 | 573.1873 |
| 329.1429 | 573.1893 |
| 329.1441 | 573.1931 |
| 329.1864 | 574.1957 |
| 330.0474 | 579.3297 |
| 332.0305 | 580.3331 |
| 333.0266 | 580.3353 |
| 333.0292 | 587.2941 |
| 333.0603 | 587.3358 |
| 333.0629 | 603.3474 |
| 333.1006 | 617.292 |
| 333.1366 | 619.3621 |
| 335.0794 | 627.3644 |
| 335.1162 | 631.305 |
| 335.1687 | 631.3088 |
| 337.0941 | 631.3106 |
| 337.0957 | 631.3134 |
| 337.1832 | 632.3121 |
| 337.1858 | 632.3147 |
| 338.1865 | 633.3063 |
| 338.1877 | 637.3715 |
| 338.1892 | 639.2659 |
| 339.165 | 639.2684 |
| 339.1688 | 639.2728 |
| 339.1714 | 640.2696 |
| 339.197 | 641.2722 |
| 339.1997 | 643.3613 |
| 340.2041 | 645.3259 |
| 341.1079 | 645.3282 |
| 341.1106 | 646.3249 |
| 341.1429 | 646.3279 |
| 341.1438 | 646.3319 |
| 341.1915 | 647.285 |
| 341.1931 | 653.2842 |
| 342.1126 | 653.2861 |
| 342.1922 | 654.2831 |
| 342.1993 | 654.2874 |
| 342.2011 | 654.2905 |
| 343.1599 | 655.2874 |
| 344.1628 | 659.335 |
| 346.0571 | 659.3449 |
| 346.058 | 660.3399 |

| | |
|---|---|
| 349.0598 | 660.343 |
| 349.096 | 660.3472 |
| 351.0753 | 661.2994 |
| 351.0828 | 661.3167 |
| 351.1109 | 663.3 |
| 351.112 | 667.2978 |
| 351.201 | 667.3007 |
| 352.0933 | 667.3037 |
| 353.0893 | 668.3048 |
| 353.0922 | 669.3036 |
| 353.1784 | 671.4643 |
| 353.1802 | 672.4702 |
| 353.2142 | 673.3533 |
| 353.2171 | 673.4732 |
| 354.2205 | 673.4784 |
| 355.1589 | 674.3615 |
| 364.109 | 675.3139 |
| 365.0888 | 676.3135 |
| 365.0901 | 676.3162 |
| 365.1395 | 676.321 |
| 365.1407 | 682.3196 |
| 365.1425 | 685.4794 |
| 366.1087 | 685.4853 |
| 366.1107 | 686.4785 |
| 367.1023 | 686.4832 |
| 367.1053 | 686.4863 |
| 367.2299 | 687.4596 |
| 367.2319 | 689.3343 |
| 367.3595 | 695.4628 |
| 368.1093 | 695.4652 |
| 368.2364 | 695.4681 |
| 369.1095 | 696.4656 |
| 369.1115 | 696.4675 |
| 369.1176 | 697.4753 |
| 369.1192 | 697.4809 |
| 369.1267 | 698.4827 |
| 369.1278 | 699.4946 |
| 375.1859 | 709.4851 |
| 376.0579 | 711.4617 |
| 377.0531 | 714.5075 |
| 377.0551 | 716.5955 |
| 377.0864 | 716.5985 |
| 378.0876 | 717.5978 |
| 378.1076 | 717.6007 |

| | |
|---|---|
| 378.1096 | 721.5032 |
| 378.1108 | 722.5074 |
| 379.0686 | 725.3904 |
| 379.0707 | 727.4544 |
| 379.0834 | 728.4591 |
| 379.1035 | 732.513 |
| 379.1054 | 734.513 |
| 379.1067 | 738.5092 |
| 379.1538 | 743.6151 |
| 379.156 | 744.6191 |
| 379.157 | 745.5032 |
| 380.0862 | 746.5002 |
| 380.1084 | 746.5056 |
| 380.1099 | 746.5089 |
| 381.0857 | 747.5085 |
| 381.0867 | 747.5134 |
| 382.1295 | 748.5149 |
| 382.1305 | 760.5441 |
| 382.1327 | 762.5465 |
| 382.2062 | 763.5474 |
| 382.251 | 769.5027 |
| 382.2531 | 775.5374 |
| 383.0611 | 776.5391 |
| 383.0996 | 776.5454 |
| 383.1007 | 787.5471 |
| 383.1028 | 789.4224 |
| 383.1434 | 790.4292 |
| 385.0703 | 792.4614 |
| 387.1136 | 803.4424 |
| 392.102 | 803.4459 |
| 392.1032 | 803.5669 |
| 392.1363 | 804.5716 |
| 392.1385 | 804.578 |
| 393.1036 | 805.5781 |
| 393.1703 | 806.5767 |
| 393.1736 | 822.492 |
| 395.1017 | 822.497 |
| 395.1285 | 831.5967 |
| 395.263 | 832.5992 |
| 396.1452 | 833.5145 |
| 396.1485 | 834.514 |
| 396.2666 | 834.5208 |
| 397.1488 | 834.5243 |
| 397.1499 | 835.5302 |

| | |
|---|---|
| 398.1054 | 848.5276 |
| 404.1052 | 849.5278 |
| 405.1078 | 850.5259 |
| 407.156 | 851.5278 |
| 408.0978 | |
| 409.1455 | |
| 409.1474 | |
| 409.2788 | |
| 409.2801 | |
| 410.161 | |
| 410.1635 | |
| 411.1638 | |
| 411.1658 | |
| 412.1213 | |
| 413.075 | |
| 413.0761 | |
| 413.0786 | |
| 415.0729 | |
| 415.0754 | |
| 417.1112 | |
| 421.0354 | |
| 421.0369 | |
| 421.2253 | |
| 422.0388 | |
| 422.04 | |
| 423.0345 | |
| 423.161 | |
| 423.2393 | |
| 423.253 | |
| 423.2542 | |
| 424.0375 | |
| 424.1779 | |
| 424.1795 | |
| 426.1371 | |
| 427.0907 | |
| 427.093 | |
| 427.2871 | |
| 428.9469 | |
| 429.0913 | |
| 431.1233 | |
| 431.1252 | |
| 431.1271 | |
| 433.2352 | |
| 433.2364 | |

433.239
434.2399
437.0951
437.0967
437.0991
439.0822
439.0839
439.0851
439.1272
439.1296
444.9193
445.1401
445.1421
445.1441
447.2528
448.254
448.2565
450.2907
450.2921
451.2874
451.2898
451.3267
452.2921
461.2662
461.269
463.2475
465.3059
465.3084
465.3402
465.3432
467.1706
467.2311
471.2766
471.2795
473.2832
473.2859
476.2776
476.2803
479.2034
479.2062
479.2079
479.2195
479.2213
479.3561

| |
|---|
| 479.358 |
| 480.2102 |
| 482.12 |
| 483.1179 |
| 483.2724 |
| 483.2737 |
| 483.2769 |
| 484.1188 |
| 484.1218 |
| 484.275 |
| 484.2783 |
| 487.1784 |
| 487.1816 |
| 494.2791 |
| 494.2826 |
| 495.1392 |
| 495.2005 |
| 495.2035 |
| 495.2759 |
| 496.1429 |
| 496.2027 |
| 496.2061 |
| 496.2818 |
| 501.1955 |
| 507.1682 |
| 507.2726 |
| 507.2758 |
| 508.276 |
| 511.194 |
| 511.1955 |
| 515.2657 |
| 515.269 |
| 519.2344 |
| 519.236 |
| 519.2395 |
| 521.1835 |
| 528.2463 |
| 528.248 |
| 529.1476 |
| 529.1566 |
| 529.1581 |
| 529.1617 |
| 529.2414 |
| 529.2443 |

529.2831
530.1645
535.2293
535.2329
535.2346
545.1214
551.2266
555.1559
555.2679
559.1389
563.3364
564.3351
564.3388
564.3408
571.1907
572.1931
573.1893
573.1931
574.1962
577.3543
579.3297
580.3331
580.3351
587.0288
587.301
587.321
587.3331
587.3356
588.3391
589.3538
603.3471
603.351
604.335
606.3758
606.3801
619.36
619.3623
622.3711
627.3655
627.3692
631.3066
631.3085
631.3134
632.3121

632.3146
634.3743
637.3687
637.3708
637.3756
638.3748
638.3778
639.2659
639.27
639.2727
640.2696
641.2757
643.3617
643.3649
645.324
645.3276
645.333
646.328
646.33
647.285
647.2891
647.306
650.3675
653.2788
653.2816
653.286
653.3648
653.3698
654.2874
655.2837
655.2908
656.2894
659.337
659.3424
660.343
660.345
660.3473
661.2993
661.3022
662.3045
667.3007
667.3042
668.3048
671.3303

671.4643
672.4647
672.4703
672.4723
673.3588
673.471
673.4733
673.4784
673.481
674.3584
674.3615
675.3139
675.3185
676.321
677.4057
681.3144
681.3194
682.3164
682.3196
685.4794
686.4808
686.4844
686.4862
688.468
689.3297
689.3342
695.4628
695.4677
696.4656
696.4735
697.4731
697.4778
697.481
698.4828
698.4879
700.5009
703.455
704.4593
709.4851
710.4847
710.4885
711.4617
712.4287
712.4694

713.4793
714.5076
714.5128
715.5873
716.5956
717.5977
717.6024
718.6021
721.4973
721.5032
721.5056
722.5015
722.5074
722.51
723.5054
723.509
725.388
725.3904
726.4447
727.4508
727.4544
727.4592
728.4591
729.4693
738.5031
738.5058
739.5086
741.3835
741.3876
742.5028
743.4871
743.6153
743.6206
744.4888
744.6191
745.5008
745.5062
746.5028
746.5089
747.5133
747.5153
748.511
748.5137
748.5197

749.5281
760.5443
760.5507
761.4985
761.5206
761.5247
762.5014
762.5465
763.5474
763.5523
767.5342
769.5028
770.5034
775.5374
776.5409
777.4904
777.4961
778.493
789.4225
789.426
790.4292
791.533
793.5184
803.4403
803.567
804.4447
804.4504
804.5716
804.5747
817.4548
817.4606
818.4615
818.4655
820.4927
831.5928
832.5991
833.5146
834.5243
835.5256
848.5276
849.5278
850.5189
851.5279

**Supplemental Table 10**

| Shared m/z | unique_to_B73 | unique_to_OG |
|---|---|---|
| 71.013935 | 89.0249397 | 73.029838 |
| 78.959158 | 96.9701443 | 96.9626762 |
| 85.03005 | 98.025334 | 102.949208 |
| 86.033229 | 100.933958 | 103.920645 |
| 90.934202 | 102.952995 | 112.937879 |
| 96.960502 | 107.044554 | 117.056099 |
| 99.00936 | 112.986057 | 127.051514 |
| 101.024813 | 113.926837 | 131.023111 |
| 104.035734 | 114.019994 | 131.034812 |
| 106.041668 | 126.881369 | 136.862659 |
| 110.985538 | 128.878412 | 139.003961 |
| 111.009358 | 129.055555 | 146.924568 |
| 113.024774 | 131.036684 | 146.966449 |
| 113.035962 | 132.041523 | 148.948909 |
| 113.989414 | 132.043892 | 150.020165 |
| 115.004165 | 133.033449 | 153.044395 |
| 116.007421 | 133.050854 | 167.033833 |
| 117.020129 | 137.892435 | 169.835927 |
| 118.023171 | 144.99212 | 178.052131 |
| 118.051186 | 145.941902 | 181.039114 |
| 119.035334 | 147.050293 | 184.985462 |
| 120.013518 | 147.97018 | 185.008129 |
| 122.025405 | 148.919113 | 186.114373 |
| 128.03566 | 159.031249 | 192.013907 |
| 128.959662 | 163.04166 | 192.030956 |
| 129.019977 | 167.838403 | 194.029263 |
| 129.976237 | 168.044435 | 197.065501 |
| 130.023318 | 170.835054 | 197.808711 |
| 130.983448 | 170.903397 | 202.887778 |
| 131.024877 | 173.046367 | 203.092698 |
| 131.046364 | 175.013767 | 203.104454 |
| 132.030277 | 175.024985 | 204.975909 |
| 132.86786 | 175.048518 | 205.840394 |
| 133.01463 | 177.093958 | 206.164145 |
| 134.017602 | 180.973482 | 207.837345 |
| 134.025113 | 187.941821 | 211.002234 |
| 134.864747 | 189.08915 | 215.86928 |
| 135.018376 | 191.058542 | 220.147666 |
| 135.028679 | 192.060227 | 221.843257 |
| 136.909688 | 193.028949 | 224.091508 |

| | | |
|---|---|---|
| 138.05617 | 193.816281 | 229.012612 |
| 143.034954 | 196.054988 | 229.024425 |
| 145.014504 | 199.034465 | 237.062289 |
| 145.049728 | 202.117312 | 240.052015 |
| 145.051302 | 203.021236 | 242.081435 |
| 145.062065 | 204.979575 | 244.155767 |
| 146.046043 | 207.013519 | 250.147196 |
| 146.054536 | 207.015453 | 255.151152 |
| 146.065439 | 209.030407 | 257.114027 |
| 146.923527 | 209.067025 | 259.023842 |
| 146.93891 | 216.936967 | 259.130302 |
| 147.029607 | 217.866347 | 260.026309 |
| 147.030776 | 222.159005 | 262.032468 |
| 147.067038 | 223.045615 | 262.034772 |
| 148.940256 | 223.095679 | 264.991715 |
| 148.951487 | 225.898165 | 265.120837 |
| 149.012861 | 226.997043 | 265.148611 |
| 149.025405 | 228.051309 | 266.979674 |
| 150.016041 | 229.023424 | 266.986487 |
| 153.868583 | 229.161628 | 267.106546 |
| 154.0631 | 231.099268 | 267.144574 |
| 154.064585 | 233.155879 | 268.939477 |
| 155.86564 | 236.024405 | 271.915748 |
| 156.068064 | 239.128029 | 272.087721 |
| 157.051425 | 250.070059 | 272.968801 |
| 158.980341 | 255.823881 | 273.031137 |
| 159.067314 | 257.154824 | 275.01765 |
| 159.981046 | 257.820875 | 275.020126 |
| 160.843784 | 262.905752 | 275.022786 |
| 160.977865 | 262.908187 | 276.02702 |
| 161.046576 | 263.165317 | 278.146664 |
| 162.840997 | 265.092803 | 278.151389 |
| 163.999289 | 269.054229 | 279.13385 |
| 164.036942 | 269.130816 | 279.164234 |
| 164.073659 | 274.955662 | 285.109284 |
| 164.838017 | 277.220725 | 291.116596 |
| 164.928423 | 277.910023 | 291.133978 |
| 165.019607 | 278.027459 | 291.160895 |
| 165.040298 | 279.037557 | 292.071883 |
| 166.040648 | 279.107555 | 292.073678 |
| 166.052188 | 280.032626 | 293.142264 |
| 168.067689 | 280.034892 | 293.149202 |

| | | |
|---|---|---|
| 168.838154 | 280.042382 | 294.069299 |
| 170.872134 | 282.116311 | 296.882828 |
| 171.007616 | 289.030096 | 298.114751 |
| 171.067306 | 289.937987 | 298.154832 |
| 172.09866 | 289.942802 | 301.145115 |
| 172.869212 | 291.199897 | 302.100101 |
| 173.010295 | 291.201628 | 302.101564 |
| 173.083478 | 293.125989 | 302.135933 |
| 174.013566 | 293.175941 | 303.106707 |
| 175.062326 | 293.177619 | 303.118882 |
| 176.936967 | 293.180739 | 303.120806 |
| 177.041668 | 293.210695 | 303.216725 |
| 179.023626 | 293.212296 | 304.12341 |
| 179.057344 | 294.184113 | 304.993484 |
| 180.026771 | 295.103411 | 305.895375 |
| 180.060571 | 296.231625 | 307.919803 |
| 180.068304 | 297.874478 | 308.861642 |
| 180.901332 | 299.870833 | 308.916627 |
| 180.932025 | 305.019037 | 309.036657 |
| 182.898121 | 305.022672 | 309.916942 |
| 182.994158 | 305.979314 | 310.172652 |
| 183.067674 | 306.05675 | 311.079281 |
| 184.986614 | 306.080889 | 311.109594 |
| 185.022307 | 307.144469 | 311.115034 |
| 185.156199 | 308.075038 | 313.217837 |
| 187.098518 | 309.121081 | 313.236691 |
| 187.135333 | 309.168218 | 315.100789 |
| 187.862429 | 309.206638 | 315.108635 |
| 187.929909 | 309.999891 | 315.253529 |
| 188.057328 | 311.213902 | 317.106083 |
| 188.934402 | 311.214931 | 319.064896 |
| 189.003638 | 311.221389 | 320.069792 |
| 189.004733 | 311.22359 | 320.189346 |
| 189.859082 | 312.126295 | 320.8842 |
| 189.932563 | 312.12768 | 322.143511 |
| 190.929449 | 312.173685 | 323.016926 |
| 191.019864 | 312.22564 | 323.045151 |
| 191.056453 | 312.229141 | 323.138634 |
| 191.85632 | 313.04585 | 325.049773 |
| 191.932754 | 313.047243 | 325.150921 |
| 192.023545 | 316.946837 | 326.127951 |
| 192.929501 | 317.142414 | 326.129885 |

| | | |
|---|---|---|
| 193.023661 | 320.886375 | 326.146862 |
| 193.036215 | 321.04765 | 326.151237 |
| 193.051763 | 322.042203 | 326.82811 |
| 193.088092 | 322.043791 | 327.090862 |
| 194.046647 | 322.199446 | 327.127761 |
| 194.083088 | 323.035485 | 328.057853 |
| 194.924958 | 325.13051 | 328.062969 |
| 194.947163 | 325.172615 | 330.092692 |
| 194.991364 | 327.10819 | 331.250573 |
| 195.050011 | 327.215255 | 333.939048 |
| 195.051704 | 327.262455 | 334.059685 |
| 195.086273 | 328.042995 | 336.865834 |
| 195.139934 | 328.05579 | 337.117452 |
| 196.050433 | 328.219957 | 338.987629 |
| 196.06225 | 329.243153 | 338.990618 |
| 197.890661 | 330.064163 | 339.153091 |
| 199.805598 | 330.233569 | 339.163281 |
| 201.114023 | 330.23577 | 339.236707 |
| 203.020016 | 332.070683 | 340.092281 |
| 203.082657 | 335.08734 | 341.075537 |
| 205.078028 | 335.847736 | 341.1074 |
| 205.161241 | 336.859011 | 341.121661 |
| 206.082674 | 339.996818 | 341.123573 |
| 206.973512 | 339.999161 | 341.270738 |
| 207.103479 | 341.118774 | 342.114067 |
| 207.919038 | 341.198051 | 343.083492 |
| 207.931995 | 341.240493 | 343.10531 |
| 208.098794 | 343.266694 | 345.079804 |
| 208.106789 | 343.268278 | 346.044055 |
| 209.155549 | 344.047687 | 347.03726 |
| 210.02365 | 353.150112 | 347.061807 |
| 210.041207 | 354.056948 | 349.112472 |
| 210.158825 | 356.100041 | 351.154136 |
| 211.087906 | 357.091032 | 351.163253 |
| 214.880552 | 357.093136 | 352.13379 |
| 215.033385 | 359.996952 | 353.053014 |
| 215.129651 | 361.086632 | 355.320145 |
| 216.036087 | 362.047724 | 356.150716 |
| 216.037344 | 364.068166 | 356.187329 |
| 217.029708 | 367.102653 | 356.242617 |
| 217.031105 | 368.993824 | 356.999119 |
| 218.033668 | 371.099928 | 357.096275 |

| | | |
|---|---|---|
| 218.104144 | 371.101211 | 357.112621 |
| 220.062312 | 372.138404 | 357.190685 |
| 221.155427 | 373.181328 | 358.259192 |
| 223.08863 | 373.191075 | 358.262636 |
| 224.056771 | 374.264331 | 359.102438 |
| 224.058051 | 375.033452 | 361.079781 |
| 224.093434 | 375.087299 | 362.99859 |
| 225.060756 | 375.267077 | 363.057308 |
| 225.062159 | 376.122786 | 365.059803 |
| 225.088725 | 377.039858 | 365.06221 |
| 225.150152 | 379.02474 | 365.065985 |
| 226.966891 | 380.085175 | 365.07131 |
| 227.128935 | 380.104247 | 367.063418 |
| 228.065165 | 380.156813 | 368.236524 |
| 228.161048 | 380.15965 | 370.010244 |
| 230.968582 | 383.136407 | 370.203166 |
| 231.932023 | 383.358392 | 372.059694 |
| 232.023318 | 384.160579 | 372.083638 |
| 232.059787 | 384.169451 | 372.096434 |
| 234.159386 | 384.903307 | 374.224874 |
| 237.036058 | 384.98175 | 375.133192 |
| 238.933067 | 385.117513 | 376.245119 |
| 239.129467 | 386.900192 | 377.164478 |
| 240.050589 | 387.15844 | 378.047919 |
| 241.145296 | 387.172052 | 378.092615 |
| 242.176682 | 388.161828 | 379.032251 |
| 242.941064 | 389.997783 | 379.15657 |
| 243.039145 | 390.991444 | 380.052149 |
| 243.062616 | 391.272111 | 381.020622 |
| 243.160915 | 391.290388 | 383.065846 |
| 243.179592 | 392.290129 | 383.107091 |
| 245.043513 | 392.294032 | 383.126647 |
| 246.01812 | 393.292979 | 383.193769 |
| 248.032943 | 394.240609 | 384.193219 |
| 248.054466 | 396.08083 | 384.939907 |
| 248.962276 | 397.083191 | 385.067955 |
| 249.06754 | 397.255484 | 386.075267 |
| 249.150443 | 397.942426 | 386.103009 |
| 250.071277 | 398.25875 | 386.932956 |
| 250.145902 | 399.214504 | 387.082486 |
| 251.050008 | 399.219161 | 387.083545 |
| 251.051238 | 399.252176 | 387.099833 |

| | | |
|---|---|---|
| 251.093248 | 400.948148 | 389.098238 |
| 251.148914 | 402.100981 | 389.272235 |
| 253.145129 | 402.167523 | 389.273854 |
| 254.148356 | 403.166966 | 390.117764 |
| 258.91621 | 404.153011 | 391.033608 |
| 259.022733 | 404.161175 | 391.262731 |
| 259.12881 | 405.046199 | 391.291772 |
| 260.031951 | 406.167794 | 392.086084 |
| 260.915173 | 406.956263 | 392.167824 |
| 262.92982 | 408.161955 | 393.018156 |
| 264.157622 | 409.238082 | 393.045638 |
| 264.929314 | 410.31687 | 393.133271 |
| 265.066548 | 411.321074 | 393.168183 |
| 265.118967 | 412.202556 | 394.890984 |
| 266.15192 | 412.972636 | 394.895778 |
| 267.072984 | 412.973829 | 396.218866 |
| 269.138963 | 413.128457 | 397.078504 |
| 269.211932 | 414.132063 | 400.04778 |
| 269.918359 | 415.162416 | 401.926235 |
| 270.142298 | 415.188296 | 402.01287 |
| 270.206614 | 415.272242 | 402.172298 |
| 270.979191 | 417.23653 | 402.996786 |
| 271.070984 | 418.237929 | 403.109491 |
| 271.228678 | 419.07532 | 403.111708 |
| 272.231541 | 419.077681 | 404.085828 |
| 272.933989 | 421.043451 | 404.111011 |
| 272.936392 | 421.166012 | 405.143097 |
| 272.957993 | 421.262822 | 408.056191 |
| 273.17049 | 422.247951 | 409.257937 |
| 273.95915 | 422.929729 | 410.063379 |
| 275.12462 | 425.258691 | 410.214411 |
| 276.161299 | 426.02332 | 410.864484 |
| 277.032981 | 427.220283 | 411.080241 |
| 277.217351 | 427.263331 | 411.084104 |
| 277.907746 | 428.204297 | 411.097565 |
| 278.036648 | 428.206809 | 411.348612 |
| 279.038853 | 429.080223 | 412.211415 |
| 279.234676 | 429.082659 | 418.084923 |
| 279.903553 | 430.055199 | 418.099343 |
| 280.908602 | 430.103552 | 418.102981 |
| 280.994929 | 431.12599 | 420.106685 |
| 281.029668 | 431.190999 | 420.995677 |

| | | |
|---|---|---|
| 281.035869 | 431.19412 | 421.072244 |
| 282.083356 | 431.246574 | 423.093713 |
| 284.098223 | 432.050946 | 423.25265 |
| 285.205679 | 433.167945 | 424.084116 |
| 286.128634 | 435.916404 | 425.08774 |
| 287.221723 | 437.24623 | 425.237537 |
| 288.084929 | 437.292428 | 426.041171 |
| 289.017343 | 437.296542 | 426.09933 |
| 289.019177 | 438.295711 | 426.310459 |
| 289.067766 | 439.174641 | 426.978269 |
| 289.119593 | 440.068349 | 427.027737 |
| 291.049532 | 441.022793 | 429.155751 |
| 292.960733 | 441.078606 | 429.163411 |
| 293.121924 | 441.094317 | 429.196102 |
| 294.029649 | 441.099458 | 430.053721 |
| 294.178798 | 441.251189 | 432.121481 |
| 295.038872 | 442.090142 | 433.117572 |
| 295.228348 | 443.139978 | 433.119022 |
| 296.043343 | 443.193825 | 433.153348 |
| 297.153245 | 443.256338 | 434.111242 |
| 297.244325 | 443.272413 | 434.119463 |
| 298.113567 | 444.068969 | 434.121498 |
| 298.15676 | 445.108296 | 434.136289 |
| 299.918315 | 447.225538 | 435.092042 |
| 301.046155 | 448.188604 | 436.104211 |
| 301.104629 | 449.121419 | 438.086087 |
| 302.065767 | 449.190456 | 439.074852 |
| 304.991095 | 449.245589 | 439.086177 |
| 305.216554 | 449.256979 | 439.254715 |
| 306.075977 | 450.10576 | 440.066762 |
| 306.078006 | 451.241887 | 440.079729 |
| 306.079308 | 452.299145 | 441.006534 |
| 306.919461 | 454.136134 | 441.211041 |
| 307.081474 | 455.148508 | 442.018773 |
| 307.141092 | 457.296672 | 444.098764 |
| 307.854018 | 459.137614 | 445.102341 |
| 311.100903 | 459.274333 | 447.242016 |
| 311.134018 | 459.278611 | 448.11468 |
| 311.168769 | 460.277416 | 448.143625 |
| 311.201837 | 461.106502 | 449.100424 |
| 311.226194 | 461.122001 | 450.117419 |
| 312.094297 | 462.073758 | 450.184885 |

| | | |
|---|---|---|
| 312.172486 | 462.881675 | 451.149945 |
| 312.838294 | 463.220441 | 453.106027 |
| 313.165281 | 463.225211 | 453.271566 |
| 313.23907 | 469.370167 | 454.29054 |
| 315.061218 | 473.163444 | 455.189971 |
| 315.109744 | 473.268824 | 455.191415 |
| 315.254933 | 473.271243 | 459.071232 |
| 316.948093 | 474.171027 | 460.065092 |
| 317.021072 | 475.215456 | 460.066889 |
| 317.137782 | 476.296273 | 460.11245 |
| 321.063564 | 478.135172 | 461.067662 |
| 321.129309 | 479.215662 | 462.060796 |
| 323.026858 | 480.131865 | 462.062309 |
| 323.028792 | 480.30808 | 462.066508 |
| 323.135517 | 484.279959 | 462.127177 |
| 323.190667 | 487.170283 | 462.135475 |
| 324.029851 | 488.167793 | 465.101237 |
| 324.033057 | 489.277968 | 468.032607 |
| 325.018482 | 492.280326 | 468.898442 |
| 325.12366 | 493.271361 | 469.15939 |
| 325.137914 | 495.251153 | 470.089596 |
| 325.140074 | 496.119877 | 471.109055 |
| 325.14875 | 501.133154 | 471.121042 |
| 325.184989 | 505.308733 | 472.111709 |
| 326.141242 | 507.108171 | 472.126812 |
| 326.18828 | 507.273443 | 473.169134 |
| 326.905407 | 507.28276 | 473.269918 |
| 327.182179 | 508.286015 | 474.067345 |
| 327.254145 | 511.191788 | 474.149079 |
| 327.291113 | 512.047355 | 474.169455 |
| 328.059496 | 515.124134 | 474.172535 |
| 328.061247 | 521.389719 | 475.118554 |
| 328.294747 | 522.120783 | 475.123734 |
| 329.076198 | 524.117547 | 476.124865 |
| 329.085958 | 529.285219 | 476.139108 |
| 329.08855 | 530.305375 | 478.123883 |
| 329.231056 | 533.137559 | 479.033505 |
| 331.015687 | 535.042267 | 480.301065 |
| 331.243903 | 535.153013 | 481.125849 |
| 331.246073 | 535.15433 | 483.1045 |
| 331.247232 | 535.523141 | 483.27566 |
| 331.259307 | 537.272629 | 485.271794 |

| | | |
|---|---|---|
| 332.251007 | 540.326281 | 485.285156 |
| 333.057446 | 540.328699 | 486.057795 |
| 333.058757 | 541.336608 | 486.292033 |
| 333.189687 | 549.357668 | 487.117117 |
| 333.850108 | 556.308477 | 487.119217 |
| 334.074051 | 557.310627 | 487.180487 |
| 336.855382 | 559.310921 | 487.214607 |
| 338.858545 | 570.237159 | 491.082137 |
| 339.200581 | 571.295834 | 491.254921 |
| 339.233697 | 571.499021 | 491.272468 |
| 339.290976 | 571.501455 | 492.275779 |
| 340.203861 | 572.502412 | 493.105121 |
| 340.237108 | 573.137045 | 493.129281 |
| 340.862383 | 573.179941 | 493.232606 |
| 341.108967 | 573.495576 | 496.102865 |
| 341.110355 | 582.531874 | 496.12118 |
| 342.082521 | 585.33762 | 497.108605 |
| 342.112129 | 598.250856 | 498.041355 |
| 342.170611 | 598.281925 | 500.925992 |
| 342.85521 | 598.51837 | 501.069013 |
| 344.031266 | 599.253004 | 501.103302 |
| 344.033692 | 600.24762 | 502.060368 |
| 344.038023 | 600.301331 | 504.311747 |
| 346.052402 | 601.250001 | 505.312989 |
| 346.056274 | 601.299607 | 508.276335 |
| 347.025613 | 601.300731 | 510.882488 |
| 347.038656 | 602.244371 | 511.097745 |
| 347.041298 | 606.079996 | 513.087907 |
| 347.054292 | 616.290098 | 513.201572 |
| 347.060241 | 617.460967 | 520.269369 |
| 349.074424 | 622.279644 | 520.271896 |
| 351.130491 | 634.226132 | 527.14052 |
| 353.039823 | 636.223078 | 527.143011 |
| 353.050846 | 637.150638 | 532.480155 |
| 353.164651 | 678.547783 | 540.055276 |
| 354.053496 | 678.549404 | 541.058753 |
| 355.156592 | 685.506538 | 541.322724 |
| 355.159075 | 686.484846 | 544.266842 |
| 355.228527 | 687.487472 | 548.143185 |
| 356.096517 | 710.484556 | 548.236969 |
| 356.153876 | 734.469594 | 551.131049 |
| 356.15932 | 754.470198 | 555.119665 |

| | | |
|---|---|---|
| 356.161554 | 766.434297 | 556.123305 |
| 357.08592 | 767.531097 | 559.283683 |
| 357.122866 | 768.432271 | 561.215076 |
| 357.164466 | 768.59229 | 564.159392 |
| 362.051637 | 784.446001 | 564.160526 |
| 363.005002 | 786.44229 | 565.048982 |
| 363.16283 | 806.186351 | 565.082964 |
| 364.059883 | | 565.336261 |
| 367.908757 | | 566.338042 |
| 368.10191 | | 571.16498 |
| 368.237765 | | 571.23203 |
| 369.008203 | | 573.123447 |
| 369.301372 | | 573.141853 |
| 370.01198 | | 575.109878 |
| 370.984676 | | 575.112013 |
| 372.095016 | | 575.113819 |
| 373.098268 | | 588.898649 |
| 374.133329 | | 599.532258 |
| 374.240277 | | 601.143001 |
| 374.24727 | | 603.004313 |
| 374.250859 | | 606.074662 |
| 374.907791 | | 612.147537 |
| 375.135718 | | 617.485796 |
| 375.243607 | | 629.095561 |
| 375.254058 | | 637.138227 |
| 376.130141 | | 641.172935 |
| 376.256548 | | 679.548074 |
| 376.258102 | | 687.556457 |
| 377.083299 | | 703.134613 |
| 377.086525 | | 712.514347 |
| 377.132755 | | 712.540566 |
| 378.0639 | | 714.549108 |
| 378.090814 | | 728.151851 |
| 379.013788 | | 745.194424 |
| 379.071623 | | 759.547801 |
| 379.078374 | | 759.550356 |
| 379.081797 | | 768.591288 |
| 379.158644 | | 799.557331 |
| 379.192302 | | 807.201494 |
| 379.203113 | | |
| 380.161474 | | |
| 380.19055 | | |

381.174863
383.064093
383.191574
384.067431
384.194319
384.936848
384.982761
385.108055
385.116399
385.119794
386.114518
386.946324
387.117042
387.11836
388.092499
388.121088
389.122254
389.246882
390.24237
390.99029
391.162653
391.164056
391.287334
392.166065
392.188667
393.172544
394.894353
396.091192
397.093757
398.132993
398.234563
399.101782
399.135499
399.238124
399.938296
399.94137
400.105021
400.129797
400.148767
400.94627
401.172548
402.10623

402.108711
402.164481
402.175839
404.044685
404.108024
404.155719
405.030669
405.965892
406.954698
407.18867
408.071799
408.073127
409.076155
409.312411
410.069273
410.075528
410.226627
410.228358
410.315495
411.221282
411.222287
411.318772
412.22496
415.217576
416.0753
417.07851
417.212083
417.234514
418.00695
418.07222
418.073435
418.10184
418.214823
419.105357
420.06906
420.22819
420.242851
420.25323
420.255761
420.256773
420.268692
421.074492

421.078127
421.24577
421.259953
422.090929
422.263772
422.92829
423.096085
423.097876
424.098755
425.082637
425.1102
425.30721
426.024661
426.027316
426.97524
427.225374
427.241068
430.000711
431.213515
432.11953
433.116271
433.237682
434.241103
435.211577
435.895508
435.914931
436.25444
437.239616
437.247258
438.083326
439.08976
439.091568
440.075395
440.081114
440.082828
440.093614
441.084873
441.110401
441.114581
441.252927
441.338218
442.072086

442.095455
442.118326
442.895606
443.094392
443.248464
444.109674
444.112224
445.113323
445.115536
445.369789
447.089515
447.124069
447.139969
447.257163
448.111689
448.139016
448.260554
449.11289
449.115147
451.051055
451.973004
452.061524
452.268469
452.277281
452.283026
452.924506
452.925895
453.104865
453.194649
453.286967
454.153854
455.10218
455.10512
455.35787
456.150555
457.052969
457.141736
459.106067
461.132274
463.096755
463.217594
466.091706

466.094305
467.097465
469.144711
469.374316
470.094325
471.079642
471.122464
473.166337
473.246459
474.068874
474.120406
474.123056
475.121908
475.13363
475.13538
476.13693
476.280157
477.28384
478.284864
479.12204
479.124703
480.125322
480.306989
480.312029
480.328336
481.314273
481.981143
482.105026
483.276909
488.134976
490.130574
491.133268
491.268297
491.269633
492.127373
492.271844
492.272966
493.130478
493.274639
493.28239
494.124081
494.288189

495.12713
496.105218
499.116662
499.120553
499.250957
500.118882
501.073891
501.155192
503.16393
503.169575
504.172965
504.308723
504.312785
505.125084
505.316396
507.269501
507.283777
507.303102
507.369848
508.273286
508.306372
509.11563
509.120848
509.226209
511.102426
515.094631
517.128958
518.132124
520.911988
525.151324
527.244582
528.24826
529.111485
529.241109
531.142545
532.145876
533.139704
533.256689
534.150236
536.117894
537.273838
538.27702

540.334932
545.220662
547.139283
553.124343
554.305101
555.221154
561.153293
561.154768
564.329576
564.333921
565.050795
565.080056
565.337348
566.053297
566.34012
569.233476
571.169778
571.49761
572.173064
573.1293
573.130549
576.187122
579.26296
595.291664
596.29574
600.108751
600.299315
601.111878
611.148512
620.134815
627.272922
629.093653
637.141222
639.121802
677.546421
677.548454
678.5513
679.547172
701.151686
712.53878
713.54247
714.54496

740.564499
740.569498
741.573279
749.173286
753.468865
758.543592
767.532765
768.536248
786.574852
791.533198
792.536814
806.197815
819.525463
819.526908
820.530166

**Supplemental Table 11**

| Cluster | m/z value |
|---|---|
| 4 | 110.9759 |
| 4 | 111.0089 |
| 4 | 115.0037 |
| 4 | 115.0042 |
| 4 | 115.0216 |
| 4 | 115.077 |
| 4 | 116.0073 |
| 4 | 116.0079 |
| 4 | 118.9928 |
| 4 | 119.0356 |
| 4 | 121.0298 |
| 4 | 129.0202 |
| 4 | 130.0235 |
| 4 | 131.0352 |
| 4 | 133.0151 |
| 4 | 134.0175 |
| 4 | 134.0182 |
| 4 | 135.0187 |
| 4 | 135.0193 |
| 4 | 135.0301 |
| 4 | 138.0201 |
| 4 | 142.0518 |
| 4 | 145.0513 |
| 4 | 145.0617 |
| 4 | 145.0625 |
| 4 | 146.0466 |
| 4 | 147.0308 |
| 4 | 150.0155 |
| 4 | 153.0225 |
| 4 | 153.0233 |
| 4 | 153.0241 |
| 4 | 155.0275 |
| 4 | 159.0116 |
| 4 | 161.0458 |
| 4 | 162.983 |
| 4 | 165.0406 |
| 4 | 167.9976 |
| 4 | 171.0663 |
| 4 | 171.0673 |
| 4 | 171.1027 |
| 4 | 171.1036 |
| 4 | 173.0104 |

| | |
|---|---|
| 4 | 174.0125 |
| 4 | 175.0145 |
| 4 | 177.0929 |
| 4 | 179.0558 |
| 4 | 179.0569 |
| 4 | 187.0974 |
| 4 | 192.0238 |
| 4 | 192.0602 |
| 4 | 194.9911 |
| 4 | 195.0505 |
| 4 | 197.0302 |
| 4 | 197.119 |
| 4 | 199.0978 |
| 4 | 200.0567 |
| 4 | 201.114 |
| 4 | 211.134 |
| 4 | 211.1353 |
| 4 | 215.0324 |
| 4 | 215.0338 |
| 4 | 217.0168 |
| 4 | 217.0305 |
| 4 | 218.0337 |
| 4 | 224.0572 |
| 4 | 225.0605 |
| 4 | 225.0611 |
| 4 | 227.0392 |
| 4 | 227.1293 |
| 4 | 227.13 |
| 4 | 227.2024 |
| 4 | 230.0364 |
| 4 | 239.0596 |
| 4 | 242.0514 |
| 4 | 242.053 |
| 4 | 244.0468 |
| 4 | 247.0639 |
| 4 | 248.0587 |
| 4 | 253.092 |
| 4 | 253.2168 |
| 4 | 254.0168 |
| 4 | 255.2318 |
| 4 | 256.2376 |
| 4 | 259.0229 |
| 4 | 263.0577 |
| 4 | 266.1528 |

| | |
|---|---|
| 4 | 267.0718 |
| 4 | 267.0728 |
| 4 | 269.1215 |
| 4 | 269.2122 |
| 4 | 269.2492 |
| 4 | 274.0029 |
| 4 | 274.0262 |
| 4 | 275.0205 |
| 4 | 275.0224 |
| 4 | 276.0771 |
| 4 | 277.0332 |
| 4 | 277.0734 |
| 4 | 278.0764 |
| 4 | 279.0794 |
| 4 | 281.2485 |
| 4 | 282.2521 |
| 4 | 283.1019 |
| 4 | 283.2632 |
| 4 | 284.2683 |
| 4 | 293.0692 |
| 4 | 295.2273 |
| 4 | 302.0687 |
| 4 | 302.071 |
| 4 | 306.088 |
| 4 | 307.0847 |
| 4 | 309.281 |
| 4 | 319.0485 |
| 4 | 320.0683 |
| 4 | 321.0642 |
| 4 | 323.0298 |
| 4 | 323.0671 |
| 4 | 327.1763 |
| 4 | 328.1769 |
| 4 | 329.1429 |
| 4 | 333.0603 |
| 4 | 333.0629 |
| 4 | 333.1366 |
| 4 | 335.1162 |
| 4 | 344.1628 |
| 4 | 349.0598 |
| 4 | 351.112 |
| 4 | 365.0888 |
| 4 | 365.1395 |
| 4 | 365.1425 |

| | |
|---|---|
| 4 | 367.1023 |
| 4 | 367.3595 |
| 4 | 369.1095 |
| 4 | 369.1176 |
| 4 | 375.1859 |
| 4 | 381.0857 |
| 4 | 381.0867 |
| 4 | 382.1327 |
| 4 | 382.2062 |
| 4 | 382.2531 |
| 4 | 383.0996 |
| 4 | 392.102 |
| 4 | 392.1032 |
| 4 | 392.1363 |
| 4 | 392.1385 |
| 4 | 393.1036 |
| 4 | 395.1017 |
| 4 | 397.1488 |
| 4 | 410.161 |
| 4 | 410.1635 |
| 4 | 411.1658 |
| 4 | 445.1401 |
| 4 | 473.2832 |
| 4 | 479.2079 |
| 4 | 501.1955 |
| 4 | 507.2758 |
| 4 | 519.236 |
| 4 | 521.1835 |
| 4 | 528.2463 |
| 4 | 528.248 |
| 4 | 529.2443 |
| 4 | 545.1214 |
| 4 | 555.2679 |
| 4 | 563.3364 |
| 4 | 564.3388 |
| 4 | 579.3297 |
| 4 | 587.3331 |
| 4 | 587.3356 |
| 4 | 606.3801 |
| 4 | 631.3085 |
| 4 | 632.3121 |
| 4 | 645.324 |
| 4 | 660.3473 |
| 4 | 671.4643 |

| | |
|---|---|
| 4 | 672.4723 |
| 4 | 673.4733 |
| 4 | 674.3615 |
| 4 | 696.4735 |
| 4 | 697.4778 |
| 4 | 704.4593 |
| 4 | 709.4851 |
| 4 | 714.5128 |
| 4 | 727.4592 |
| 4 | 761.4985 |
| 4 | 762.5014 |
| 4 | 803.4403 |
| 4 | 804.4504 |
| 4 | 804.5716 |
| 4 | 817.4548 |
| 3 | 110.9764 |
| 3 | 111.0094 |
| 3 | 124.0079 |
| 3 | 127.0771 |
| 3 | 128.0354 |
| 3 | 129.0196 |
| 3 | 130.0228 |
| 3 | 131.0237 |
| 3 | 133.0145 |
| 3 | 147.03 |
| 3 | 154.0626 |
| 3 | 173.0094 |
| 3 | 173.0824 |
| 3 | 174.0135 |
| 3 | 175.0254 |
| 3 | 187.0985 |
| 3 | 191.0204 |
| 3 | 195.0517 |
| 3 | 205.0532 |
| 3 | 216.0368 |
| 3 | 241.2175 |
| 3 | 253.2185 |
| 3 | 258.9827 |
| 3 | 260.0468 |
| 3 | 260.9803 |
| 3 | 262.0622 |
| 3 | 263.0595 |
| 3 | 265.1491 |
| 3 | 266.151 |
