## Supplemental Images for "Computational method for mapping mass signatures along developmental gradients reveals a novel role for a monosaccharide tetrose in maize salt-stress response"

### Supplemental Figures

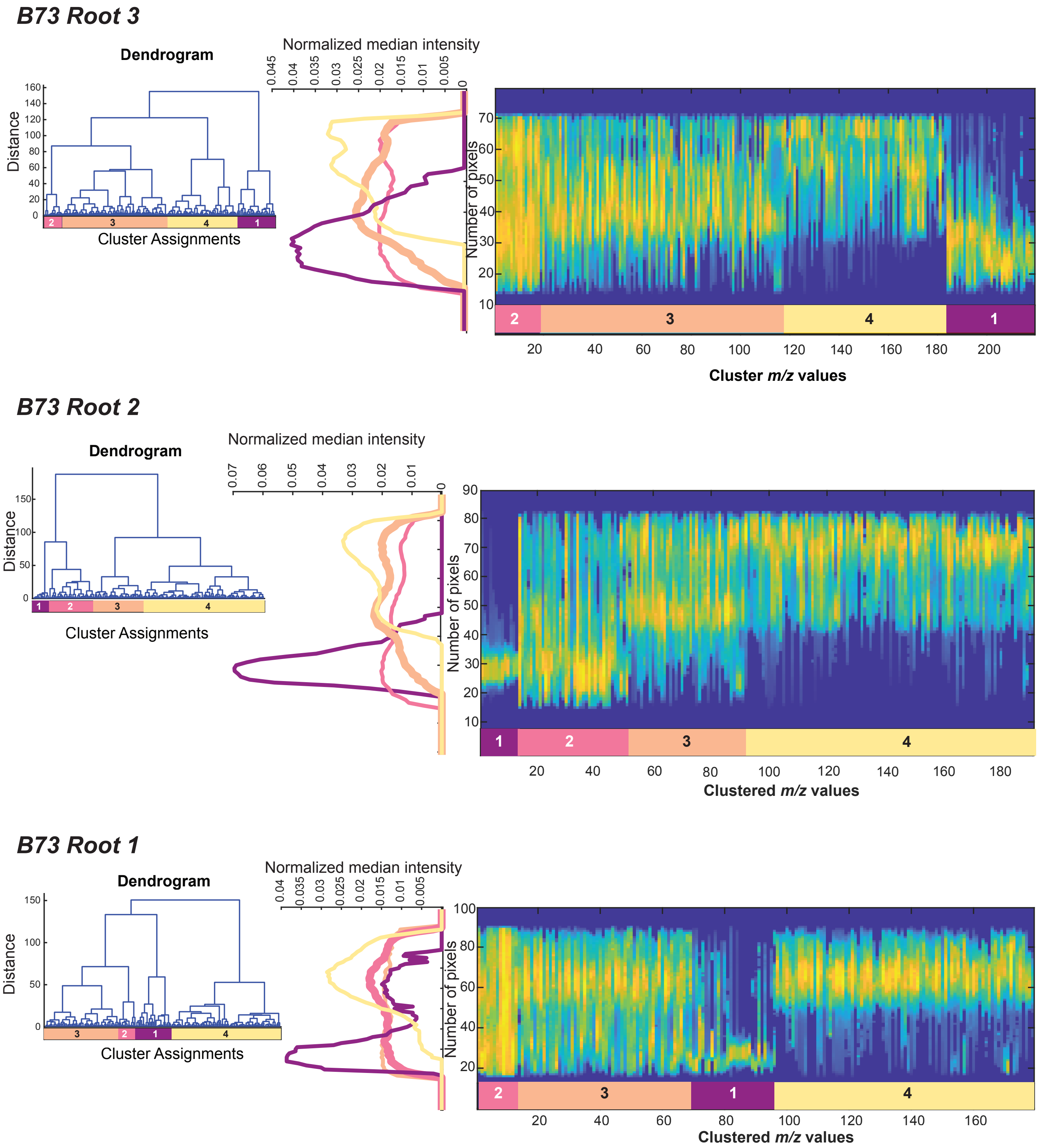


**Supplemental Figure 1. Dendrograms and linescan graphs for three B73 root replicates.** Dendrograms use Ward's method clustering to identify four main clusters. Linescan graphs showing the intensity profiles of mass signatures in each cluster. The y-axis is the position along the root axis where the root tip is at the origin. The x-axis corresponds to the clustered *m/z* linescans arranged smallest to largest within each cluster. The intensity of each linescan is normalized to a maximum value of 1. The intensity plots next to the linescan show the normalized median linescan intensity for each cluster along the root axis.

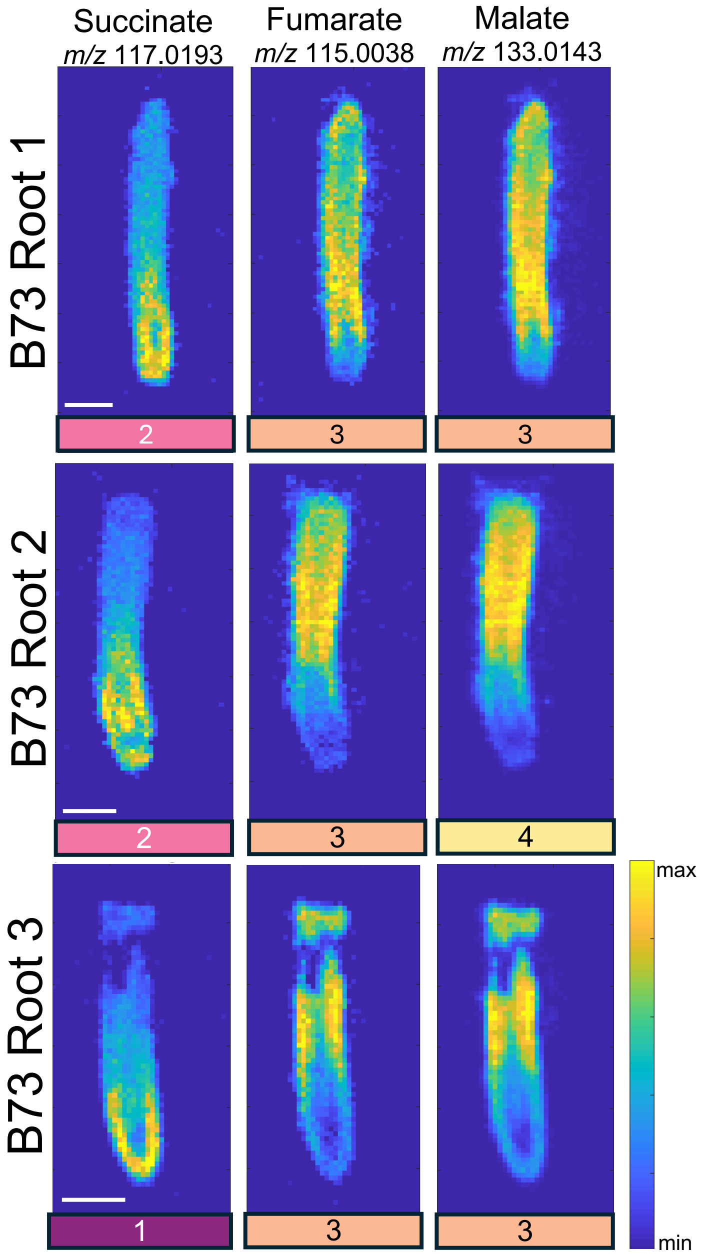


**Supplemental Figure 2.** TCA metabolites, succinate, fumarate and malate. MSI patterns from three B73 roots are shown. TIC-normalized with MSiReader +/- 5 ppm. Scale bar = 1 mm.

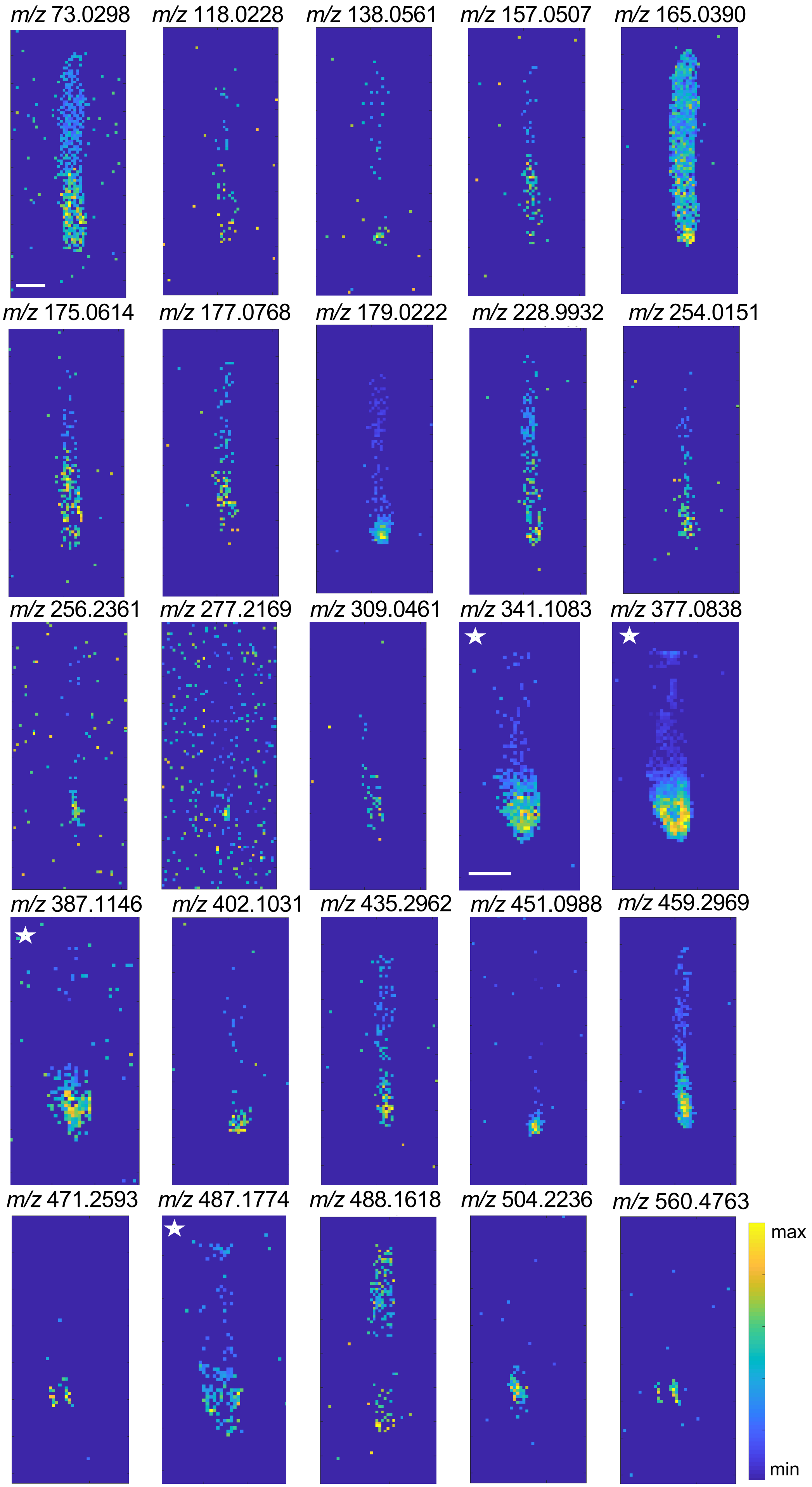


**Supplemental Figure 3.** New mass signatures identified with DIMPLE Cluster 1 characterized by meristem localization. MSI patterns from B73 root 1 (unstarred) and B73 root 3 (starred) are shown. TIC normalized in MSiReader, +/- 5ppm.

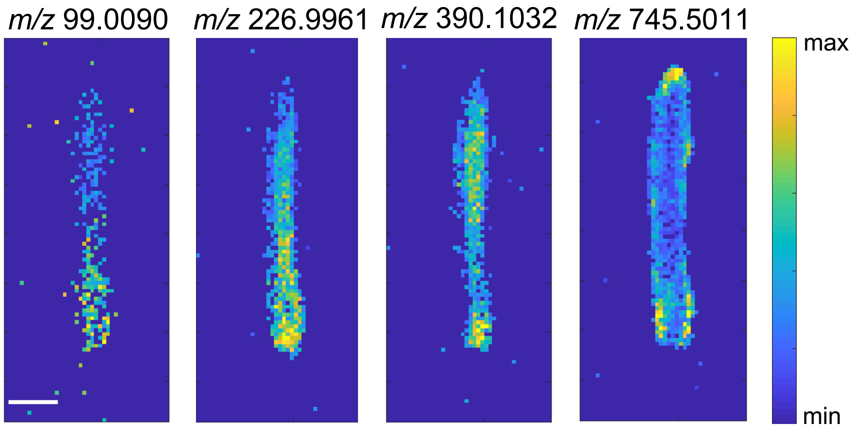


**Supplemental Figure 4.** Mass signatures with meristem enrichment identified in Zhang et al. 2023 that didn't localize to DIMPLE Cluster 1. MSI patterns from B73 root 1 shown. TIC normalized in MSiReader +/- 5ppm. Scale bar = 1 mm.

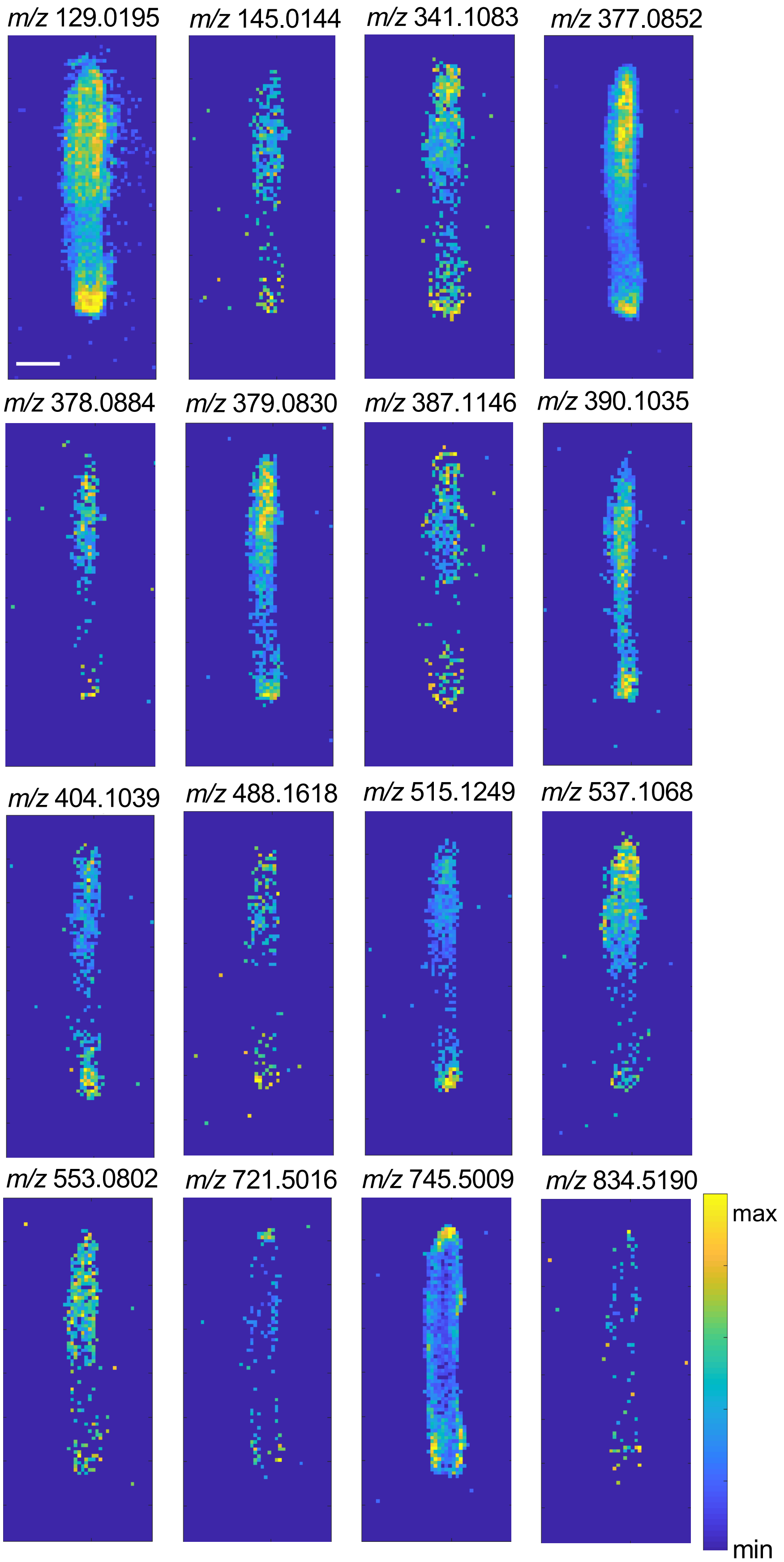


**Supplemental Figure 5.** Bimodal distribution patterns observed in the DIMPLE GUI. MSI patterns from B73 root 1 is shown. TIC normalized with MSiReader +/- 5ppm. Scale bar = 1mm

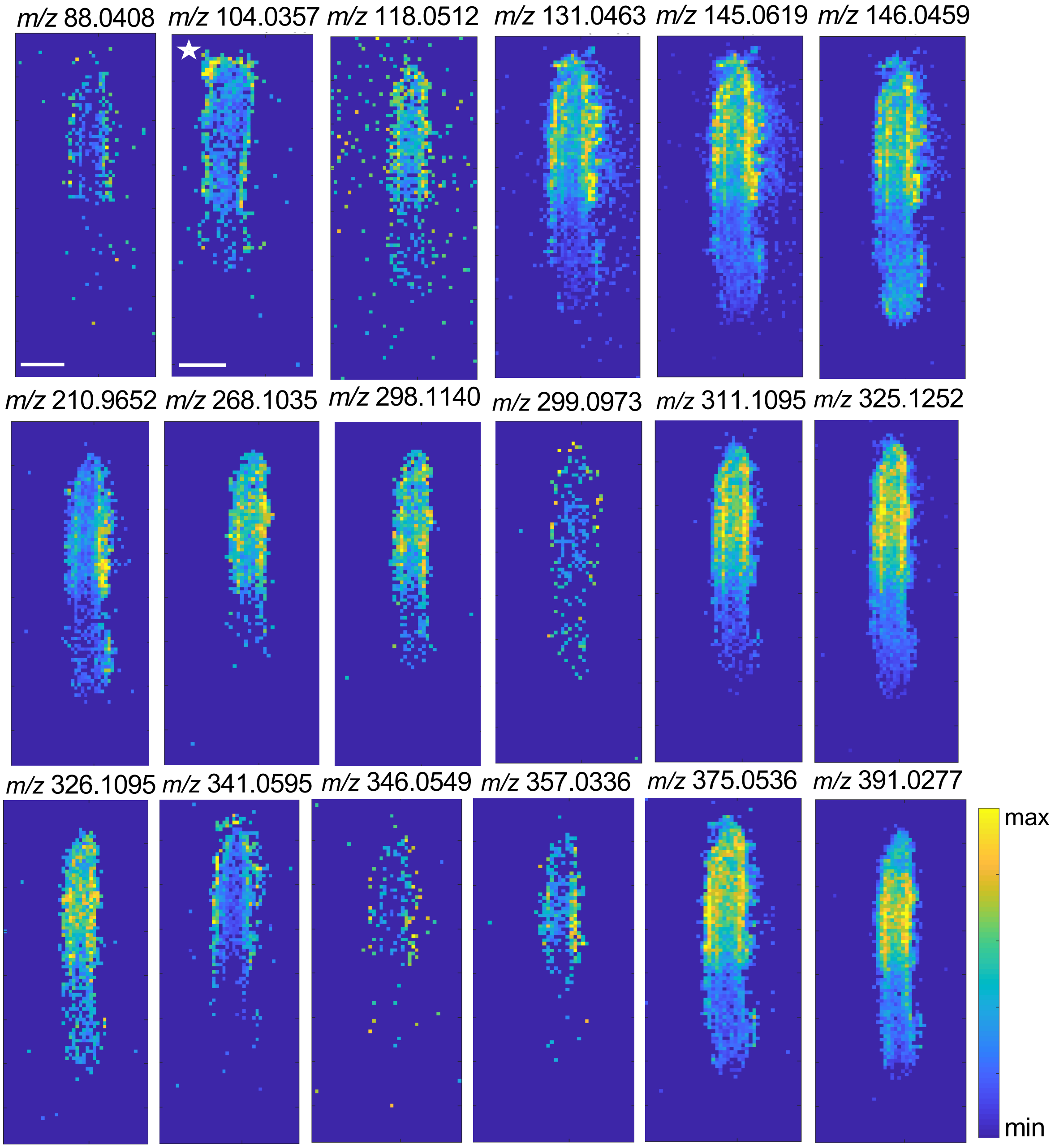


**Supplemental Figure 6.** Cortex localized mass signatures identified with the DIMPLE GUI, MSI patterns from B73 root 1 (unstarred) and B73 root 2 (starred) are shown. TIC normalized with MSiReader +/- 5ppm. Scale bar = 1mm.

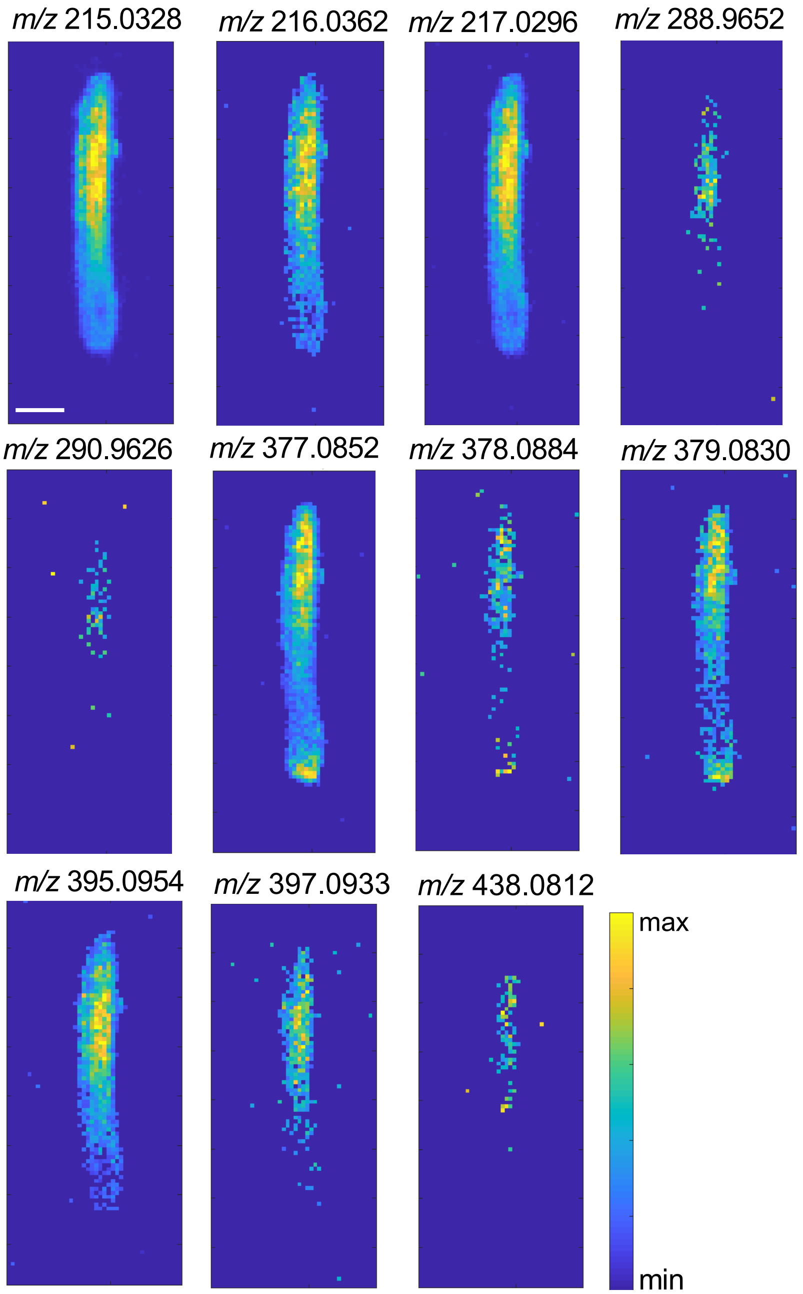


**Supplemental Figure 7.** Vasculature localized mass signatures identified with DIMPLE GUI. MSI patterns from B73 root 1 is shown. TIC normalized with MSiReader +/- 5ppm.

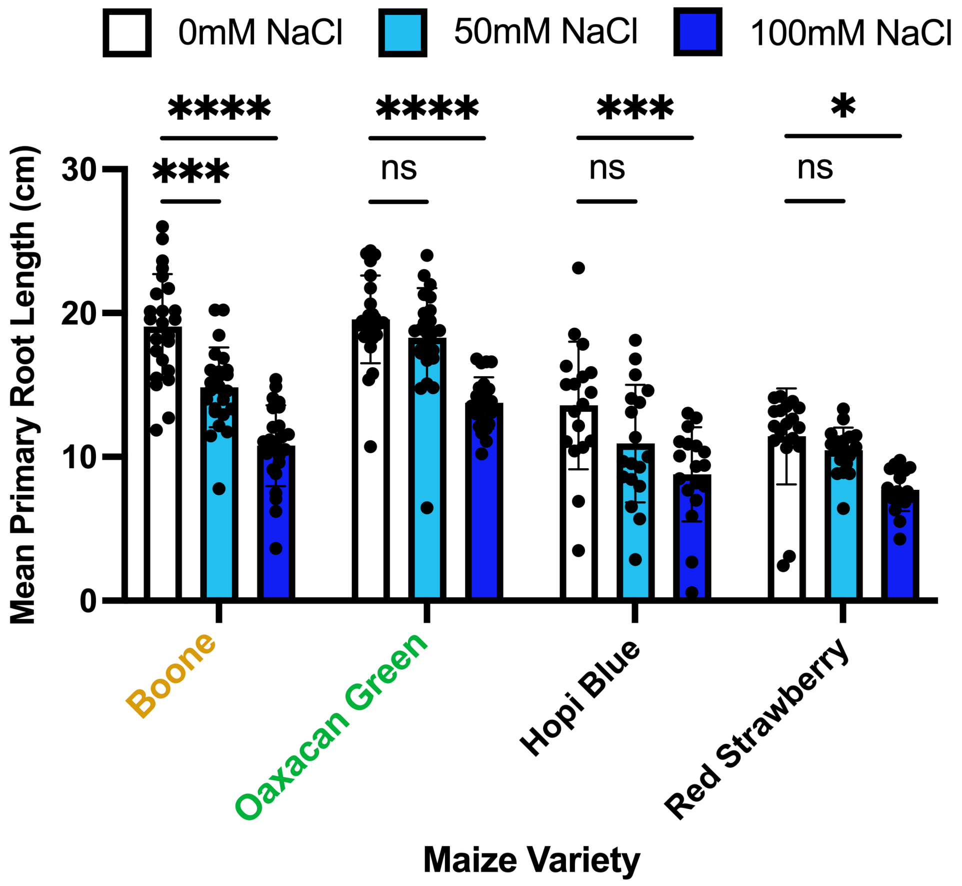


**Supplemental Figure 8.** Heirloom maize varieties, Oaxacan Green, Hopi Blue and Red Strawberry were more salt resilient compared to Boone County Dent maize. (50 mM and 100 mM NaCl). Maize was treated with either control, 50 mM NaCl or 100 mM NaCl conditions for 5days. The primary root length was measured for each condition. 2-way ANOVA, Tukey's multiple comparisions. Sample size for Boone n=24, Oaxacan Green n=24, Hopi Blue n=18, Red Strawberry n=18. P-value <0.0001 = ****, p-value < 0.001 = ***, p-value <0.05 =*

#### *Oaxacan Green Root 1*

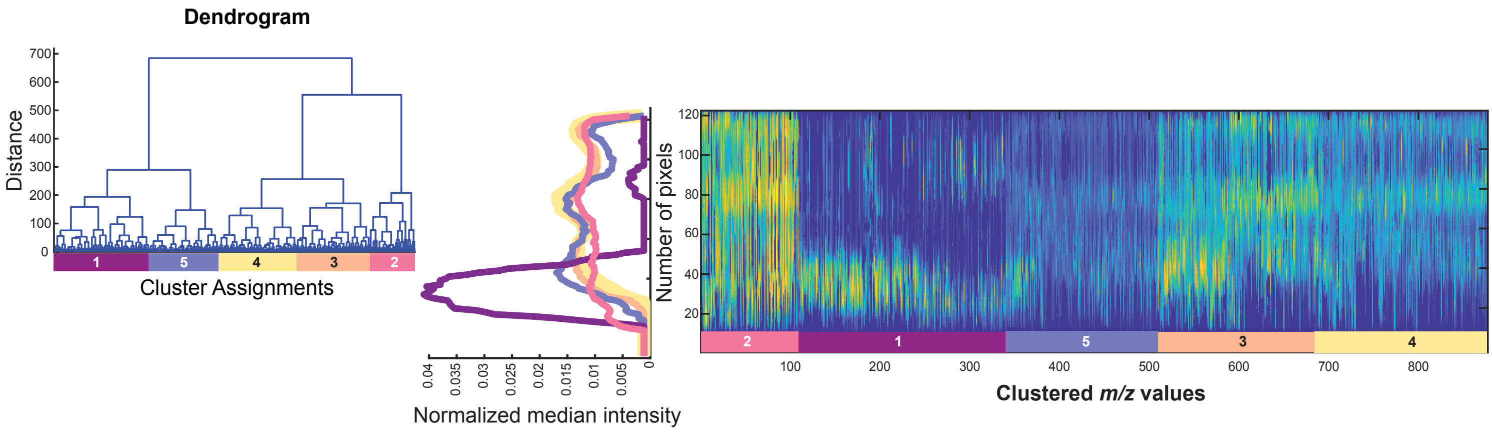


#### *Oaxacan Green Root 2*

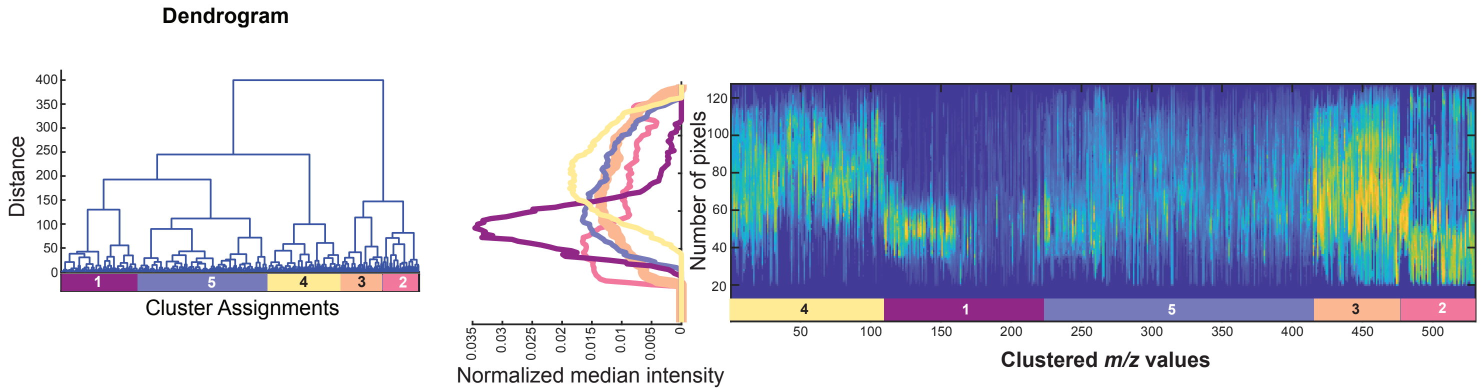


#### *Oaxacan Green Root 3*

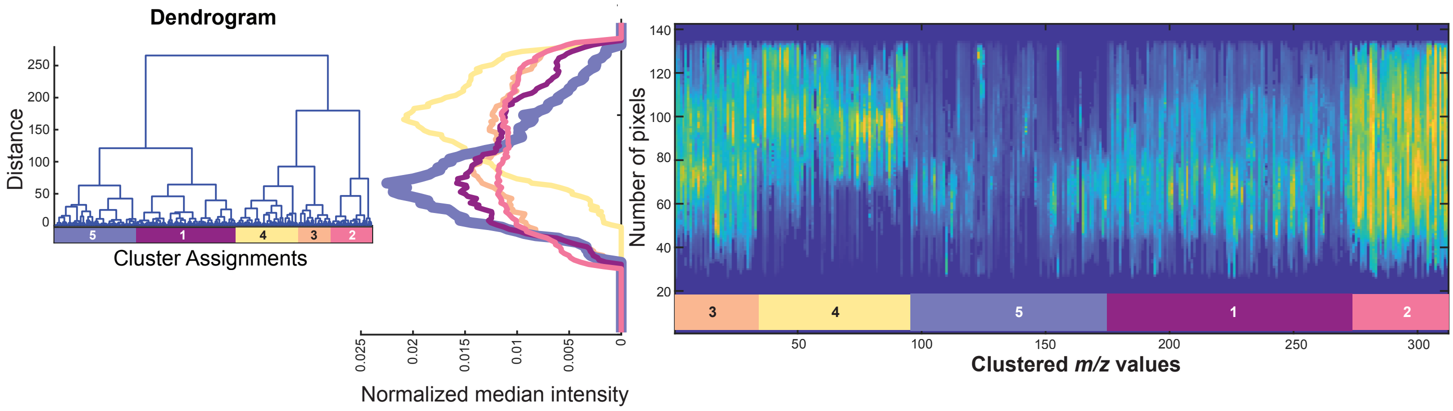


**Supplemental Figure 9. Dendrograms and linescans for three replicate Oaxacan Green roots.** Dendrograms use Ward's method clustering to identify five main clusters. Linescan graphs show the intensity profiles of mass signatures in each cluster. The y-axis is the position along the root axis where the root tip is at the origin. The x-axis corresponds to the clustered *m/z* linescans arranged smallest to largest within each cluster. The intensity of each linescan is normalized to a maximum value of 1. The intensity plots next to each linescan show the normalized median linescan intensity for each cluster.

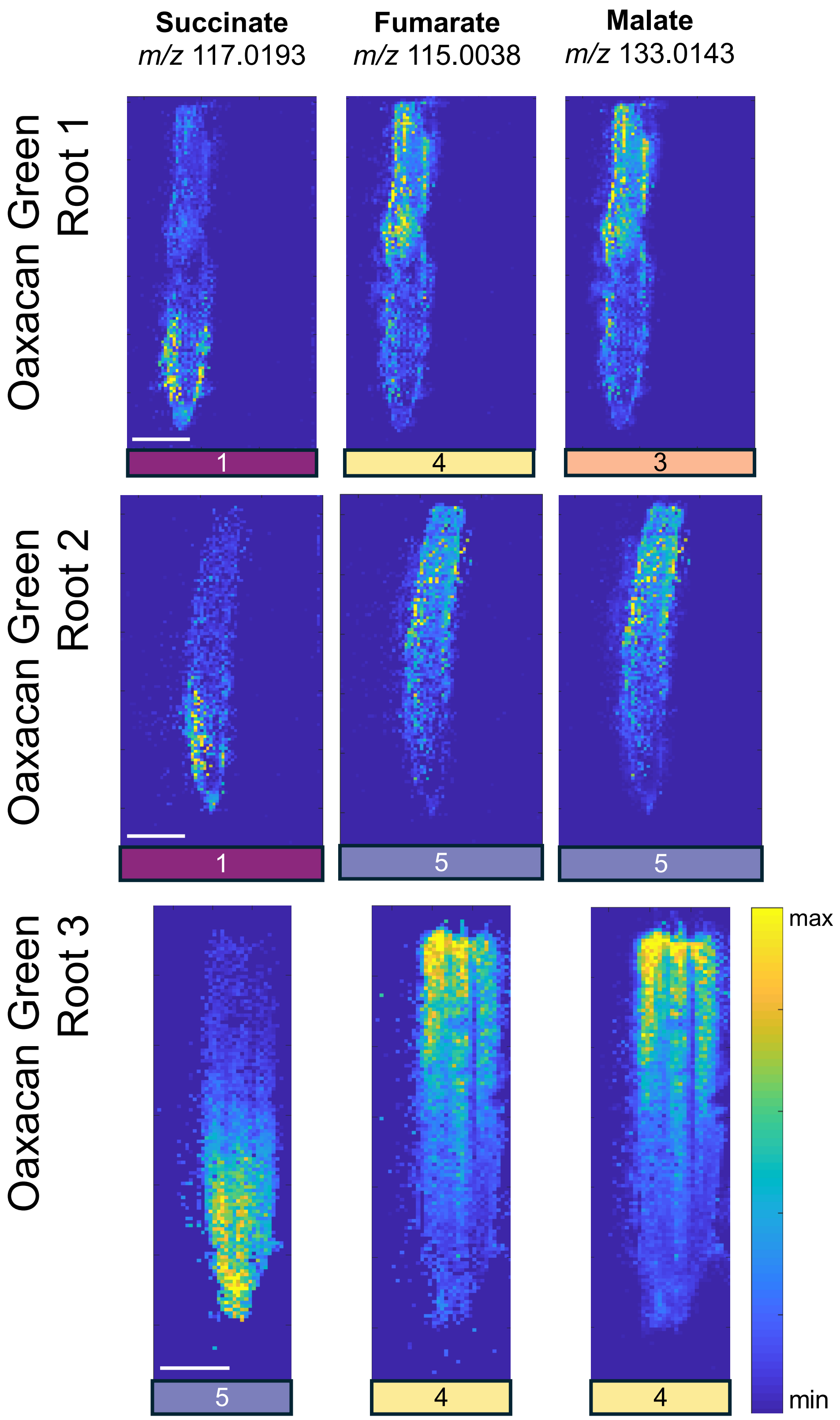


**Supplemental Figure 10.** TCA metabolites, succinate, fumarate and malate. MSI patterns from three Oaxacan Green roots. TIC-normalized metabolites with MSiReader, +/- 5ppm. Scale bar = 1mm.

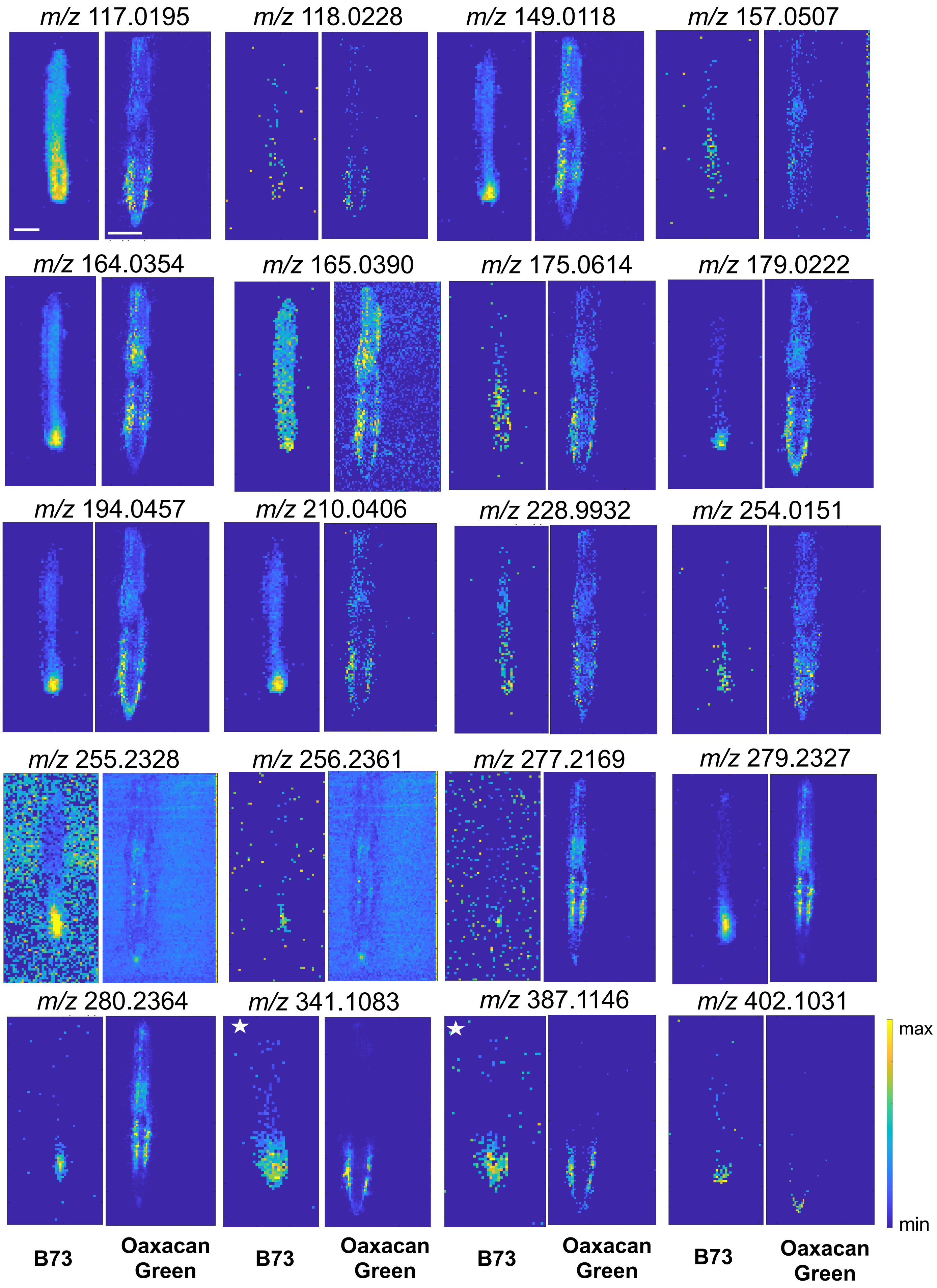


**Supplemental Figure 11.** Comparison of meristem enriched compounds identified in B73 and their localization in Oaxacan Green. 20 meristem enriched metabolites in B73 were present in the Oaxacan Green data, several show different localization patterns. MSI patterns from B73 root 1 (unstarred), and B73 root 3 (starred) and Oaxacan Green root 1 are shown. TIC normalized with MSiReader, +/- 5ppm. Scale bar = 1 mm.

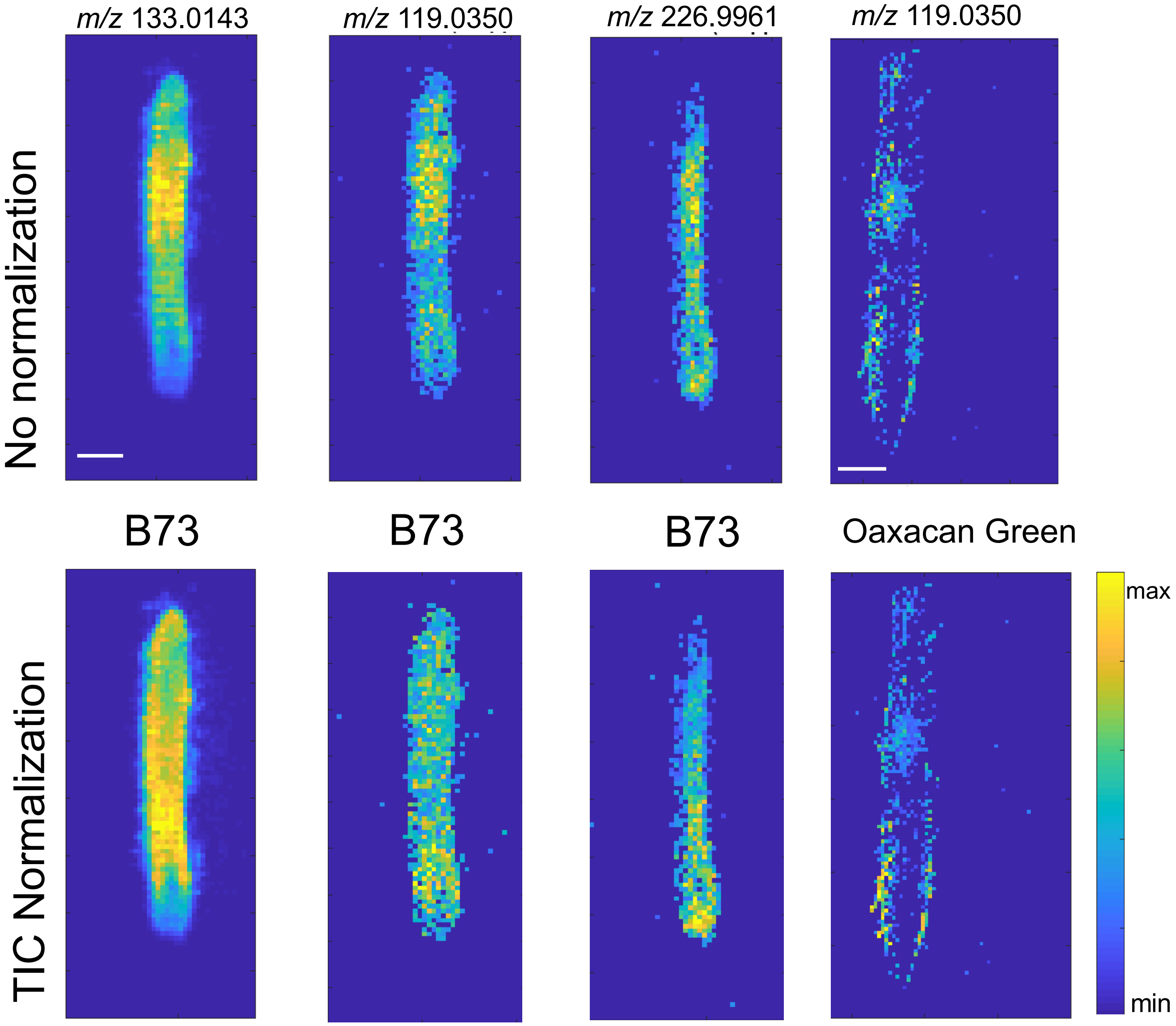


**Supplemental Figure 12.** Shifts in metabolite localization with TIC normalization. B73 and Oaxacan Green root metabolites with and without TIC normalization, MSI patterns from B73 root 1 and Oaxacan Green root 1 are shown. Generated in MSiReader +/- 5ppm.

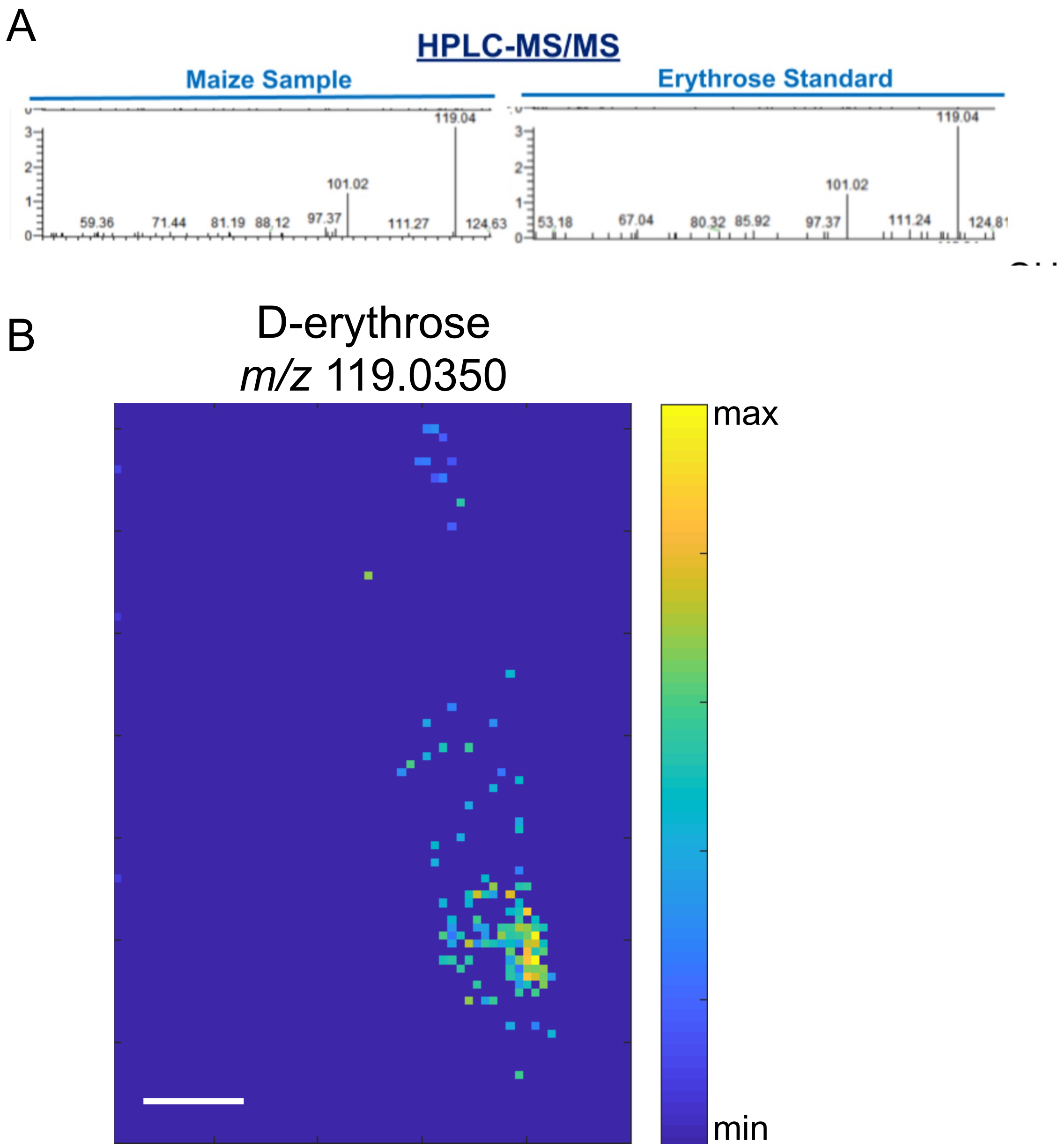


**Supplemental Figure 13.** A) HPLC-MS/MS fragmentation patterns for 119.0350 parent ion into m/z 101.02 in Boone County Dent maize roots and an erythrose standard (52 mM) visualized in Xcalibur. B) DESI-MSI of erythrose signature in a Boone root. TIC normalized with MSiReader +/- 5ppm.

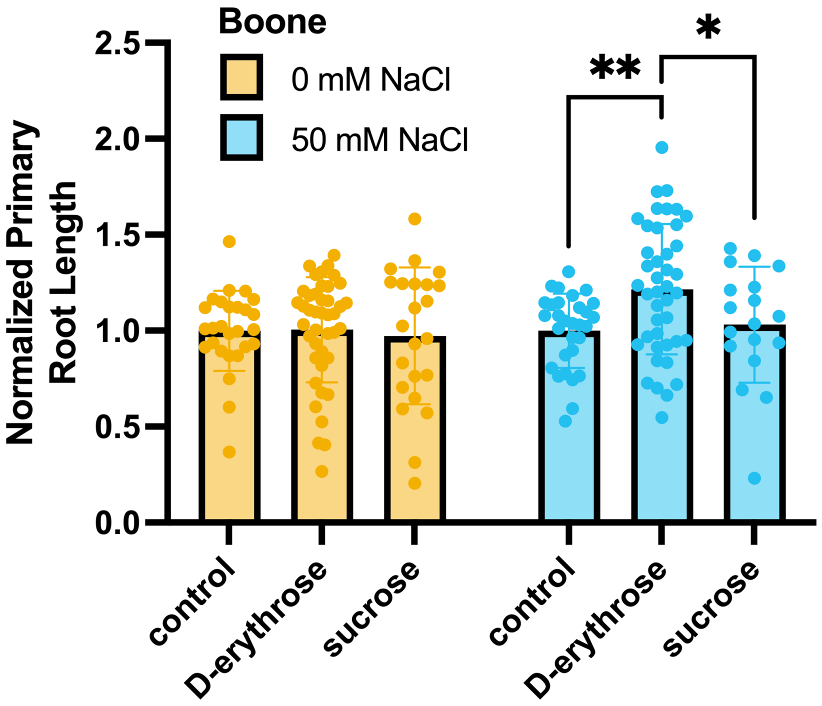


**Supplemental Figure 14.** Boone maize treated with salt and erythrose conditions. Normalized to control conditions for each salt treatment. Normalized Boone primary root length in control and 50 mM NaCl conditions with 200 μM D-erythrose or 200 μM sucrose treatment. Ordinary -way ANOVA Tukey's multiple comparisons test, p-value = 0.0019 = **, p-value = 0.0230 = *. Sample sizes for control conditions: control n = 27, D-erythrose n = 42, sucrose n = 23, NaCl conditions: control n = 30, D-erythrose n = 42, sucrose n = 18.
